# AlphaFold 3 captures oligomeric states and interaction dynamics of MLO ion channels

**DOI:** 10.64898/2026.04.10.716904

**Authors:** Jan W. Huebbers, Chandan K. Das, Alexander Speck, Myriam E. Fürst, Hanna Simon, Marie Laufens, Sophie C. J. Levecque, Matthias Freh, Maria Fyta, Ralph Panstruga

## Abstract

Mildew resistance Locus O (MLO) proteins have been originally identified as susceptibility factors for the fungal powdery mildew disease. Beyond immunity, they function in polarized secretion, including root and root hair elongation, trichome development, and fertilization. Moreover, MLO proteins mediate Ca²⁺ influx, either indirectly by recruiting Ca²⁺-permeable channels to the plasma membrane or by acting as ion channels themselves. The latter raises the question of whether MLO proteins oligomerize to mediate ion transport across membranes. Here, we present an AlphaFold 3-based modeling pipeline for the reproducible assessment of MLO-containing protein complexes using AlphaFold’s built-in confidence metrics together with structural and dynamic analyses.

The resulting predictions for homo-oligomers of the prototypic barley Mlo support dimeric and trimeric assemblies, with the trimer forming a central membrane-spanning pore. Notably, AlphaFold 3 captured discrete conformational states of this trimer, as reflected by the clustering of confidence metrics. Computational structural analyses indicated that higher-confidence models adopt a closed pore conformation, whereas lower-confidence predictions reflect progressively expanding pore diameters. Molecular dynamics simulations further showed Ca²⁺ permeability of the putative open models. Our pipeline similarly predicts trimeric assemblies for MLO variants from *Arabidopsis thaliana* and *Marchantia polymorpha*, suggesting a conserved MLO structural scaffold within the land plant lineage. Additional Molecular Dynamics simulations revealed that closed models of barley Mlo and *A. thaliana* MLO2 open under simulated membrane tension, supporting the notion that MLO proteins are mechanosensitive ion channels. Moreover, predictions of MLO proteins with its known interactors, EF-hand proteins and exocyst complex subunit EXO70 proteins, suggest a mechanism for feedback inhibition of MLO-mediated ion flux and provide comprehensive experimental support for AlphaFold 3-predicted protein interfaces. Altogether, our results provide a structural framework for MLO channel architecture and regulation, while our prediction, modeling, and simulation pipeline should be useful beyond the study of this specific protein family.

**One-sentence summary:** This article describes AlphaFold 3-based analyses of MLO proteins, revealing the predicted structure of MLO membrane pores, their dynamic opening and closing, and their association with interacting proteins, including calmodulin and calmodulin-like calcium sensor proteins and exocyst complex subunit EXO70 proteins.

## Introduction

Mildew resistance locus O (MLO) proteins are integral membrane proteins found exclusively in plants. They were first identified in barley (*Hordeum vulgare*) as susceptibility factors for the powdery mildew disease (Jørgensen, 1971). To date, MLO-dependent powdery mildew susceptibility has been reported in numerous monocot and eudicot species (Consonni *et al*., 2006; Kusch and Panstruga, 2017). MLO proteins are commonly grouped into seven phylogenetic clades (Kusch *et al*., 2016) and, beyond immunity, they are linked to processes that involve cell polarity and polarized secretion, including root thigmomorphogenesis (clade I; Chen *et al*., 2009), root hair growth (clade II; Ogawa *et al*., 2025), pollen tube growth and reception (clade III; Ju *et al*., 2021; Meng *et al*., 2020), as well as callose deposition and trichome development (clades V and VI; Kusch *et al*., 2019; Huebbers *et al*., 2024).

MLO proteins share a conserved topology of seven membrane-spanning domains, an extracellular/luminal N-terminus (NT), and an intracellular C-terminus (CT; Devoto *et al*., 1999). They have been associated with Ca^2+^ signaling, due to their conserved binding site for the calcium sensor calmodulin (CAM) in the proximal part of their C-terminus (Kim *et al*., 2002; Bongartz *et al*., 2023). Moreover, they either recruit Ca^2+^-permeable channels to the plasma membrane (Meng *et al*., 2020; Ogawa *et al*., 2025) and/or act as such channels themselves (Gao *et al*., 2022; Gao *et al*., 2023). In the latter scenario, oligomerization has been suggested as a mechanism to generate a membrane-spanning, ion-permeable pore (Li and Xiao, 2025). Consistent with this idea, Fluorescence/Förster Resonance Energy Transfer (FRET) and FRET-Fluorescence Lifetime Imaging (FRET-FLIM) experiments support MLO self-association (Elliott *et al*., 2005; Jones *et al*., 2017). Of course, these approaches do not resolve the oligomeric state (e.g., dimer, trimer, or tetramer) of MLO complexes or whether the resulting assemblies allow ion transition. Structural information would directly address these questions. However, structure determination for polytopic membrane proteins remains technically demanding (Carpenter *et al*., 2008; Moraes *et al*., 2014).

The advent of AlphaFold (AF) made deep learning-based, high-quality protein structure prediction accessible (Senior *et al*., 2020). This was recognized by half the Nobel Prize in Chemistry for AF2 (Jumper *et al*., 2021; Callaway, 2024) and is reflected in results from the Critical Assessment of Structure Prediction (CASP) community benchmark. During CASP16, AF3 (Abramson *et al*., 2024) predicted all monomer targets correctly and performed well on homo-oligomers and membrane proteins (Yuan *et al*., 2026). Especially for the latter, scientists have been skeptical due to the relative scarcity of training data for membrane proteins, which is itself a consequence of the aforementioned technical challenges in determining such structures. Nevertheless, AF2 already has predicted membrane proteins accurately and even captures alternative conformational states of membrane protein complexes, depending on the supplemented template structures (Ngo *et al*., 2025).

Molecular Dynamics (MD) simulations complement structural biology by embedding a static structure in an explicit physicochemical environment (lipids, water, and ions) and calculating molecular motion under a molecular mechanics force field. By numerically integrating Newton’s equations of motion, MD simulations sample thermally sensible conformations and provide an atomistic view of how proteins couple to, for example, bilayer properties such as thickness, curvature, and lateral pressure (Hollingsworth and Dror, 2018). This interplay between membrane, protein, ions, and solvent is particularly relevant for channels and transporters, whose function can depend on pore hydration/dehydration, transient side-chain rearrangements, and subtle helix motions that may not be apparent from a single static model (Aryal *et al*., 2015; Aryal *et al*., 2017). For ion channels, MD is used to probe permeation pathways, including transient coordination sites that shape selectivity (Acharya *et al*., 2025; Gabriel *et al*., 2021; Schackert *et al*., 2023).

Here, we used an AF3-based pipeline to predict the oligomeric state of *Hv*Mlo, which supported a trimeric arrangement of *Hv*Mlo protomers into a membrane-spanning pore. We carried out all-atom MD simulations of the predicted trimers inserted in a membrane model to assess Ca²⁺ ion transitions and to test whether lateral tension promotes pore opening. We further leveraged our AF3 pipeline to explore whether MLO proteins from *A. thaliana* and *Marchantia polymorpha* likewise assemble into homo-trimers and how MLO proteins engage known interaction partners, including *At*CAM2, *At*CML12 and EXO70 proteins.

## Results

### AF3 simulations suggest that *Hv*Mlo forms trimeric membrane channels

We created a reproducible AF3-based pipeline to probe structures of protein complexes (Figure 1A). Our pipeline enables the swift submission of multiple simulation jobs to the AF3 server *via* pre-compiled JSON input files and the subsequent automated extraction and processing of the AF3 server’s output using structure visualization, the extraction of confidence metrics, and the visualization of these metrics. For reproducibility, all our AF3 server runs were fixed using twelve different seeds. As the AF3 server provides five models per seed, we retrieved 60 models in total per input condition.

**Figure 1:**
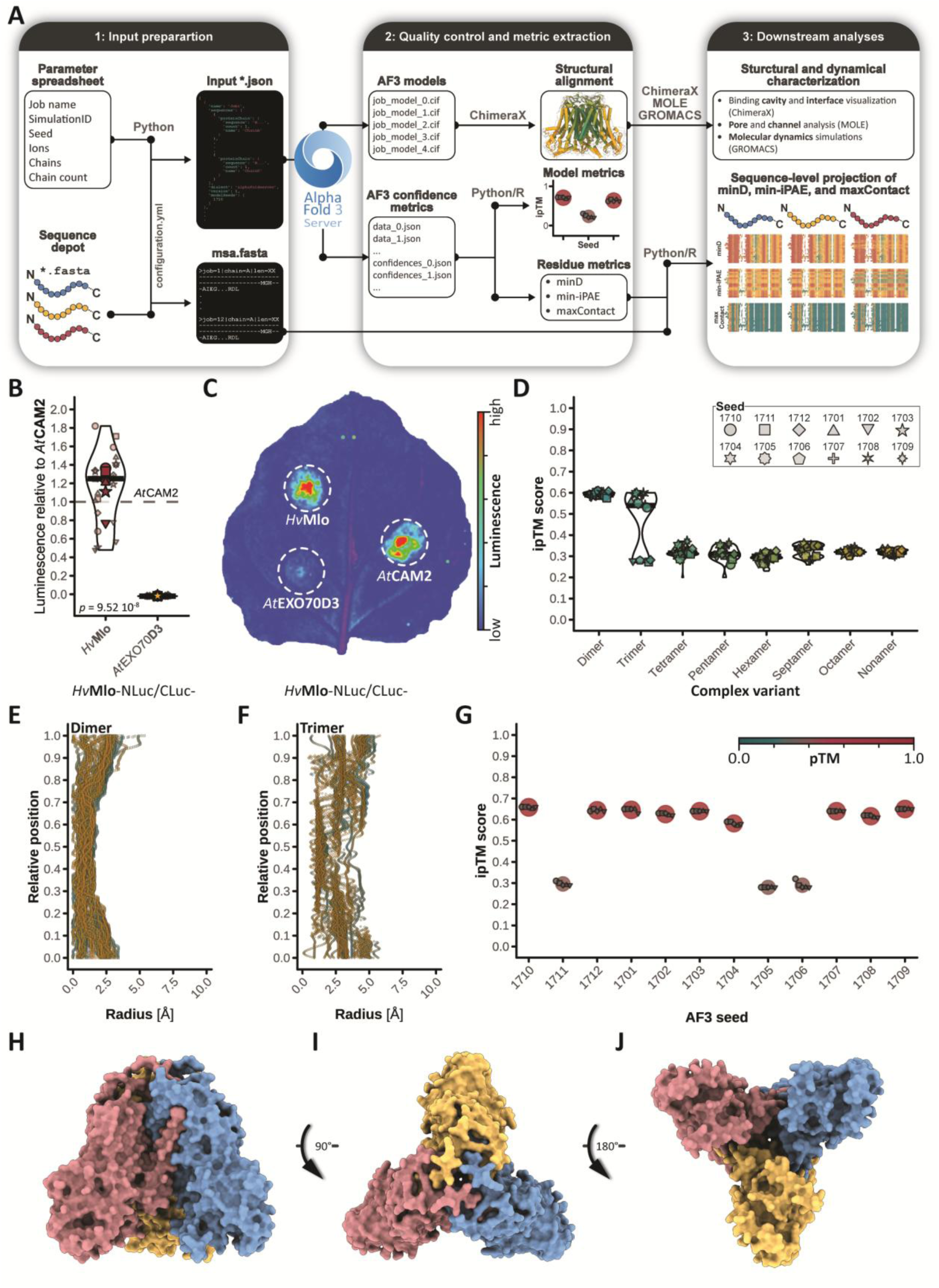
AlphaFold 3 predictions suggest that *Hv*Mlo proteins form homo-trimers. **A)** Overview of our AF3-based modeling and analysis pipeline. **1:** AF3 batch jobs are submitted as *.json files generated from a configuration file, a parameter spreadsheet, and a *.fasta sequence repository using *Python* scripts that are executed *via* the command line. **2:** AF3 outputs are downloaded from the server and assessed by structural alignment in ChimeraX and inspection of pTM and ipTM confidence metrics. Per-residue confidence matrices are extracted, transformed, and exported as *.csv files. **3:** AF3 models are inspected in ChimeraX, analyzed with MOLE for pore characterization, and subjected to molecular dynamics simulations in GROMACS. Per-residue confidence metrics are integrated with multiple-sequence alignments (MSAs) of input chains to identify interface-relevant residues. **B, C)** Luciferase complementation imaging of *HvMlo-NLuc* and *HvMlo-CLuc* coexpressed in *N. benthamiana* leaves. CLuc-*At*CAM2 served as a positive control and CLuc-*At*EXO70D3 as a negative control (see Huebbers *et al*., 2024). **B)** Violin plots show the distribution of relative luminescence signals from six independent experiments (large points), with four replicates (small points) each. Statistical significance was determined by Student’s *t*-test. **C)** Pseudo-colored leave representing a single replicate. **D)** AF3 ipTM score distributions for AF3 predictions of *Hv*Mlo homo-oligomers. Violin plots summarize scores across twelve prediction runs (one per seed, as indicated), with five models per run. **E, F)** Pore-radius profiles for *Hv*Mlo dimer models (**E**; 60 models) and higher-confidence *Hv*Mlo trimer models (**F**; 40 models), obtained via the MOLE API. The color grading indicates different seeds. **G)** pTM and ipTM confidence scores for *Hv*MloΔIDR homo-trimers. Large points indicate the mean ipTM per seed, and small points indicate ipTM values for individual models. Colors correspond to pTM scores as indicated. **H–J)** Representative AF3 prediction of the *Hv*MloΔIDR homo-trimer (seed 1710, model 0). Subunits are color-coded: A, blue; B, yellow; C, red. Perspectives show a side view (subunit B in the back), the extracellular/luminal (top) view, and the cytosolic (bottom) view.

FRET and FRET-FLIM experiments indicated that MLO proteins homo-oligomerize (Elliott *et al*., 2005; Jones *et al*., 2017). We used luciferase complementation imaging (LCI) in *Nicotiana benthamiana* leaves to corroborate these findings and observed strong relative luminescence (mean ± SD = 1.15 ± 0.38) upon coexpression of *HvMlo-NLuc* with *HvMlo-CLuc* (Figure 1B, C). Based on this *in-planta* experimental data, we used our modeling pipeline to assess the oligomerization behavior of *Hv*Mlo (UniProt: P93766). We submitted up to nine *Hv*Mlo chains to the AF3 server. AF3 reports an interface predicted template modeling score (ipTM) that captures confidence in the relative positioning of subunits within a complex (Abramson *et al*., 2024) and provides a useful proxy for whether AF3 supports a given protein complex. Dimer predictions consistently reached ipTM values of ∼0.60, whereas trimer predictions split into higher-confidence (ipTM ∼ 0.55) and lower-confidence (ipTM ∼ 0.25) models associated with specific seeds (Figure 1D; Supplemental File 1). Visual inspection further revealed that for assemblies with n > 4 (i.e., tetramer to nonamer), the predicted subunit arrangements were either incompatible with the experimentally validated membrane topology of *Hv*Mlo (Supplemental Figure 2A, C, D; Devoto *et al*., 1999) or appeared distorted, resembling, for instance, side-by-side packing of lower-order complexes (Supplemental Figure 1D, E; Supplemental Figure 2B).

Given their comparably high ipTM scores and plausible architectures (Supplemental Figure 1B, C), we focused subsequent analyses on the *Hv*Mlo dimer and trimer. Following the hypothesis that *Hv*Mlo oligomers form ion-permeable pores in membranes (Aryal *et al*., 2017; Gao *et al*., 2022; Gao *et al*., 2023; Li and Xiao, 2025), we used MOLE (Raček *et al*., 2025) to probe all predicted *Hv*Mlo dimers and the higher-confidence *Hv*Mlo trimer models for intra-complex pore pathways (Figure 1E, F). Consistent with the visual inspection of the models (Supplemental Figure 1C), the profiles indicate a marked yet narrow pore in the *Hv*Mlo trimers with a minimal diameter of ∼1.0 Å and a maximum diameter of ∼5.0 Å. By contrast, comparable pore pathways were not detected in the dimers. Given that the trimer is the highest-order *Hv*Mlo assembly among the AF3-supported stoichiometries that remains structurally plausible and that it forms a central pore, we consider the trimer the physiologically relevant candidate, although multiple quaternary states may coexist (Lin *et al*., 2026; Ferreira and Felice, 2001).

The predicted *Hv*Mlo trimers (Supplemental Figure 1C), are threefold rotationally symmetric (C3) complexes that have a central membrane-spanning pore (Figure 1H–J). AF3 per-residue confidence metrics support the characterization of interfaces between subunits in protein complexes (subunits A–C). These metrics include distance, individual predicted aligned error (iPAE), and contact scores. Briefly, the distance score reports predicted residue–residue distances, the iPAE score reflects confidence in the relative positioning of residues, and the contact score summarizes these features as the probability that two residues are in contact. To identify interface-relevant residues, we compute minimal values for distance and iPAE scores (minD, min-iPAE), whereas for contact scores we calculate maximal values (maxContact), thereby collapsing each residue-by-residue matrix to a per-residue vector of summary statistics. For the *Hv*Mlo trimer, these adjusted per-residue metrics indicate that transmembrane domains (TMs) TM1–TM4, and TM6, extracellular/luminal loops (ECs) EC1–EC3, as well as the extracellular/luminal NTs contribute to trimer interfaces (Supplemental Figure 3A; Supplemental File 1). This pattern is also supported by direct visualization of interface residues in subunit A and its corresponding residues within a 4 Å-radius in subunits B and C (Supplemental Figure 3B). By contrast, intracellular loops (ICs) and the intracellular CTs do not contribute to complex formation across the eight higher-confidence predictions (Supplemental Figure 3A).

In line with these observations, the intrinsically disordered region (IDR; Kusch *et al*., 2016) in the distal CT did not show interface participation. In AF3, residues in this region were positioned with comparatively low confidence, as reflected by reduced per-residue confidence in their inter-chain relationships (Supplemental Figure 3A, min-iPAE). Because non-interfacial IDRs depress ipTM scores (Dunbrack, 2025), we removed the IDR in the distal CT (residues 434–533) and repeated trimer predictions. These *Hv*MloΔIDR trimers again exhibited a bimodal distribution of ipTM scores (Figure 1G; Supplemental File 2), while IDR removal increased ipTM scores by ∼0.10. For *Hv*MloΔIDR trimers, AF3 pTM scores, reflect confidence in the overall fold, were positively correlated with ipTM scores and displayed the same seed-dependent bimodal distribution (Figure 1G). Altogether, our data indicate that AF3 confidently predicts *Hv*MloΔIDR homo-trimers that adopt the same C3-symmetric architecture as trimers formed by full-length *Hv*Mlo subunits.

### AF3 captures dynamic states of the *Hv*Mlo trimer pore

Structural superpositions of the five models returned per AF3 run (e.g., output per seed) showed closer agreement among higher-confidence predictions than among lower-confidence predictions. This difference is captured by mapping the per-residue root mean square deviation (RMSD; relative to model 0) onto a higher-confidence run (Figure 2A) and a lower-confidence model (Figure 2B). We also noticed that lower-confidence models form a wider central pore than higher-confidence models (Figure 2A, B, top view; Supplemental Movie 1). Therefore, we submitted all *Hv*MloΔIDR trimer models to MOLE for pore characterization. Pore profiles from higher-confidence predictions (Figure 2C) showed a minimal radius of about 1.0 Å, whereas lower-confidence models exhibited larger minimal radii of ∼3.0 Å (Figure 2D). We selected model 0 of seed 1710 (Figure 2C) and model 2 of seed 1705 (Figure 2D) as representative higher- and lower-confidence structures for three-dimensional pore visualization in ChimeraX. In both models, the narrowest constriction is formed by L72 and I75 (Figure 2E, F), while, in 1705_02, the pore remains open to a radius of ∼3.0 Å at this position (Figure 2F).

**Figure 2:**
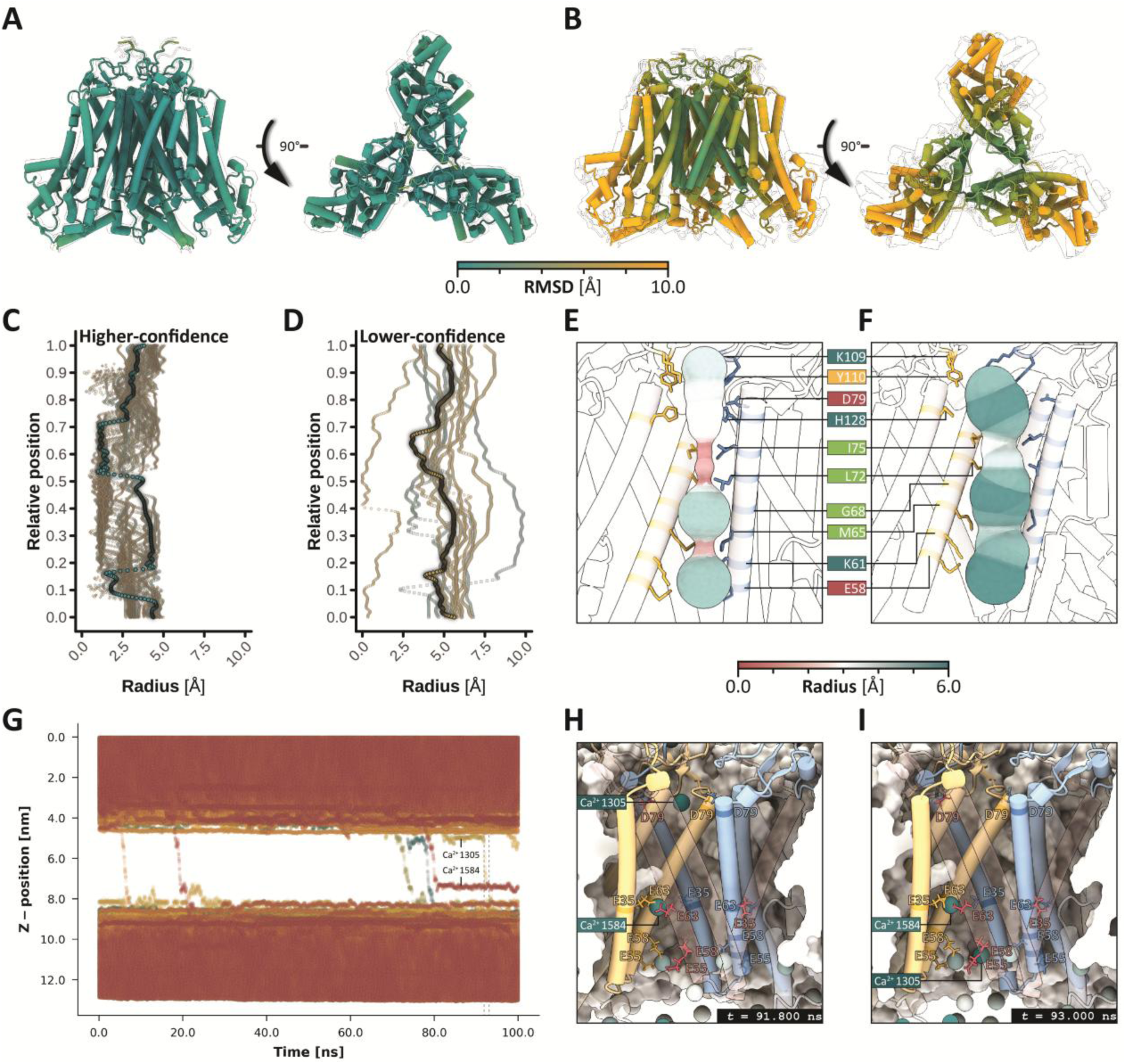
AF3 predictions capture different conformations of the *Hv*Mlo trimer pore. **A, B)** Structural superposition of *Hv*MloΔIDR trimer models from a higher-confidence (seed 1710; **A**) and a lower-confidence AF3 run (seed 1705; **B**). The five models per seed were aligned and superposed in ChimeraX. Per-residue RMSD was mapped onto model 0, while models 1–4 are displayed as transparent overlays. Perspectives (left to right): side view (subunit C in the back) and extracellular/luminal (top) view. **C, D)** Pore-radius profiles of *Hv*MloΔIDR models from higher-confidence (**C**) and lower-confidence (**D**) AF3 runs. Pore descriptors were obtained *via* the MOLE API. Point colors indicate different AF3 seeds. Profiles of seed 1710, model 0 (**C**) and seed 1705, model 2 (**D**) are highlighted and correspond to the three-dimensional pore representations in **E** (1710_00) and **F** (1705_02). **E, F)** Three-dimensional “balloon” representation of transmembrane pores, obtained *via* the MOLE API. To improve pore visibility, residues 1–50 of subunit B (yellow) and subunit C are omitted. Pore-lining residues were identified in ChimeraX (*interface select*) for the higher-confidence model (**E**) and are highlighted in subunits A and B. **G)** Ca²⁺ trajectories along the membrane normal (z-axis) in an all-atom MD simulation initiated from the lower-confidence model 1705_02 (**F**). Dashed lines indicate the time points shown in **H** and **I** (93.000 ns). **H, I)** Three-dimensional models for the two simulation frames indicated in **G.** Non-transparent cartoons of residues 1–160 of subunits A (blue) and B (yellow) are shown. Side chains of pore-lining acidic residues are displayed as sticks for all subunits (with subunit C in front) and labeled using the one-letter amino acid code. The membrane is shown in cross-section.

We carried out all-atom MD simulations for the lower-confidence *Hv*MloΔIDR model 1705_02 in a sphingomyelin membrane to test whether the putatively open pore allows ion permeation. Over 100 ns, we observed six Ca²⁺ translocation events into the cytosol (Figure 2G), suggesting that the pore formed by the 1705_02 *Hv*MloΔIDR trimer allows Ca²⁺ permeation. In addition, at least 14 Cl⁻ ions traversed from the intracellular to the extracellular side (Supplemental Figure 4A).

Whereas Cl⁻ ions passed the pore with little apparent hindrance (Supplemental Figure 4B, C), Ca²⁺ ions formed transient coordination complexes with acidic side chains (Figure 2H, I; Supplemental Movie 2). Of these, the pore-lining residues E35, E55, and E58 form a prominent ring-like arrangement on the cytosolic mouth (Supplemental Figure 4D, E). Similar arrangements of acidic residues shape selectivity for divalent over mono- or trivalent cations in other channels, such as mammalian TRPV6 (Yelshanskaya *et al*., 2021).

### The architecture of MLO trimers is conserved within the land plant lineage

We applied our AF3 pipeline to ΔIDR variants of the 15 *A. thaliana* and the four *M. polymorpha* MLO proteins to assess whether a trimeric scaffold is generally plausible across MLO clades (Figure 3A). As for *Hv*Mlo, pTM and ipTM scores across all tested MLOΔIDR proteins displayed bimodal distributions (Figure 3B, C; Supplemental File 4). Notably, clade III MLO proteins consistently exhibited a larger fraction of seeds associated with lower-confidence models than in other clades (Figure 3B). Visual inspection revealed that, for the phylogenetically closely related *At*MLO7, -8, and -10 (Figure 3A), the higher-confidence AF3 predictions included an additional, eighth transmembrane helix formed by the extended NT of these isoforms (Supplemental Figure 5A–C).

**Figure 3:**
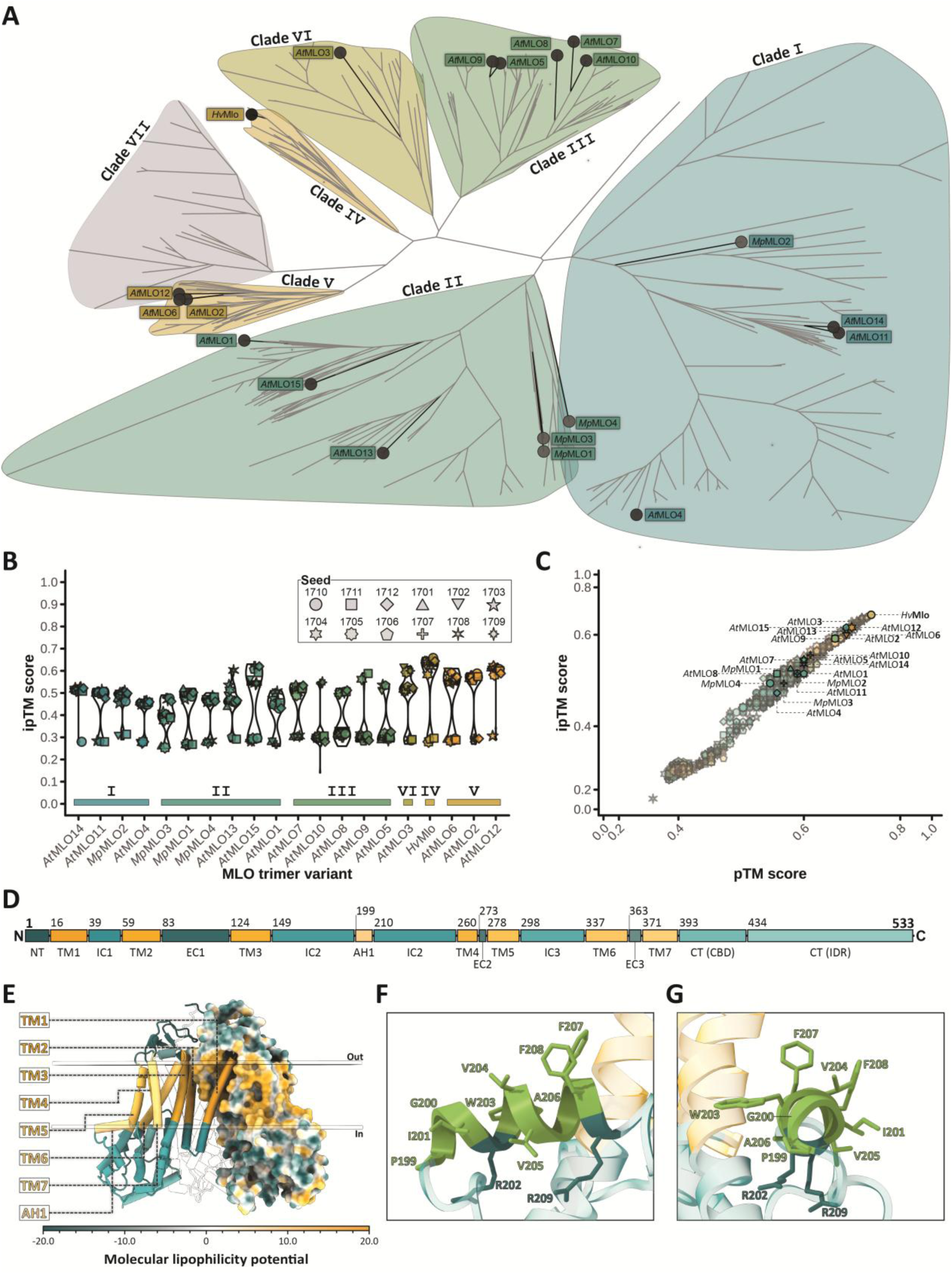
Land-plant MLO proteins are predicted to form trimers and share a conserved membrane topology. **A)** Phylogenetic tree inferred from 351 MLO protein sequences. MLO proteins analyzed in this study are labeled. **B)** Distribution of ipTM scores for AF3 predictions of *A. thaliana* and *M. polymorpha* MLO trimers and the prototypic *Hv*Mlo, modeled without their C-terminal IDRs. Proteins are ordered according to the phylogeny shown in **A**, and phylogenetic clades are indicated. Violin plots and data points show the distribution of ipTM scores across twelve prediction runs (one per seed) with five AF3 models per run. **C)** Joint distribution of pTM and ipTM scores for individual MLO trimer models. For each MLO isoform, the model with the highest AF3 ranking score is labeled and highlighted. Axes were transformed using sigmoidal (logistic) functions to improve the visibility of data points. **D)** Domain architecture of *Hv*Mlo. Numbers indicate the first residue in each domain and the total sequence length. **NT**: extracellular/luminal N-terminal domain; **TM**: transmembrane domain; **IC**: intracellular loop; **AH**: amphipathic helix; **EC**: extracellular/luminal loop; **CT**: intracellular C-terminal domain; **CBD**: CAM-binding domain; **IDR**: intrinsically disordered region. **E)** Representative AF3 prediction of the *Hv*MloΔIDR homo-trimer (seed 1710, model 0) with domains of subunit A color-coded as shown in **D**. The surface of subunit B was colored according to its lipophopicity. Membrane boundaries were inferred using the PPM 3.0 web server and are indicated by transparent discs representing the outer and inner leaflets. **F, G)** Close-up views of AH1 shown in side view (**F**; corresponding to the orientation in **E**) and viewed along the helix normal (**G**). Helix residues are shown as sticks and are labeled. Hydrophobic/nonpolar residues are colored green, whereas cationic residues are colored petrol.

Further inspection of the predicted MLO assemblies indicated that all tested MLO isoforms adopt the same C3 architecture as observed for the *Hv*MloΔIDR trimer. We selected the models with the highest AF3 ranking scores for each isoform for visual analysis (Figure 3C; Supplemental Figure 6–Supplemental Figure 9; Supplemental File 4). Electrostatic surface coloring revealed variations in the charge distribution at the extracellular/luminal mouth of the predicted pores. For example, the *At*MLO3 model shows a pronounced density of anionic residues in this region (Supplemental Figure 9A), whereas others, such as the *At*MLO4 model (Supplemental Figure 6D), appear comparatively neutral. These differences at the extracellular/luminal pore opening may reflect isoform-specific properties that modulate ion conduction.

We revised the MLO membrane domain architecture for all 20 MLO proteins, using multiple sequence alignments (MSAs; Supplemental Figure 10A; Supplemental File 4). We found that the general core topology of MLOΔIDRs is conserved. For *At*MLO7, -8, and -10, we propose a putative additional N-terminal membrane-spanning domain (TM0) and an additional extracellular loop (EC0). Moreover, we detected differences in domain length and sequence composition for intracellular and extracellular/luminal regions, including EC1, EC3, IC2, and IC3. However, the relative positions of residues that form the interface between the three subunits are largely conserved (Supplemental Figure 10B). The predicted domain architecture of *Hv*Mlo (Figure 3D, E) matches its experimentally determined topology (Devoto *et al*., 1999). In addition, we identified a ten-amino acid amphipathic α-helix (AH1) in the middle of IC2 (Figure 3D, E, AH1). AH1 lies approximately parallel to the membrane plane and is positioned at the periphery of the inner leaflet as predicted by PPM 3.0 (Lomize *et al*., 2022). It comprises eight hydrophobic residues oriented toward the membrane core, whereas the basic residues R202 and R209 face the membrane–solvent interface (Figure 3F, G). In other ion channels, amphipathic helices coupled to the lipid bilayer convert membrane stretch into pore opening (Kefauver *et al*., 2020; Bavi *et al*., 2016). The conservation of AH1 (Supplemental File 4) and its membrane-associated positioning raise the possibility that membrane tension-promoted pore opening is a shared mechanism across MLO proteins.

### MD simulations reveal that conserved extracellular cysteine residues support the expansion of *Hv*Mlo and *At*MLO2 trimer pores during membrane tension

To assess whether MLO proteins respond to mechanical stimulation, we carried out all-atom MD simulations of the representative closed-state *Hv*MloΔIDR trimer model 1710_00. Compared with our initial simulations, we embedded the protein into an authentic plant plasma membrane model (Pogozheva *et al*., 2022) and carried out five 100 ns simulations while gradually decreasing pressures in the x–y plane (1, −20, −40, −60, and −80 bar; Supplemental Figure 11A–D; Supplemental Figure 12A, B; Supplemental File 3). Pore profiles were computed from the final frame of each simulation using MOLE. We did not observe substantial dilation when comparing minimal pore radii (r_min_(1 bar) = 0.599 Å, r_min_(−80 bar) = 0.678 Å; Supplemental Figure 11A–D), although per-residue RMSD values in the final models for the 1 bar and −80 bar simulations revealed substantial displacement at the periphery of the *Hv*MloΔIDR trimer (Supplemental Figure 11E). For instance, AH1 residues showed elevated backbone deviations (RMSD = 4.845 Å). By contrast, the pore-constricting I75 residues exhibited only minor displacement (RMSD = 0.303 Å), indicating that peripheral expansion was not transmitted to the pore bottleneck (Supplemental Figure 11E; ΔRMSD_AH1,I75_ = RMSD_AH1_ – RMSD_I75_ = 4.243).

We wondered whether disulfide bonds may alter force transmission within MLO trimers, as MLO proteins contain several conserved cysteine residues, including four invariant residues in EC1 and EC3 (Supplemental Figure 13; Supplemental File 4). In *Hv*Mlo, these four cysteines—C86, C98, C114, and C367—are essential for mediating powdery mildew susceptibility (Elliott *et al*., 2005). Our AF3 models show that disulfide bonds between C86 and C114 would couple the pore-forming helices TM2 and TM3, while C98–C367 would additionally bridge TM2/TM3 to the outer helices TM6 and TM7 near AH1 (Figure 4A, B). We introduced C86–C114 and C98–C367 into each subunit of the *Hv*MloΔIDR trimer and subjected the modified system to 50 ns MD simulations under progressively more negative x–y pressures (1, −40, and −80 bar; Supplemental Figure 12C, D; Supplemental File 3). Pore profiles and three-dimensional pore models inferred from the final frame of each simulation indicate that the pore radius increases with higher lateral tension (Figure 4C–F), consistent with the measured minimal pore radii (r_min_(1 bar) = 0.413 Å, r_min_(−40 bar) = 1.020 Å, r_min_(−80 bar) = 1.251 Å). Moreover, RMSD values of 1.689 Å for the I75 backbone and 4.229 Å for the AH1 backbone resulted in a markedly decreased ΔRMSD_AH1,I75_ of 2.540 Å compared to the *Hv*MloΔIDR trimer without disulfide bonds (Supplemental Figure 11E).

**Figure 4:**
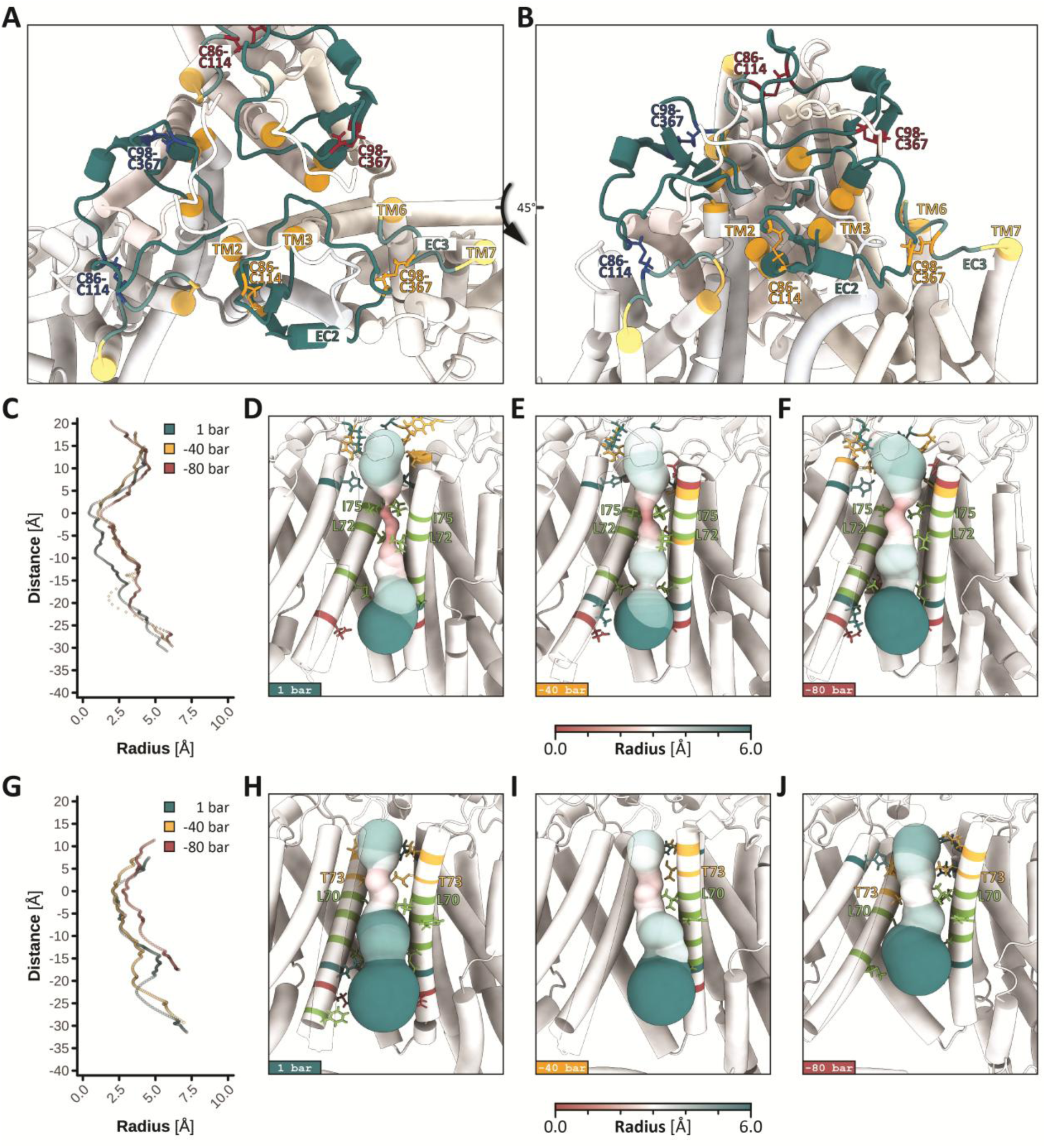
MD simulations reveal that disulfide bridges support the expansion of MLO trimer pores under membrane tension. **A, B)** Extracellular/luminal view of an *Hv*Mlo homo-trimer (seed 1710, model 0) with disulfide bonds between conserved extracellular residues C86–C114 and C98–C367. Disulfide bonds are color-coded by subunit (A, blue; B, yellow; C, red). EC1 and EC3 domains are shown in teal, while the extracellular/luminal ends of TM2, TM3, TM6, and TM7 are depicted in orange. **C, G)** Pore-radius profiles of *Hv*MloΔIDR (**C**) and *At*MLO2ΔIDR (**G**) models (both seed 1710, model 0 with disulfide bonds). Pore descriptors for the last frame of each 50 ns MD run were obtained *via* the MOLE API. Point colors indicate the x–y pressures during MD simulations. **D–F, H–J)** Three-dimensional “balloon” representations of transmembrane pores generated from MOLE output. To improve pore visibility, residues 1–50 of subunit B (left) are shown with reduced opacity, and subunit C is omitted. Pore-lining residues were identified in ChimeraX (*interface select*) and are shown as sticks, colored by polarity (nonpolar, green; polar, yellow; basic, petrol; acidic, red). Bottleneck residues are labeled.

*Hv*Mlo I75 is not conserved in *A. thaliana* and *M. polymorpha* MLO proteins (Supplemental File 4). At the corresponding position, *At*MLOs and *Mp*MLOs carry smaller and less hydrophobic residues. Such substitutions may influence pore hydration and the tension required for pore opening. To test this hypothesis, we subjected an *At*MLO2ΔIDR (UniProt: Q9SXB6) trimer (seed 1710, model 0), including the aforementioned disulfide bonds, to simulated lateral tension (1, −40, and −80 bar; Supplemental Figure 12E, F; Supplemental File 3). As for *Hv*Mlo, we observed progressive pore opening (Figure 4G–J) at the narrowest constriction (T73) yet with higher minimal pore radii (r_min_(1 bar) = 2.084 Å, r_min_(−40 bar) = 2.036 Å, r_min_(−80 bar) = 2.751 Å). We also tested an *Hv*MloΔIDR I75T mutant version (Supplemental Figure 11G–J; Supplemental Figure 12G, H; ^S^upplemental File ^3^) and measured minimal pore radii of r_min_(1 bar) = 1.292 Å, r_min_(−40 bar) = 1.606 Å, and r_min_(−80 bar) = 1.742 Å. Accordingly, the *Hv*MloΔIDR I75T mutant has a wider pore than the native variant, yet pore opening remained consistently smaller than observed for *At*MLO2ΔIDR. Notably, RMSD values for the AH1 domains of both models (RMSD(*Hv*MloΔIDR I75T) = 2.804 Å, RMSD(*At*MLO2ΔIDR) = 7.853 Å) suggest that the *At*MLO2ΔIDR trimer undergoes substantially larger overall expansion than the *Hv*MloΔIDR trimer under these conditions.

Finally, we simulated *Hv*MloΔIDR and *At*MLO2ΔIDR trimers carrying disulfide bonds under −80 bar lateral pressure, while applying a transmembrane electric field corresponding to a 0.5 V potential drop across the periodic box length. Under these conditions, we did not observe Ca²⁺ permeation, indicating that pore expansion at −80 bar lateral tension was insufficient to allow ion conduction.

### AF3 predictions reveal the binding modes of EF-hand proteins to *At*MLO2

CAM proteins bind to conserved leucine and tryptophan residues in the proximal CT of MLOs, upon association with Ca^2+^ at their four EF-hands (Bongartz *et al*., 2023; Kim *et al*., 2002). To test whether AF3 reproduces this interaction, we modeled *At*CAM2 (UniProt: P0DH97) in complex with different monomeric *At*MLO2 variants. In total, we tested twelve conditions using our standard AF3 setup (Figure 5A, B).

**Figure 5:**
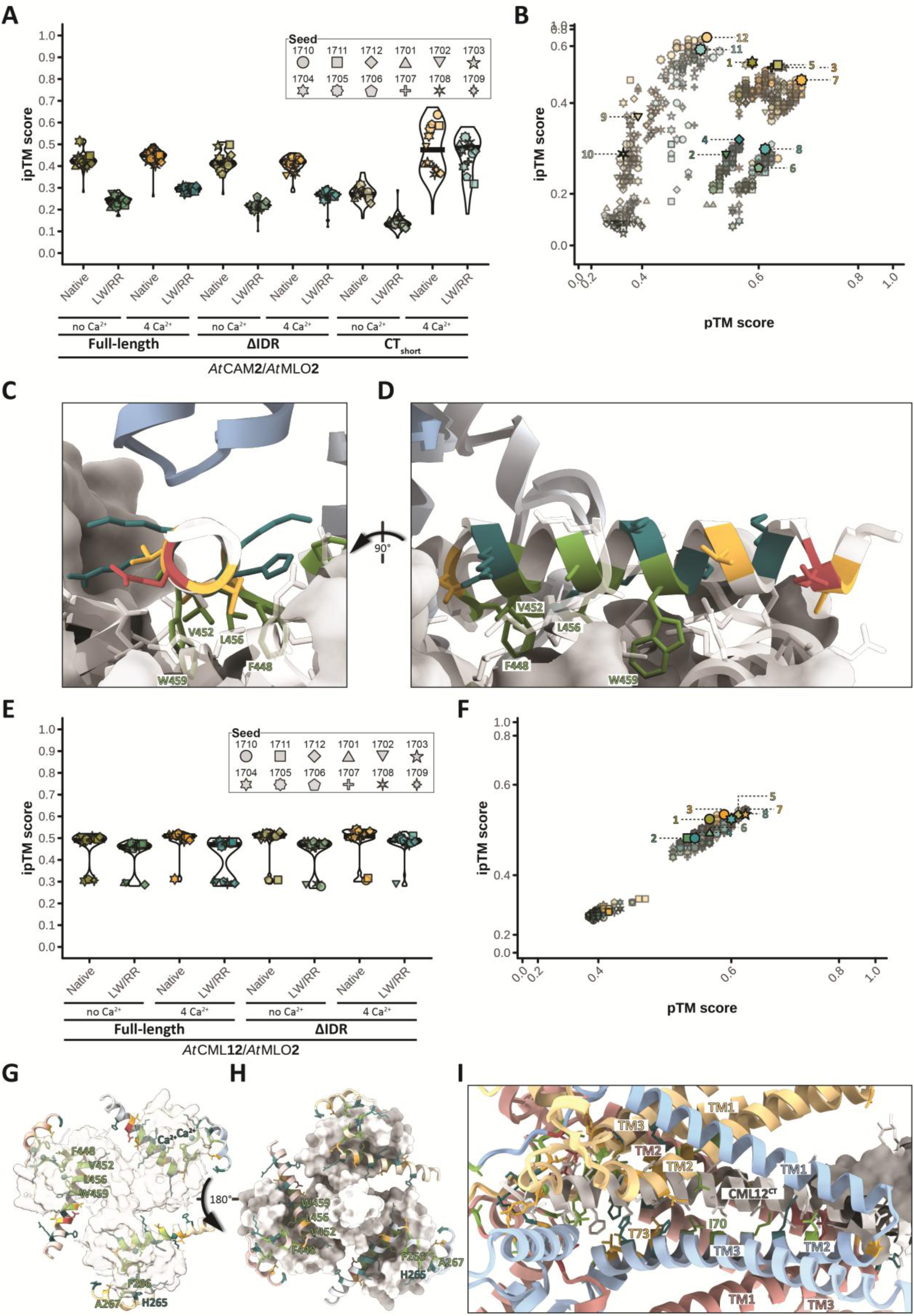
AF3 predicts binding of *At*CAM2 and *At*CML12 to *At*MLO2. **A, E)** Distribution of ipTM scores for AF3 predictions of one *At*MLO2 chain with one *At*CAM2 chain (**A**) or three *At*MLO2 chains with one *At*CML12 chain (**E**). *At*MLO2 was modeled as full-length, ΔIDR, or CTshort version (as indicated), and either as native or as L456R/W459R (LW/RR) mutant variant. Each condition was simulated with or without Ca²⁺. Violin plots and data points show the distribution of ipTM scores across twelve prediction runs (one per seed) with five AF3 models per run. **B, F)** Joint distribution of pTM and ipTM scores for individual *At*MLO2–*At*CAM2 (**B**) and *At*MLO2–*At*CML12 (**F**) complex models. For each condition, the model with the highest AF3 ranking score is highlighted and labeled according to its position on the x-axis in **A** or **E**. Axes were transformed using sigmoidal (logistic) functions to improve the visibility of data points. **C, D)** Close-up views of the CBD in the top-ranked *At*MLO2ΔIDR prediction (seed 1705, model 0) shown along the helix normal (**C**) and in side view (**D**). *At*CAM2-contacting residues are shown as sticks and colored by polarity (nonpolar, green; polar, yellow; basic, petrol; acidic, red); prominent residues are labeled. **G, H)** Cytosolic (**G**) and extracellular/luminal (**H**) views of the top-ranked *At*MLO2-*At*CML12 hetero-tetramer (seed 1710, model 0). For *At*MLO2, only contacting regions in the IC2s (263–273) and in the CBDs (439–476) are shown. *At*CML12-contacting residues are displayed as sticks, colored by polarity. Prominent residues and Ca^2+^ are labeled for one MLO subunit. **I)** Close-up view of the pore region (residues 1–199) of the full-length *At*MLO2 trimer in complex with the C-terminus of *At*CML12 (seed 1710, model 0). Transmembrane domains and prominent residues are labeled. *At*CML12-contacting residues are shown as sticks, colored by polarity.

*At*MLO2 L456R, W459R mutants (LW/RR) reduce CAM binding *in vitro* and *in vivo* (Bongartz *et al*., 2023). For full-length *At*MLO2 and *At*MLO2ΔIDR, AF3 ipTM scores suggested reduced *At*CAM2 affinity for LW/RR mutants (ipTM ∼ 0.25) compared with the corresponding native *At*MLO2 versions (ipTM ∼ 0.45; Figure 5A), while the presence of Ca²⁺ did not markedly affect ipTM scores. By contrast, for the CT_short_ variants (residues 439–573), Ca²⁺ supplementation substantially increased ipTM scores for both the native variant (ipTM from ∼ 0.25 to ∼ 0.50) and the LW/RR mutant (ipTM from ∼ 0.12 to ∼ 0.50), whereas the LW/RR mutations no longer strongly reduced ipTM scores. Inspection of pTM scores indicated that CT_short_ predictions had comparatively low overall fold confidence (Figure 5B), likely reflecting the higher intrinsically disordered content of the CT-only construct. Notably, this decrease in pTM did not preclude high ipTM values. A similar, albeit less pronounced, trend was observed for full-length versus ΔIDR versions, as ΔIDR variants generally exhibited higher pTM scores than their full-length counterparts (Figure 5B).

Consistent with the experimentally defined CAM-binding domain (CBD; Bongartz *et al*., 2023), *At*CAM2 binds to residues 442–467 in *At*MLO2 (Supplemental File 5). Notably, W459 shows the highest maxContact score (0.76) among *At*MLO2 residues (Supplemental Figure 14A; Supplemental File 5) and penetrates deeply into one of the EF-hand-containing CAM lobes (Figure 5C, D). In addition to L456 and W459, the hydrophobic residues F448 and V452 contribute to *At*CAM2 binding (Figure 5C, D), which is also reflected by high maxContact scores (F448 = 0.70; V452 = 0.76) across all Ca²⁺-supplemented predictions for native *At*MLO2 variants (Supplemental Figure 14A; Supplemental File 5). Moreover, AF3 predictions consistently indicate a previously undescribed contact of *At*CAM2 to the distal part of the *At*MLO2^IC2^ that is formed by residues H265, F266, A267, N270, and R273 (maxContact = 0.69, 0.72, 0.69, 0.54, 0.49; Supplemental Figure 14A; Supplemental File 5).

Although AF3 captures key characteristics of the association between MLO and CAM proteins, *At*CAM2 comprises four EF-hands (i.e., two MLO-binding sites), while MLO trimers have three CBDs. Non-mutually exclusive scenarios reconciling this stoichiometric mismatch are (i) that the physiologically relevant oligomeric assembly is the MLO dimer (Supplemental Figure 1B), (ii) that multiple CAM lobes engage a single target helix, as described for other channels (Larsen *et al*., 2024), or (iii) that a single MLO chain harbors multiple CBDs, for example, in the MLO^IDR^, as also observed in some predictions (Supplemental Figure 14A; Supplemental File 5). Alternatively, EF-hand proteins beyond canonical CAMs may bind to MLO proteins. In *A. thaliana*, *At*CML12 (UniProt: P25071) is the only six-EF-hand protein (McCormack and Braam, 2003), providing a stoichiometric match for three CBDs. Notably, *At*CML12 interacts with *At*MLO4 (Zhu *et al*., 2021). We modeled full-length and ΔIDR *At*MLO2 trimers with a single *At*CML12 chain. For all conditions, we observed bimodal distributions of ipTM scores with higher-confidence models around 0.50 and lower-confidence models around 0.30 (Figure 5E), consistent with closed and dynamic conformational states of the *At*MLO2 trimer pore. This pattern was also reflected in the corresponding pTM scores (Figure 5F). As observed for the *At*MLO2–*At*CAM2 complex (Figure 5A, B), the LW/RR mutation had a stronger effect on *At*CML12 binding than Ca^2+^ supplementation (Figure 5E, F)

AF3 per-residue metrics (Supplemental Figure 15A; Supplemental File 6), together with visual inspection of models (Figure 5G, H; Supplemental Figure 15B), indicate that one EF-hand–containing lobe of *At*CML12 engages one CBD per *At*MLO2 protomer. Similar to *At*CAM2, residues F448, V452, L456, and W459 in the CBDs contribute prominently to *At*CML12 binding, whereas H265, F266, and A267 in the IC2s seem to stabilize the hetero-tetramer (Figure 5G, H). Compared with the *At*MLO2–*At*CAM2 hetero-dimer, maxContact scores for these residues are modestly reduced (Supplemental Figure 15A; Supplemental File 6), potentially reflecting promiscuity in how *At*CML12 lobes dock onto different *At*MLO2 subunits.

We also noticed that in some—but not all—models, the extended C-terminal tail of *At*CML12 inserts into the *At*MLO2 trimer pore (Figure 5I; Supplemental Figure 15B). This insertion was more frequent in lower-confidence AF3 predictions than in higher-confidence predictions (i.e., closed models; Figure 5I; Supplemental Figure 15B). Consistent with this notion, average maxContact scores for the pore-lining residues L70 and T73 were higher in lower-confidence models (0.73 and 0.66, respectively) than in higher-confidence models (0.24 and 0.16), considering only native, Ca²⁺-supplemented predictions (n = 120; 15 lower-confidence and 105 higher-confidence models). This preference of the *At*CML12^CT^ for the open conformation of the *At*MLO2 trimer is consistent with a potential pore-occluding mechanism that may act in response to Ca²⁺ influx.

### AF3 pinpoints EXO70-binding sites in MLO proteins

MLO proteins interact with specific EXO70 proteins, which is, for example, physiologically relevant for the polarized secretion of callose synthases in trichomes (Huebbers *et al*., 2024). To disentangle interaction specificity between MLO and EXO70 proteins, we carried out a comprehensive LCI screen in *N. benthamiana* leaves covering about 2160 data points (Figure 6A; Supplemental Figure 16A, C, E, G; Supplemental Figure 17A, C, E, G, I) and complemented these experimental data with 5400 AF3 models (Figure 6B; Supplemental Figure 16B, D, F, H; Supplemental Figure 17B, D, F, H, J).

**Figure 6:**
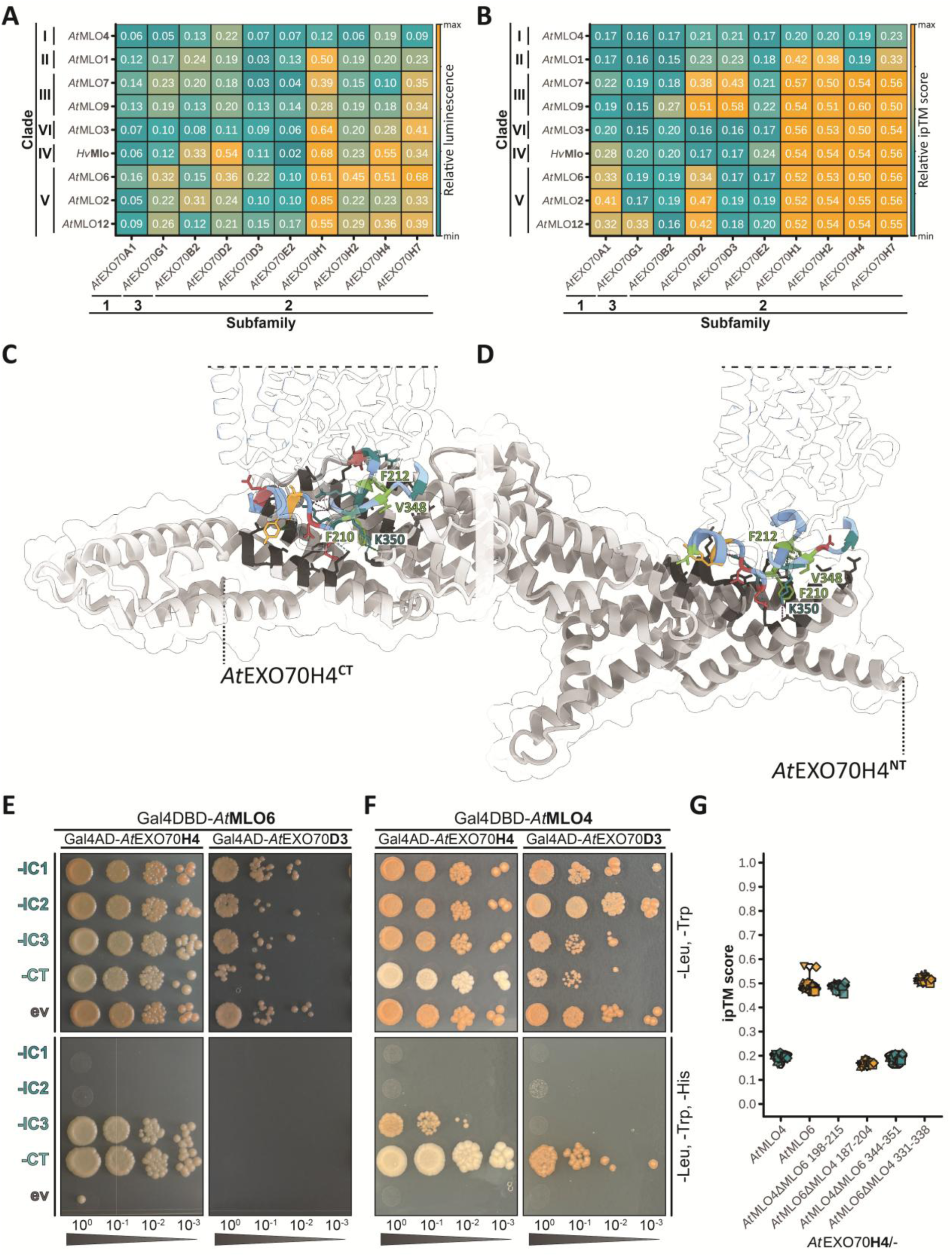
AF3 predictions and protein-protein interaction experiments reveal *At*EXO70-*At*MLO binding modalities. **A, B)** Heatmaps of LCI-based (**A**) and AF3-predicted interaction data for *A. thaliana* MLO and EXO70 isoforms across clades. Colors and numbers represent mean luminesce relative to the interaction of the indicated MLO protein with *At*CAM2 (**A**) or mean ipTM scores across twelve runs (one per seed) with five models per interaction (**B**). **C, D)** Top-ranked AF3 models for *At*MLO6 binding to the C-terminal (seed 1702, model 0; **C**) or N-terminal (seed 1701, model 0; **D**) cavity of *At*EXO70H4. *At*MLO6 models are shown semi-transparently without their C-termini (residues 437–583) and cropped at the dashed lines. *At*EXO70H4-contacting residues in *At*MLO6 are shown as sticks and colored by polarity (nonpolar, green; polar, yellow; basic, petrol; acidic, red). Colored dashed lines indicate salt bridges (purple), hydrogen bonds (cyan), and aromatic interactions (green). Prominent residues are labeled. *At*EXO70H4 is shown without its terminal IDRs (residues 1–29 and 603–628). Residues within 4 Å of the colored residues in *At*MLO6 are shown in dark gray. **E, F)** Y2H assays probing interactions of intracellular domains from *At*MLO6 (**E**) and *At*MLO4 (**F**) with *At*EXO70H4 and *At*EXO70D3. Yeast cultures were spotted as 1:10 serial dilutions onto control medium lacking leucine and tryptophan (-Leu, -Trp) or selection medium additionally lacking histidine (-Leu, -Trp, -His). **AD**: activation domain; **DBD**: DNA-binding domain; **IC**: intracellular loop; **CT**: C-terminus; **ev**: empty vector. **G)** Distribution of ipTM scores for AF3 predictions of native and domain-swapped *At*MLO4 and *At*MLO6 variants modeled with *At*EXO70H4. Violin plots and data points show the distribution of ipTM scores across twelve prediction runs (one per seed) with five AF3 models per run.

The LCI data indicated that *At*MLO proteins from clades II–VI interact with members of the *At*EXO70H clade, whereas clade I *At*MLO4 did not show detectable *in planta* interaction with the tested *At*EXO70 proteins (Figure 6A). Notably, *At*MLO4 produced a robust signal with *At*CAM2, demonstrating its accumulation in the heterologous system. AF3 hetero-dimer predictions largely recapitulated the preferences of *At*MLO proteins (other than clade I) for *At*EXO70H isoforms (Figure 6B). However, for *At*MLO clades III–VI, AF3 predicted broadly similar mean ipTM values (mean ± SD = 0.54 ± 0.04) across *At*EXO70H isoforms, whereas the LCI signals displayed substantial isoform-specific variation (mean ± SD = 0.39 ± 0.19). Beyond the *At*EXO70H clade, AF3 predicted comparatively strong associations of *At*EXO70D- isoforms with clade III and clade V *At*MLO proteins (Figure 6B), which was only partially reflected in the LCI data (Figure 6A). Lastly, *At*EXO70A1 and -G1 belong to the EXO70.I and EXO70.III subfamilies that likely carry out canonical EXO70 functions (La Concepcion *et al*., 2025). Whereas LCI assays yielded modest signals (∼0.30) for clade V *At*MLO2, -6, and -12 isoforms with *At*EXO70G1 (Figure 6A), AF3 predictions exhibit increased ipTM scores (∼0.35) for the association of these MLO proteins with *At*EXO70A1 (Figure 6B).

For AF3 predictions testing associations between *At*EXO70H isoforms and clade V MLO proteins, ipTM scores were frequently bimodally distributed (Supplemental Figure 17F, H, J). For example, for the *At*MLO6-*At*EXO70H4 hetero-dimer we observed a higher-confidence cluster (ipTM ∼0.60) that represents *At*MLO6 binding near the CT of *At*EXO70H4 and a lower-confidence cluster (ipTM ∼0.50) that reflects *At*MLO6 binding towards the NT of *At*EXO70H4 (Supplemental Figure 18A, B). By contrast, clade III *At*MLO9 was predicted to bind exclusively to the C-terminal cavity of *At*EXO70H4 (Supplemental Figure 18C, D). In these predictions, residues in *At*EXO70H domains A and B form the N-terminal MLO-binding cavity, whereas residues in the C domain form the C-terminal cavity (Supplemental Figure 18E, F), raising the possibility that *At*EXO70H isoforms support multivalent interactions with MLO proteins.

For EXO70-binding sites in MLO proteins, maxContact scores indicate that the interface involves a region in the proximal IC2 (i.e., upstream of AH1) and the central part of IC3, although we observed occasional, erratic contacts to the IDR of the MLO^CT^ (Supplemental File 8). Inspection of *At*MLO6–*At*EXO70H4 heterodimers shows that three residues each in the relevant IC2 (F210, R211, and F212) and IC3 (V348, V349, and K350) segments fold into two antiparallel β-strands that form an exposed patch at the cytosolic edge of the *At*MLO6 core (Figure 6C, D). Among these, F210, F212, V349, and K350 exhibit high average maxContact scores (0.80, 0.71, 0.76, and 0.71, respectively; Supplemental File 8). Notably, *At*MLO6 R209 shows the highest average maxContact score (0.86) and consistently exceeds 0.80 across interactions of clade III–VI MLO proteins with *At*EXO70H, but not *At*EXO70D, isoforms (Supplemental File 8).

We previously probed the binding of *At*EXO70H4 to the IC2 and the CT of *At*MLO2, -6, and -12 in Y2H assays (Huebbers *et al*., 2024) and found that *At*EXO70H4 binds the MLO^CT^, but not the MLO^IC2^. Following our AF3 predictions, we repeated these experiments, including the MLO^IC1^ and the MLO^IC3^. Adding to the previously described interaction with the MLO^CT^, we detected interactions of the IC3 domains of *At*MLO2, *-*6, and *-*12 with *At*EXO70H4 but not *At*EXO70D3 (Figure 6E; Supplemental Figure 19A–D). We also tested the intracellular domains of *At*MLO4 (clade I) and likewise found that the IC3 and the CT of *At*MLO4 bind to *At*EXO70H4 but not *At*EXO70D3 (Figure 6F), although yeast growth was generally weaker for the *At*MLO4^IC3^ than for the AtMLO4^CT^ or the corresponding interacting domains from the clade V MLO proteins (Figure 6E). For IC1 and IC2 constructs, as well as empty-vector controls, we did not observe growth on selection plates despite accumulation of the recombinant proteins (Supplemental Figure 19E, F). We complemented this expanded Y2H setup with AF3 predictions. To this end, we exchanged the predicted EXO70-binding regions in *At*MLO6 (IC2: 198–215; IC3: 344–351) with the corresponding regions from *At*MLO4 (IC2: 187–204; IC3: 344–351) and *vice versa* and submitted these domain-swapped versions for complex predictions with *At*EXO70H4. Replacing the IC3 region in *At*MLO6 did not diminish the predicted interaction with *At*EXO70H4 (ipTM ∼0.52). By contrast, replacing the *At*MLO6^IC2^ region with its *At*MLO4 counterpart reduced the ipTM score to ∼0.17, comparable to that of the *At*MLO4-*At*EXO70H4 hetero-dimer (∼0.20; Figure 6G). Likewise, introducing the *At*MLO6^IC3^ region into *At*MLO4 did not alter its predicted affinity for *At*EXO70H4 (ipTM ∼0.19), whereas the *At*MLO6^IC2^ region in *At*MLO4 improved complex confidence to ∼0.50, similar to that for the *At*MLO6-*At*EXO70H4 hetero-dimer (∼0.49; Figure 6G). Together with our Y2H data, these predictions suggest that the MLO^IC3^ harbors an *At*EXO70H4 interaction site that is conserved across clade I and clade V MLO proteins, whereas the binding region in MLO^IC2^ is not conserved and determines clade-specific EXO70 binding.

## Discussion

Given the disruptive potential of AI-based protein structures, it remains essential to constantly test the plausibility of predictions against experimental data. Here, we report an AF3-based trimeric architecture for MLO proteins that agrees with previous experimental data collected on this plant-specific protein family. We propose that MLO are gated by a tension-based mechanism, suggesting that they represent previously undiscovered mechanosensitive channels that are subject to feedback inhibition by the six-EF-hand protein *At*CML12. Moreover, we present a comprehensive comparison of LCI and AF3 data that not only pinpoints mutual binding sites in MLO and EXO70 proteins, but also supports the feasibility of AF3 for interaction screens. Lastly, AF3 produced multiple plausible conformations for some of these assemblies in our simulations, suggesting that it captures at least part of the conformational heterogeneity and dynamics of protein complexes.

AF3 predicted two plausible oligomeric states for *Hv*Mlo, a dimer and a trimer (Figure 1D). This is consistent with the finding that ipTM score distributions can capture multiple physiologically relevant states of ion channels (Lin *et al*., 2026). Where experimental structures were available, these predictions agreed with known oligomeric assemblies. For *Hv*Mlo, this raises the possibility that both oligomeric states, the dimer and the trimer, occur *in planta*. Whereas the trimer is a plausible ion-conducting pore, the function of the dimer remains unclear. One possibility is that the dimer represents an intermediate during trafficking, as oligomerization may be a requirement for ER exit and transport to the correct membrane compartment (Martzoukou *et al*., 2015). Transporting *Hv*Mlo proteins as a dimer rather than a trimer may prevent accidental ion flux during trafficking. Therefore, the trimeric *Hv*Mlo pore may assemble and stabilize preferentially at its target membrane, which has been described as “kinetic trapping” (Anderluh *et al*., 2017).

We further suggest that lower-confidence MLO trimer models capture conformational heterogeneity linked to MLO pore gating. Previous studies have shown that AF2 may produce different conformational states, although, to our knowledge, this usually required encouraging AF2 to sample alternative conformations (Tao and Corry, 2025; Pinto-Anwandter, 2025; Lopez-Mateos *et al*., 2026; Lidbrink *et al*., 2025; Torres *et al*., 2024; Ngo *et al*., 2025). We did not explicitly force AF3 to generate alternative states for MLO homo-trimers or MLO hetero-complexes with *At*CML12 and EXO70 proteins (Figure 5; Figure 6). Instead, clustering models by ipTM and pTM scores provided a practical way to identify multiple plausible conformations from AF3 outputs. Therefore, we suggest that lower-scoring models, if they co-occur with higher-scoring models, do not necessarily represent failed predictions, as reduced confidence may arise from increased structural variability within the same prediction run.

We hypothesized that MLO gating is driven by a dragging model (Kefauver *et al*., 2020), following the force-from-lipids paradigm, due to a conserved amphipathic helix (AH1) positioned at the periphery of the predicted trimers (Figure 3F–G). All-atom MD simulations show that AH1 is indeed dragged outwards during membrane expansion, while force transmission to the pore-lining TM2 domains requires a total of six disulfide bonds in the extracellular loops of *Hv*Mlo and *At*MLO2 (Figure 4). Simulated pressures often exceed physiological gating thresholds to accelerate opening within accessible simulation times (Vecchis *et al*., 2021). However, the minimum applied lateral pressure of −80 bar in our simulations was more extreme than pressures reported for other channels such as PIEZO1 (-40 bar; Vecchis *et al*., 2021) or TREK-2 (-50 bar; Aryal *et al*., 2017). Although, a complete mechanical interpretation requires analysis of membrane pressure anisotropy and local lateral pressure profiles (Aryal *et al*., 2017), channel geometry may be important here. For example, the extended PIEZO triskelion (Lin *et al*., 2019) has larger lever arms that likely transmit membrane tension to the pore more effectively than AH1 in the compact MLO trimers.

Typical minimum pore radii that allow the transmission of hydrated Ca²⁺ are on the order of 3.0–4.0 Å (Demidchik *et al*., 2018; Liu *et al*., 2018; Zhao *et al*., 2016). This is in line with the tested open *Hv*Mlo trimer (model 1705_02, r_min_ = 3.003 Å). *At*MLO2’s minimal pore radius after -80 bar lateral tension (2.751 Å) is presumably just below the reported 3.0 Å threshold, whereas *Hv*Mlo remained markedly narrower after simulation (r_min_ = 1.251 Å). In addition, AF3 predicts that the *Hv*Mlo trimer pore adopts even more dilated states exceeding 5 Å (Figure 4D), suggesting that our MD setup does not capture the full expansion potential of MLO channels. Several factors may contribute to this discrepancy: (i) MLO trimer gating might be facilitated by cytoskeletal or cell wall elements (force-from-filaments). (ii) The C-terminal IDRs of MLO proteins are not present in the tested AF3-derived models and may be important for mechanical force transmission, as described for cytosolic IDRs in PIEZO2 (Verkest *et al*., 2022). (iii) Crowding of MLO proteins within a confined membrane area may streamline force transmission, as reported for PIEZO1 (Jiang *et al*., 2021). (iv) Although we used an authentic plant plasma membrane model during simulations (Pogozheva *et al*., 2022), MLO proteins may require a different lipid environment for full expansion. Altogether, MLO pore opening probably depends on cellular contexts and/or membrane conditions beyond those captured in our current MD simulations.

In addition to Ca^2+^, we observed that Cl⁻ counterions translocated from the intracellular to the extracellular/luminal space during MD simulations. However, under physiological electrochemical conditions, sustained Cl⁻ efflux to the extracellular space would not be favored (Geilfus, 2018). Furthermore, Cl⁻ ions did not form stable interactions with pore-lining residues, suggesting that their passage has been assisted by the applied electric field. By contrast, we identified several acidic residues in or near the *Hv*Mlo pore (E35, E55, E58, E63, and D79) that coordinated Ca^2+^ ions during MD simulations and may contribute to ion selectivity and channel gating (Figure 2E, F, H, I; Supplemental Figure 4D, E). Ca^2+^ may also function as an allosteric activator of MLO channels, as described for other channels (Lam *et al*., 2021). Extracellular/luminal residues at the pore entrance, such as D79, may contribute to selectivity among different cations (Feng *et al*., 2001; Zhao *et al*., 2016; Voets *et al*., 2002; Xue *et al*., 2025). Although D79 itself is not conserved, acidic residues are common in the EC1 domain of MLO proteins. Experimental data showed that *Hv*MLO, *At*MLO2, and *At*MLO7 conduct currents carried by Ba^2+^ and Mg^2+^, but not by monovalent cations such as K^+^ and Na^+^, whereas the Ca^2+^ channel activity of *At*MLO7 was inhibited by La^3+^ and Gd^3+^ (Gao et al., 2022). Thus, it will be interesting to see how these ions behave during MD simulations with the predicted structures of MLO proteins.

We found that disulfide bonds between conserved extracellular cysteines facilitate force transmission between outer complex domains such as AH1 and the pore-constricting residues (Figure 4B–D). These cysteines are invariant across the 20 MLO proteins analyzed here and are conserved in >90% (C98, C114) or >95% (C86, C367) of land-plant MLOs (Kusch *et al*., 2016). In line with our models, they have previously been proposed to form disulfide bridges (Elliott *et al*., 2005) and *Hv*Mlo cysteine-to-alanine mutant variants showed reduced—yet not abolished—protein accumulation. Notably, relative accumulation was similar for the paired mutants C86A/C114A and C98A/C367A, consistent with the disulfide pairings suggested by our AF3 models. Moreover, all four cysteines were required for *Hv*Mlo to complement powdery mildew susceptibility in barley *mlo* mutants. Together, these genetic data and our observations support the idea that MLO channel activity (i.e., Ca²⁺ flux) contributes to successful powdery mildew entry.

Apart from homo-oligomers, we also probed MLO proteins for association with their experimentally validated interactors, including EF-hand proteins such as calmodulin and EXO70 exocyst complex components. Notably, AF3 still suggested residual *At*CAM2 binding to the tested LW/RR variants, consistent with reported *in planta* interaction data (Bongartz *et al*., 2023; Yuan *et al*., 2025). Our predictions suggest that this residual binding *in planta* may have three, not mutually exclusive, explanations: (i) Beyond L456 and W459, AF3 predicts additional *At*MLO2 residues that contribute to calmodulin binding. These include the hydrophobic residues F448 and V452, both of which showed higher maxContact scores than L456. To the best of our knowledge, neither residue has yet been investigated in protein–protein interaction assays. (ii) Our models suggest that *At*CAM2 binding is further stabilized by residues in the IC2 domain. Although these residues are less conserved than those in the CBD, they represent potential targets for future site-directed mutagenesis. (iii) Lastly, in addition to homo-oligomerization, MLO proteins likely also hetero-oligomerize with phylogenetically related MLO isoforms (Jones *et al*., 2017). This raises the possibility that, *in planta*, MLO hetero-complexes may contain wild-type subunits that mediate calmodulin binding and thereby generate a residual interaction signal.

AF3 predicted that each lobe of *At*CML12 engages one CBD of the *At*MLO2 trimer (Figure 5E–H). Notably, the extended C-terminal tail of *At*CML12 inserted into the *At*MLO2 trimer pore, particularly in lower-confidence or open *At*MLO2 predictions. This observation suggests that *At*CML12 acts as a negative-feedback regulator that terminates MLO-mediated Ca^2+^ flux upon Ca^2+^ influx. A similar hypothesis has previously been proposed for calmodulin (Gao *et al*., 2022). *AtCML12* belongs to the touch-inducible (*TCH*) genes in *A. thaliana* and is allelic to *TCH3* (*AtTCH3*; Braam and Davis, 1990; Sistrunk *et al*., 1994). Thus, *At*CML12 may act downstream of mechanosensitive MLO Ca^2+^ channels to decode mechanically induced Ca^2+^ signatures. This may occur either through effects on downstream signaling cascades (Sun *et al*., 2022) or through direct physical association of *At*CML12 with MLO trimers.

For the interaction of MLO and EXO70 proteins, we found a substantial agreement between experimental data and AF3-predicted interactions. Therefore, we conclude that AF3 reliably captures authentic binding modes for MLO–EXO70 hetero-dimers. The finding that *At*EXO70H4 contains two potential binding sites for MLO proteins is particularly interesting, as our previous study raised the question of whether MLO proteins recruit *At*EXO70H4 to distinct plasma-membrane domains, or *vice versa* (Huebbers *et al*., 2024). The presence of two binding sites in *At*EXO70H4 regard that MLO proteins recruit other MLO proteins to specific membrane domains with *At*EXO70H4 acting as an adaptor. Whether this occurs in the presence or absence of the exocyst holo-complex remains to be determined. However, recent data suggest that EXO70.2 subunits may have escaped their canonical role within the exocyst complex (La Concepcion *et al*., 2025). From a steric perspective, our models suggest that the binding sites in *At*EXO70H4 favor the simultaneous binding of two MLO proteins residing in the same membrane (Figure 6C, D). Therefore, we *At*EXO70H4 may also bridge MLO protomers in adjacent trimers to facilitate or even synchronize force transition during channel opening.

To date, five families of mechanosensitive ion channels have been identified in plants—MSL, MCA, TPK, PIEZO, and OSCA channels (Kaur *et al*., 2021). However, plant mechanoperception likely involves additional mechanosensory modules including other channel families (Hamilton *et al*., 2015; Demidchik *et al*., 2018; Ali *et al*., 2025; Tyagi *et al*., 2023) and the identities of these channels are still being resolved (Zhou *et al*., 2025). Our data strongly support the notion that MLO proteins adopt a conserved homo-trimeric architecture that permits tension-driven pore expansion in MD simulations. As this is in line with the finding that different members of the MLO family function as channels for divalent cations (Gao *et al*., 2023; Gao *et al*., 2022; Luan, 2026), we propose that MLO proteins act as mechanosensitive channels in plants. This interpretation is consistent with the reported roles of MLO proteins in cells that exhibit Ca^2+^ signals in response to tactile stimulation, including pollen tubes (Meng *et al*., 2020), synergid cells (Ponvert and Johnson, 2024; Ju *et al*., 2021), root tips (Bidzinski *et al*., 2014; Binci *et al*., 2025), root hairs (Ogawa *et al*., 2025), and trichomes (Huebbers *et al*., 2024; Matsumura *et al*., 2022). Moreover, a mechanosensitive function of MLO proteins would also be consistent with their established role in powdery mildew host cell entry, because penetration by fungal hyphae is itself expected to impose mechanical stress on the host plasma membrane. However, the molecular basis by which MLO proteins promote powdery mildew susceptibility remains elusive. We speculate that, upon CML12 binding, MLO proteins may stabilize stretched membrane conformations and thereby support the establishment of bent or curved membrane shapes that would facilitate fungal entry into host cells. In this scenario, the relevant MLO proteins represent previously unrecognized molecular touch receptors during fungal invasion (Ryder *et al*., 2025), albeit ones that act in favor of the pathogen, at least in powdery mildew interactions. This hypothesis will require further experimental testing, for example by assessing powdery mildew susceptibility in *cml12* mutants.

We show that distinct pore states captured by AF3, together with MD simulations, provide a useful framework for investigating MLO channel gating. However, experimental validation will be required to corroborate these findings, for example through patch-clamp measurements under controlled membrane stretch or mechanical stimulation combined with Ca^2+^ imaging in wild type and *mlo* mutants. Moreover, MLO proteins are associated with cytoskeletal dynamics (Opalski *et al*., 2005; Miklis *et al*., 2007), cell wall biogenesis (Huebbers *et al*., 2024; Cao *et al*., 2024), and regulators of mechanotransduction, including receptor-like kinases such as FERONIA (Leicher *et al*., 2025; Kessler *et al*., 2010; Ogawa *et al*., 2025). It will therefore be interesting to determine how these components, together with EXO70 proteins, integrate into a functional mechanosensitive module.

## Materials and Methods

### Code development and availability

All code used in this study is publicly available on GitHub (Table 1). Generative AI assisted in drafting and debugging Python, R, and Bash scripts. All code was critically reviewed, tested, and revised by the authors, who take full responsibility for the final code, analyses, and results.

**Table 1:**
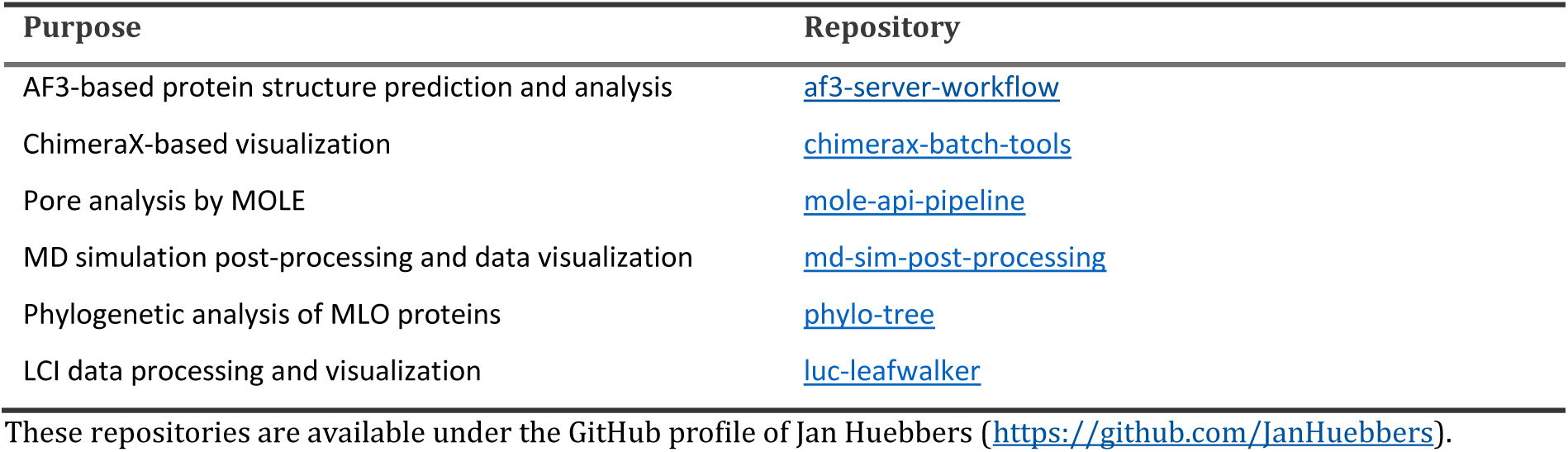
GitHub repositories associated with this study.

### AF3-based protein structure prediction and analysis

For AF3-based structure prediction and the downstream extraction and analysis of the resulting data, we organized our workflow into different modules written in Python (version 3.10.16). In module 1, we generated JSON files for AF3 server input and MSAs used for the mapping of per-residue metrics in module 2. The parameters for each JSON file and AF3 run were specified in a spreadsheet (.xlsx format). Each run usually comprised twelve jobs, each yielding five structural models. These parameters included the input chains to be retrieved from a folder containing sequence files (.fasta format), the ion type and ion count, if applicable, and the job-specific AF3 seed (here: 1710, 1711, 1712, 1701, 1702, 1703, 1704, 1705, 1706, 1707, 1708, 1709), which were chosen based on the date October 17. A configuration file (.yml format) contained run-specific information such as the run identifier, the alignment algorithm to be used (usually Clustal Omega), and plot-specific parameters for module 3. The resulting JSON input files were uploaded to the AF3 server (accessed between October 2025 and March 2026). After modeling, the output of each run, usually comprising twelve jobs, was downloaded as a single compressed archive and unpacked into a target folder on a Linux system.

In module 2, AF3 structure files (.cif format) were used to generate structural alignments of the five models per job in ChimeraX (version 1.10.1; Pettersen *et al*., 2021). AF3 summary_confidence.json files were parsed to retrieve global confidence metrics such as pTM, ipTM, and ranking scores, whereas AF3 full_data.json files were parsed for per-residue metrics, including distance, iPAE, and contact scores. The latter served as input for the calculation of minD, min-iPAE, and maxContact scores. Here, *i* denotes the residue of interest, *j* denotes residues in the interacting partner chain, and *c_i_* and *c_j_* indicate the corresponding chain identities. For each residue *i*, inter-chain residue-pair matrices were reduced to per-residue summary vectors by taking the minD, the min-iPAE, or the maxContact score across all residues *j* in the interacting partner chain, while retaining only inter-chain residue pairs (*c_i_* ≠ *c_j_*).

For minD:

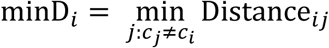

For min-iPAE:

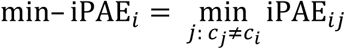

min-iPAE values were normalized to an approximate 0–1 range:

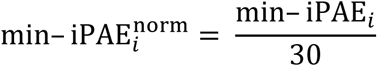

For maxContact:

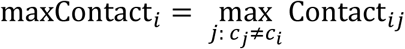

When multiple models were available, the corresponding per-residue summary vectors were averaged across models. The resulting per-residue extreme metrics were then mapped onto the MSAs generated in module 1 and exported to spreadsheet files (.xlsx format), in which per-residue metrics were displayed as heat maps and residues were color-coded according to their polarity.

In module 3, R (version 4.4.3) was used for data visualization by ggplot2 (version 3.5.2; Wickham, 2016). For individual runs, this module generated ipTM dot plots, pTM–ipTM scatter plots, and per-residue score heat maps. In module 4, data from different simulation runs were merged to generate datasets and plots across multiple runs, whereas in module 5, different merged datasets are combined to create summary plots and heat maps across multiple merged runs, for example for *At*EXO70/MLO matrices. All proteins used for AF3 predictions in this study, including their UniProt identifiers, are listed in Supplemental Table 1.

### Pore analysis by MOLE

We used the MOLE application programming interface (API; https://api.mole.upol.cz/; accessed between December 2025 and March 2026) to automatically submit PDB files to the MOLE server (Raček *et al*., 2025) for pore prediction. Submission of PDB files and extraction of the resulting data were implemented in Python (version 3.10.16) and organized using the Snakemake workflow management system (version 9.18.2; Mölder *et al*., 2021). For each structure, the input PDB file was first uploaded to the /Init endpoint of the MOLE API to obtain a computation ID. Computations were then submitted *via* the /Submit endpoint, in “Pores” mode, using job-specific parameters passed as a JSON object. To account for transient server-side issues, submission requests were automatically retried. After submission, computation metadata were retrieved via the /CompInfo endpoint and the corresponding submission was identified by its SubmitId. Job status was monitored through repeated queries to the /Status endpoint at 15-s intervals until completion, with a timeout of 1800 s. In cases where the API returned a transient “SubmitId not found” error, computation metadata were refreshed and the most recent available submission was used. After successful completion, results were downloaded via the /Data endpoint as a JSON file and a report archive. From the latter, per-sample pore-profile tables corresponding to the top-ranked predicted pore (path_1.csv) were merged into a single long-format table in .csv format. In R (version 4.4.3), pore-axis coordinates were reoriented on a per-sample basis to ensure a consistent bottom-to-top orientation across structures. For merged analyses, absolute pore-distance coordinates were additionally aligned to the bottleneck position, defined as the location of minimum of the pore radius. The resulting pore profiles were visualized as radius-versus-position plots using ggplot2 (version 3.5.2; Wickham, 2016). Three-dimensional pore models in .json format were imported into ChimeraX with the *format mole* option and superimposed on the corresponding input structures for visualization.

### Molecular Dynamics simulations and analysis

All systems subjected to all-atom MD simulations were built in CHARMM-GUI (Jo *et al*., 2008), whereas equilibration and production simulations were carried out with GROMACS (Abraham *et al*., 2015) on the RWTH Aachen University HPC system CLAIX-2023. In total, we built five systems (Supplemental File 3), all parameterized with the CHARMM36 force field (Lee *et al*., 2014). For the lower-confidence model 1705_02, we used the CHARMM-GUI Bilayer Builder (Wu *et al*., 2014) to generate a system with a membrane composed of sphingomyelin (18:1/24:0) lipids and 0.50 M calcium chloride. A transmembrane electric field corresponding to a 0.5 V potential drop across the periodic box length (Supplemental File 3) was applied to drive ion translocation during the simulation. We also applied positional restraints (1000 kJ mol⁻¹ nm⁻²) to the protein backbone and lipid headgroups, to stabilize the pore and membrane during the simulation. For the other systems, we used the Quick Bilayer tool (Park and Im, 2026) and selected a realistic plant plasma membrane composition (Pogozheva *et al*., 2022) supplemented with 0.15 M calcium chloride. In all cases, MLO proteins were oriented in the membrane using the PPM 2.0 server (Lomize *et al*., 2012). The temperature was set to 303.15 K, and hydrogen mass repartitioning as well as CHARMM minimization were enabled during system setup. Parameters of the resulting systems such as molecule number and box margins are provided in Supplemental File 3. The GROMACS input files were transferred to the cluster for equilibration and MD simulations.

All systems were energy-minimized in GROMACS (version 2024.4). Successful minimization was confirmed by monitoring the convergence of the potential energy and by inspection of the minimization log. Equilibration was subsequently carried out in six consecutive steps, comprising two NVT steps for 0.15 ns each followed by four NPT steps for 0.15, 1.00, 1.00, and 20.00 ns, while system restraints were gradually removed. Temperature coupling was carried out using the V-rescale thermostat (τ_t = 1.0 ps), and pressure coupling during the NPT steps was performed using the C-rescale barostat (τ_p = 5.0 ps) with semi-isotropic coupling. Temperature was monitored during NVT and NPT equilibration, and pressure was additionally monitored during the NPT steps.

Following equilibration, production MD simulations were carried out with a 2-fs time step for 25 or 50 million steps, corresponding to 50 or 100 ns per simulation, as indicated in Supplemental File 3. Temperature was maintained at 303.15 K using the V-rescale thermostat (τ_t = 1.0 ps; Bussi *et al*., 2007) with separate coupling groups for solute, membrane, and solvent. Pressure was controlled semi-isotropically at 1 bar using the C-rescale barostat (τ_p = 5.0 ps; Bernetti and Bussi, 2020). Long-range electrostatics were treated with the particle-mesh Ewald method (Darden *et al*., 1993) using a Coulomb cutoff of 1.2 nm. Van der Waals interactions were treated with the Verlet cutoff scheme and a force-switch modifier between 1.0 and 1.2 nm. Bonds involving hydrogen atoms were constrained using the LINCS algorithm (Hess *et al*., 1997). Position restraints, applied electric field conditions, and x–y pressure settings were varied between individual simulations (Supplemental File 3).

The analysis of the MD trajectories was performed using the GROMACS post-processing tools. Temperature, pressure, density, total energy, and box size along the membrane normal (Box-Z) were extracted from production energy files using gmx energy. Backbone RMSD was calculated with gmx rms. The z coordinates of individual Ca^2+^ and Cl^−^ ions over time were extracted with gmx traj to assess ion movement and potential pore-translocation events. For structural inspection, final production frames were extracted as .gro files using gmx trjconv. Data visualization was carried out by a custom Python script (version 3.10).

### Phylogenetic analysis of MLO proteins

For phylogenetic analysis of MLO proteins, we compiled a dataset of 351 MLO protein sequences from 25 species (Supplemental File 9). Sequence processing, alignment, tree inference, and visualization were carried out in R (version 4.4.3). Amino acid sequences were aligned with Clustal Omega using the msa package (version 1.36.0; Bodenhofer *et al*., 2015). The alignment was converted into phyDat format and used for maximum-likelihood phylogenetic inference with the phangorn package (version 2.11.1; Schliep, 2011). Maximum-likelihood distances were calculated under the JTT amino acid substitution model, and a neighbor-joining tree served as the starting topology for likelihood optimization. Branch lengths were optimized without additional topology rearrangement, and bootstrap support was estimated from 100 replicates. The final tree was visualized in daylight layout with clade assignments using the ggtree package (version 3.12.0; Yu *et al*., 2017).

### Generation of constructs for LCI and Y2H assays

Genes of interest were amplified from cDNA using appropriate oligonucleotides (Supplemental Table 2) or, in the case of *AtMLO7*, obtained as gBlock fragments (IDT, Coralville, Iowa, USA) with *att*B overhangs. PCR products were recombined into Gateway entry vectors (pDONOR201 and pDONOR207) by BP reaction and subsequently transferred into the respective destination vectors for LCI (pAM-PAT-CLuc-GWY, pAM-PAT-GWY-NLuc, pAM-PAT-GWY-NLuc; Huebbers *et al*., 2024) and Y2H assays (pGADT7, pGBKT7; Clonetech Laboratories, Mountain View, California, USA) by LR reactions according to the manufacturer’s instructions (Thermo Fisher Scientific, Waltham, Massachusetts, USA). Plasmids were introduced into *Escherichia coli* Top10 cells (Thermo Fisher Scientific) by heat-shock transformation. Plasmid DNA was isolated using the NucleoSpin Plasmid Mini kit (Macherey-Nagel, Düren, Germany). Plasmid integrity was verified by Sanger sequencing (Eurofins Scientific, Luxembourg, Luxembourg) using appropriate primers. Destination vectors were transformed for LCI assays into *Agrobacterium tumefaciens* GV3101 (Koncz and Schell, 1986) by heat-shock transformation or electroporation. For Y2H assays, *S. cerevisiae* AH109 (James *et al*., 1996) were transformed by the LiAc/PEG method (Gietz and Woods, 2002)

### Cultivation of *N. benthamiana* plants

*N. benthamiana* plants were cultivated in ED73 soil (Einheitserde Weksverband, Sinntal-Altengronau, Germany) in a Percival (Perry, Iowa, USA) growth cabinet (model SE-41ARLBS) with 23/20 °C average day/night temperature, a photoperiod of 16/8 h d^-1^ light/darkness, and a photosynthetic photon flux density of 92–162 *μ*mol m^-2^ s^-1^. After infiltration of *A. tumefaciens*, *N. benthamiana* plants were transferred to a growth chamber with a photoperiod of 16/8 h d^-1^ light/darkness with a photosynthetic photon flux density of 105–120 μmol m^-2^ s^-1^, an average day/night temperature of 23/20 °C and a relative humidity of 80% to 90%. Post-infiltration incubation was carried out for three days.

### Luciferase complementation imaging

LCI assays were carried out as described previously (Huebbers *et al*., 2024). Briefly, *N. benthamiana* leaves were infiltrated with *A. tumefaciens* cultures harboring the appropriate constructs for transient gene expression. Oligonucleotides used for the generation of constructs and Sanger sequencing are listed in (Supplemental Table 2). Before infiltration, *A. tumefaciens* cultures were grown overnight and resuspended in infiltration medium (10 mM MES pH 5.6; 10-mM MgCl_2_; 150 *µ*M acetosyringone). For co-infiltration, equal volumes of each *A. tumefaciens* strain were mixed and infiltrated into the abaxial side of fully expanded leaves of 4-week-old *N. benthamiana* plants. Luciferase complementation was assessed at three days after infiltration by spraying the transformed leaves of *N. benthamiana* with 1 mM D-luciferin (Revity, Waltham, Massachusetts, USA) Leaves were incubated in the dark for 10 min before luminescence was detected using a ChemiDoc XRS+ imaging system (BioRad, Feldkirchen, Germany). Luminescence intensities per unit area were quantified with the Image Lab software (BioRad, Feldkirchen, Germany). Luciferase data were normalized in three steps using a Python script. First, luminescence volume per area (VpA) was calculated as

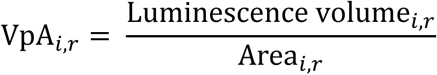

where *i* denotes the sample and *r* the replicate. Second, baseline correction was carried out separately for each replicate (e.g., each assessed leaf) using the 5% quantile of the corresponding VpA values:

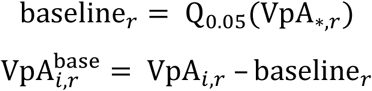

Third, baseline-corrected values were normalized to the *At*CAM2 reference sample within each replicate:

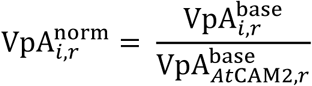

We used an *R* script for statistics and luminescence data visualization. Data processing was carried out with the dplyr package and violin plots and heat maps were generated with ggplot2, both from the tidyverse collection (Wickham *et al*., 2019). The distribution of normalized relative luminescence values was assessed by quantile–quantile plots and Shapiro–Wilk tests (*α* = 0.05; Shapiro and Wilk, 1965). Homoscedasticity was evaluated using Levene’s test (*α* = 0.05; Gastwirth *et al*., 2009). Normally distributed data with homogeneous variances were analyzed using Student’s *t*-test (Student, 1908), whereas non-normally distributed samples were compared using the Wilcoxon–Mann–Whitney test (Mann and Whitney, 1947). We used false discovery rate (FDR) correction (FDR; Benjamini and Hochberg, 1995) to adjust *p* values for multiple testing.

### Yeast two-hybrid assays

Y2H assays were carried out as described elsewhere (Huebbers *et al*., 2024). Briefly, *S. cerevisiae* cells accumulating Gal4AD*-At*EXO70D3 or Gal4AD*-At*EXO70H4 (Huebbers *et al*., 2024) were transformed (see Generation of constructs for LCI and Y2H assays) with genes encoding the IC1, IC2, IC3, or CT fragments of the MLO proteins of interest. Yeast transformants were spotted on a medium lacking leucine and tryptophan for growth control, and on a medium lacking leucine, tryptophan, and histidine to detect putative interactions. For these tests, yeast cells were grown overnight in liquid synthetic complete medium (James *et al*., 1996). The cells were harvested by centrifugation at 3,000 × *g* for 1 min at room temperature and washed with sterile water before adjusting the OD_600_ of the solutions to 1.0. Ten-fold dilution series were established over 4 orders of magnitude and 3 *µ*L per dilution and construct combination were spotted onto the aforementioned media. Yeast growth was documented after three days.

### Protein extraction, SDS-PAGE, and immunoblot analysis

To extract yeast total protein, yeast cells were cultured to an OD_600_ of 0.6 and cells were harvested by centrifugation at 3000*g* for 1 min at room temperature. The supernatant was discarded, and the cells were transferred to fresh 1.5-mL reaction tubes. On ice, 300 μL trichloroacetic acid buffer (25.0 mM ammonium acetate; 1.0 mM EDTA; 100.0 mL L^-1^ tricholoroacetic acid; 10.0 mM Tris-HCl, pH 8.0) and about 10 glass beads (0.7–1.0 mm) were added to the cell pellet. The samples were mixed five times for 1 min on a benchtop mixer (Vibrax VXR, IKA) with a 3 min incubation period on ice between each mixing step. The cell lysates were transferred to fresh chilled 1.5-mL reaction tubes. The remaining glass beads were washed with 100 μL trichloroacetic acid buffer and the wash solution was pooled with the cell lysate. Lysates were centrifuged at 16,000*g* for 10 min at and 4 °C, supernatants were discarded, and the pellets were resuspended in 150 μL resuspension solution (100.0 mL L^-1^ 20% SDS solution (m/v); 100.0 mM Tris, pH 11.0). After incubation for 5 min at 99 °C, the samples were cooled to room temperature and centrifuged for 16,000*g* for 30 s. Eventually, 120 μL supernatant was transferred to a fresh 1.5-mL reaction tube. Protein concentrations within the extracts were determined by the BCA method (Smith *et al*., 1985) using a Pierce BCA protein assay kit (Thermo Fisher Scientific).

SDS-PAGE and immunoblot detection of Gal4DBD fusion proteins was carried out as described previously (Huebbers *et al*., 2024). Briefly, protein samples were resuspended in LDS Sample buffer and incubated at 95 °C for 5 min. Subsequently, 5 µg of total protein was separated by SDS-PAGE, transferred to a nitrocellulose membrane (Carl Roth, Karlsruhe, Germany), and subjected to immunoblot analysis. A mouse α-Gal4DBD antibody (Santa Cruz Biotechnology, Dallas, Texas, USA; product number: sc-510) was used as primary antibody, while a goat α-mouse antibody coupled to horseradish peroxidase (Thermo Fisher Scientific; product number: 32430) served as secondary antibody. Antigen–antibody complexes were detected by chemiluminescence using SuperSignal West Femto Western substrate (Thermo Fisher Scientific). As a loading control, the nitrocellulose membrane was stained with a Ponceau S solution (AppliChem, Darmstadt, Germany; 5% acetic acid, 0.5% (m/v) Ponceau S).

## Acknowledgments

We thank Lana Kozmer for contributing to *At*MLO9–*At*EXO70 LCI assays, Sarah Züscher for supporting the work on *At*MLO4 protein–protein interaction assays, Jianing Li for supporting the acquisition of *Hv*Mlo oligomerization LCI data, and Anja Reinstädler for keeping the lab running even under severe circumstances. We apologize for not mentioning or citing relevant work due to space limitations.

## Author contributions

**JWH** designed and generated expression constructs, carried out LCI experiments and analyzed the resulting data, developed and used the AF3 pipeline, developed and used the MOLE pipeline, developed ChimeraX scripts, carried out MD simulations, performed phylogenetic analyses of MLO proteins, designed the figures, performed statistical analyses, conceived the study, and drafted and edited the manuscript.

**CKD** contributed to the design of the MD simulations, guided the setup and optimization of the MD simulations, contributed substantially to discussions of the computational aspects, and discussed the organization of the manuscript.

**AS** generated expression constructs, conducted LCI and Y2H experiments for EXO70-MLO interactions, and carried out immunoblotting.

**MEF** designed and generated expression constructs and conducted LCI and Y2H experiments for EXO70-MLO interactions.

**HS** generated expression constructs and conducted LCI and Y2H experiments for EXO70-MLO interactions.

**ML** generated expression constructs and conducted LCI experiments for EXO70-MLO interactions.

**SCJL** designed and generated expression constructs for LCI and Y2H experiments of EXO70 and MLO proteins and contributed to MLO-CML12 interaction studies.

**MFr** designed and generated expression constructs and detected *Hv*Mlo oligomerization by LCI experiments.

**MFy** contributed to the initial discussions on the concept and computational aspects of this work and reviewed and revised the manuscript.

**RP** contributed to the conceptual framework for the study, discussed the data, reviewed and revised the manuscript, and arranged research funding and key resources.

## Data availability statement

All relevant data are available within the manuscript and its supporting materials. All code used in this study is publicly available on <u>GitHub</u>.

## Supplemental Materials

### Supplemental Figures

**Supplemental Figure 1:**
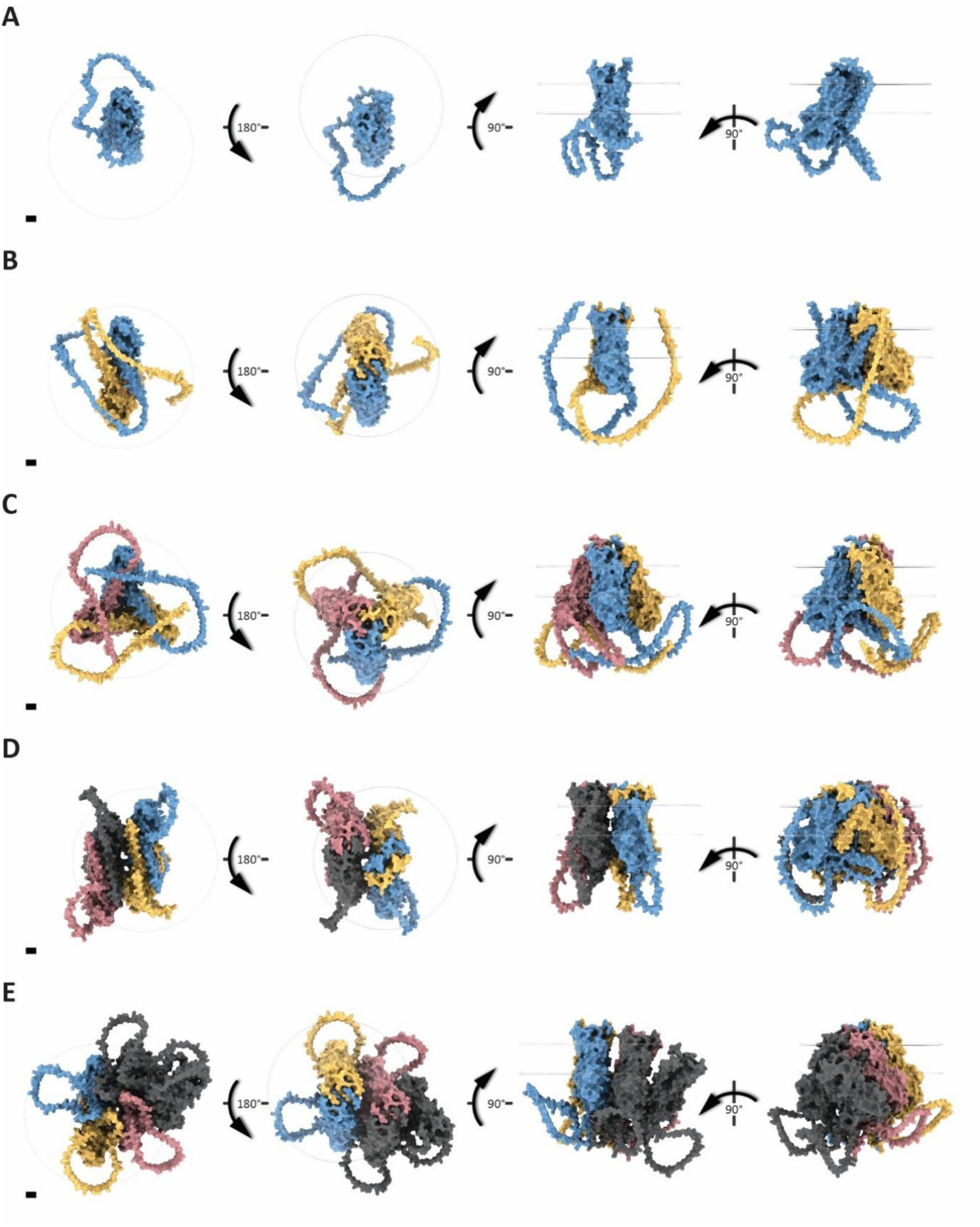
Oligomeric state modulates subunit arrangement of AF3 predictions for *Hv*Mlo. AF3 models (seed 1710, model 0) are shown for the *Hv*Mlo monomer (**A**) and homo-oligomers from dimer to apentamer (**B–E**). Depending on prediction accuracy, models may appear differently in runs associated with other seeds. *Hv*Mlo subunits are color-coded: A, blue; B, yellow; C, red; others, gray. Transparent discs indicate the plasma membrane. Perspectives from left to right show the cytosolic (bottom) view, extracellular/luminal (top) view, and two side views (subunit C in the back). Scale bars (bottom left) indicate 10.0 Å.

**Supplemental Figure 2:**
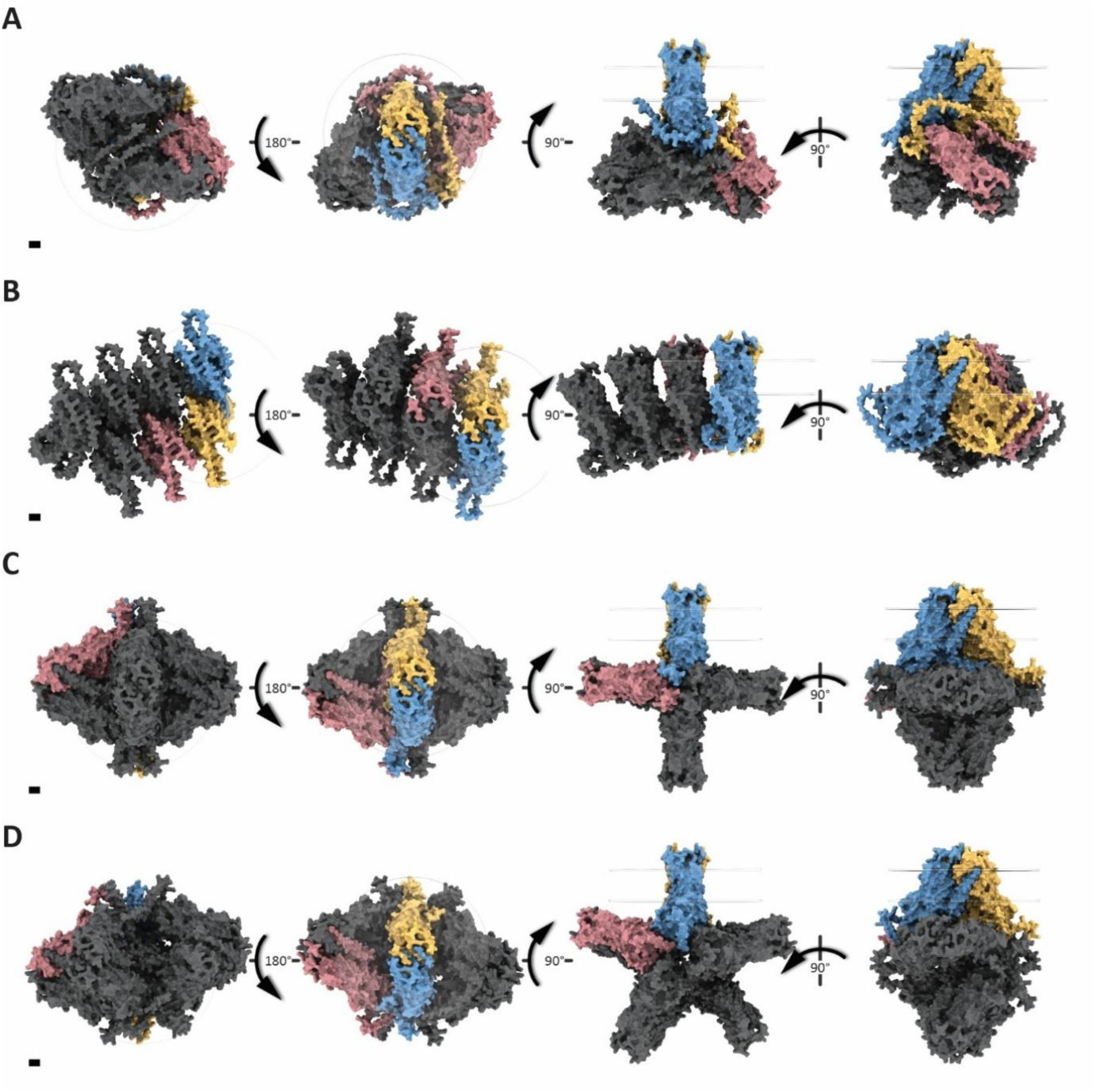
Oligomeric state modulates subunit arrangement of AF3 predictions for *Hv*Mlo. Representative AF3 models (seed 1710, model 0) are shown for *Hv*Mlo homo-oligomers from hexamer to nonamer (**A–D**). Depending on prediction accuracy, models may appear differently in runs associated with other seeds. *Hv*Mlo subunits are color-coded: A, blue; B, yellow; C, red; others, gray. Transparent discs indicate the plasma membrane. Perspectives from left to right show the cytosolic (bottom) view, extracellular/luminal (top) view, and two side views (subunit C in the back). Scale bars (bottom left) indicate 10.0 Å.

**Supplemental Figure 3:**
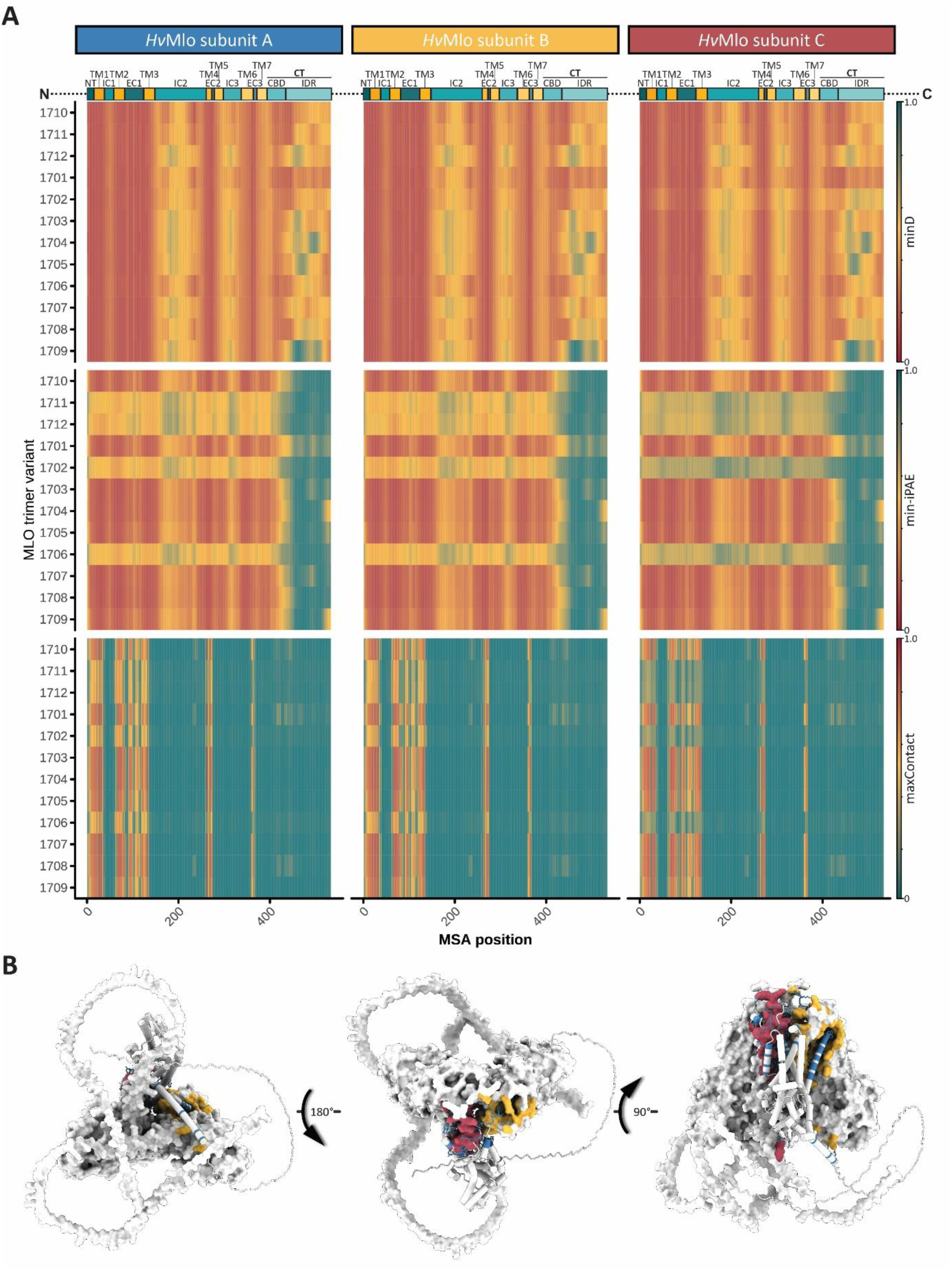
*Hv*Mlo transmembrane domains and extracellular/luminal loops mediate trimerization. **A)** AF3 per-residue metrics highlight *Hv*Mlo trimer interfaces. Heatmaps show average minD, min-iPAE, and maxContact values across the three subunits of the *Hv*Mlo trimer for 12 AF3 runs, predicting five models each. The color scale indicates the respective metric values as shown. The *Hv*Mlo domain architecture is depicted above the heatmaps. **NT**: extracellular/luminal N-terminal domain; **TM**: transmembrane domain; **IC**: intracellular loop; **EC**: extracellular/luminal loop; **CT**: intracellular C-terminal domain; **CBD**: CAM-binding domain; **IDR**: intrinsically disordered region **B)** Representative AF3 prediction of the *Hv*Mlo trimer (seed 1710, model 0). Interface residues in subunit A are highlighted (blue) and side chains are shown in stick representation. Subunits B and C are shown as surfaces; residues within 4 Å of the highlighted subunit A residues are colored yellow (subunit B) or red (subunit C). Perspectives from left to right show the cytosolic (bottom) view, extracellular/luminal (top) view, and a side view (subunit A in the front).

**Supplemental Figure 4:**
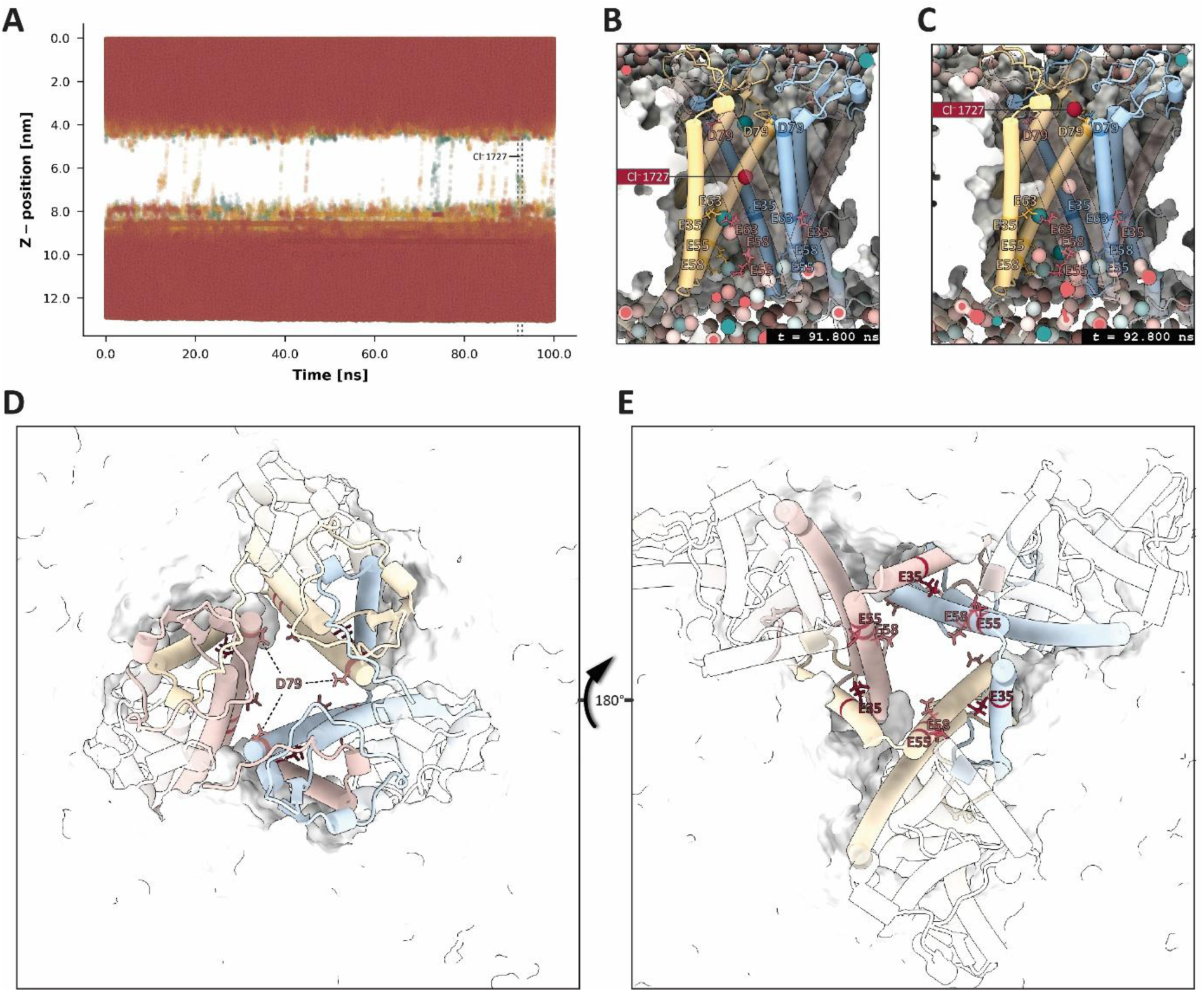
MD simulations indicate ion permeability of *Hv*Mlo trimers. **A)** Cl^-^ trajectories along the membrane normal (z-axis) in an all-atom MD simulation initiated from the *Hv*MloΔIDR lower-confidence model 1705_02. Dashed lines indicate the time points shown in **B** (91.800 ns) and **C** (92.800 ns). **B, C**) Close-up views of three-dimensional models corresponding to two frames from the simulation shown in **A** (t = 91.800 ns, **B**; t = 92.800 ns, **C**). Non-transparent cartoons of residues 1–160 of subunits A (blue) and B (yellow) are shown. Side chains of pore-lining anionic residues are displayed as sticks for all subunits (with subunit C in front) and labeled using the one-letter amino acid code. Cl^-^ ions are shown in red and Ca^2+^ ions in petrol. The membrane is shown in cross-section. **D, E)** Extracellular/luminal (top; **D**) and cytosolic (bottom; **E**) views of *Hv*MloΔIDR 1705_02 embedded in a sphingomyelin (18:1/24:0) membrane. Residues 1–160 of each subunit are colored (A, blue; B, yellow; C, red), and pore-lining anionic residues are shown as red sticks.

**Supplemental Figure 5:**
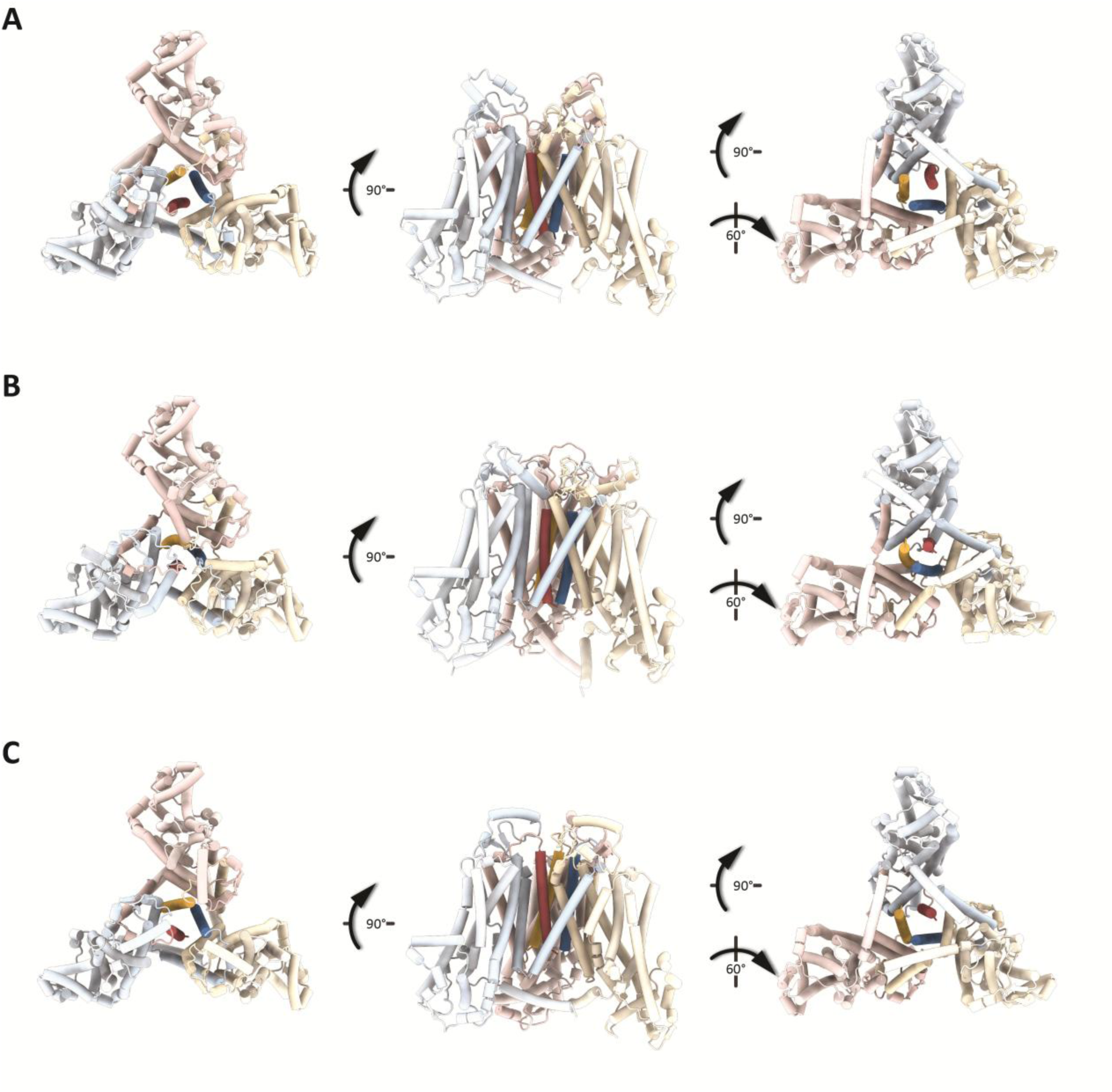
AF3 predicts the N-termini of *At*MLO7, -8, and -10 as additional transmembrane domains. **A-C)** Top-ranked models for the predictions of *At*MLO7ΔIDR (**A**; seed 1712, model 0), *At*MLO8ΔIDR (**B**; seed 1711, model 0), and *At*MLO10ΔIDR (**C**; seed 1707, model 0). Perspectives from left to right show the extracellular/luminal (top) view, a side view (subunit C in the back), and the cytosolic (bottom) view. The N-terminal membrane-spanning domains are highlighted.

**Supplemental Figure 6:**
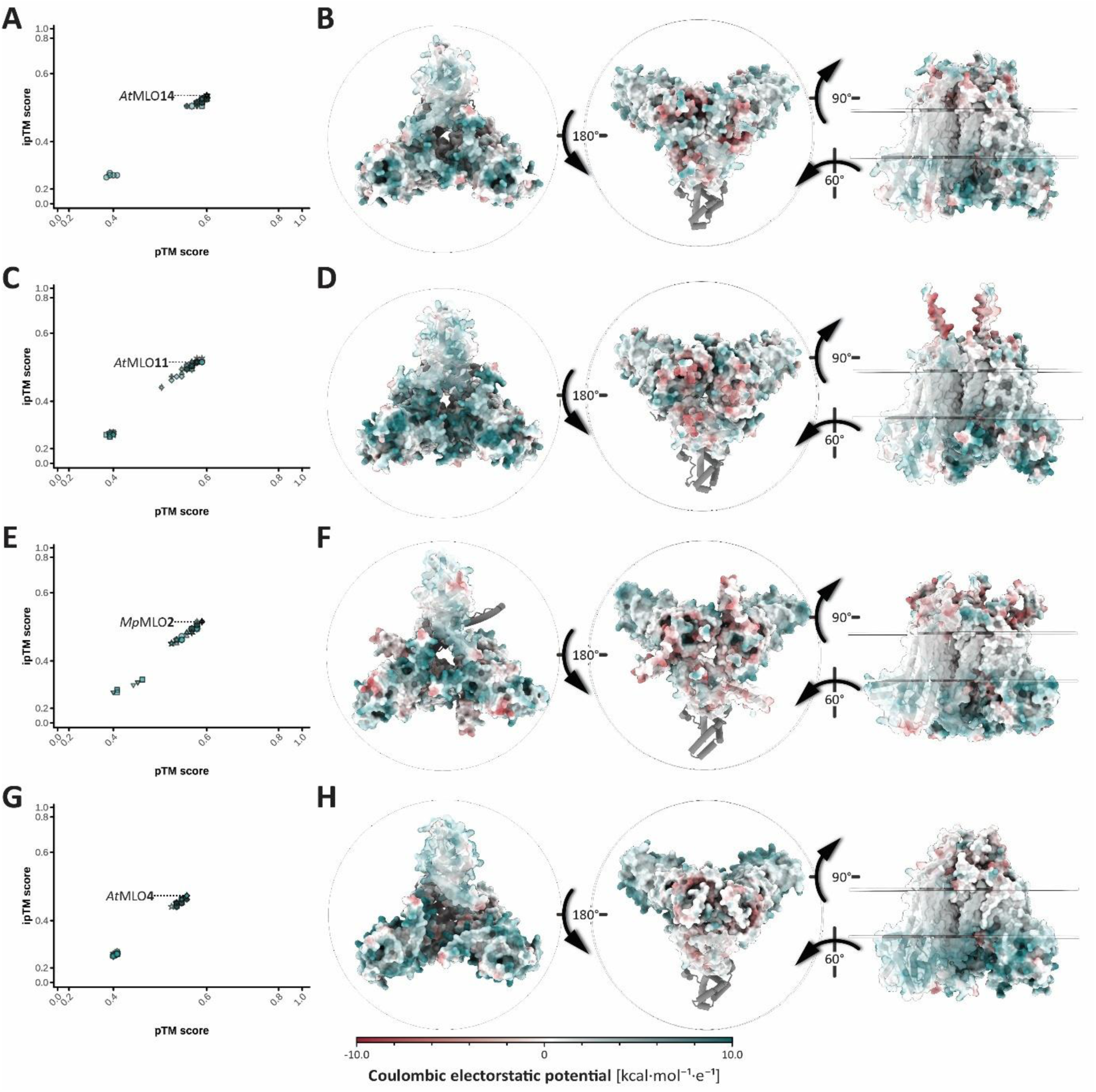
AF3 prediction of Clade I MLOΔIDR trimers. **A, C, E, G)** Joint distribution of pTM and ipTM scores for individual MLOΔIDR trimer models. For each MLO isoform, the model with the highest AF3 ranking score is labeled. Axes were transformed using sigmoidal (logistic) functions to improve the visibility of data points. **B, D, F, H)** Representative AF3 predictions corresponding to the labeled models in the plots to the left. Molecular surfaces are colored by Coulombic electrostatic potential as indicated by the color scale. The surface of subunit A is shown with 40% transparency. Transparent discs indicate the plasma membrane. Perspectives from left to right show the cytosolic (bottom) view, extracellular/luminal (top) view, and a side view (subunit C in the back).

**Supplemental Figure 7:**
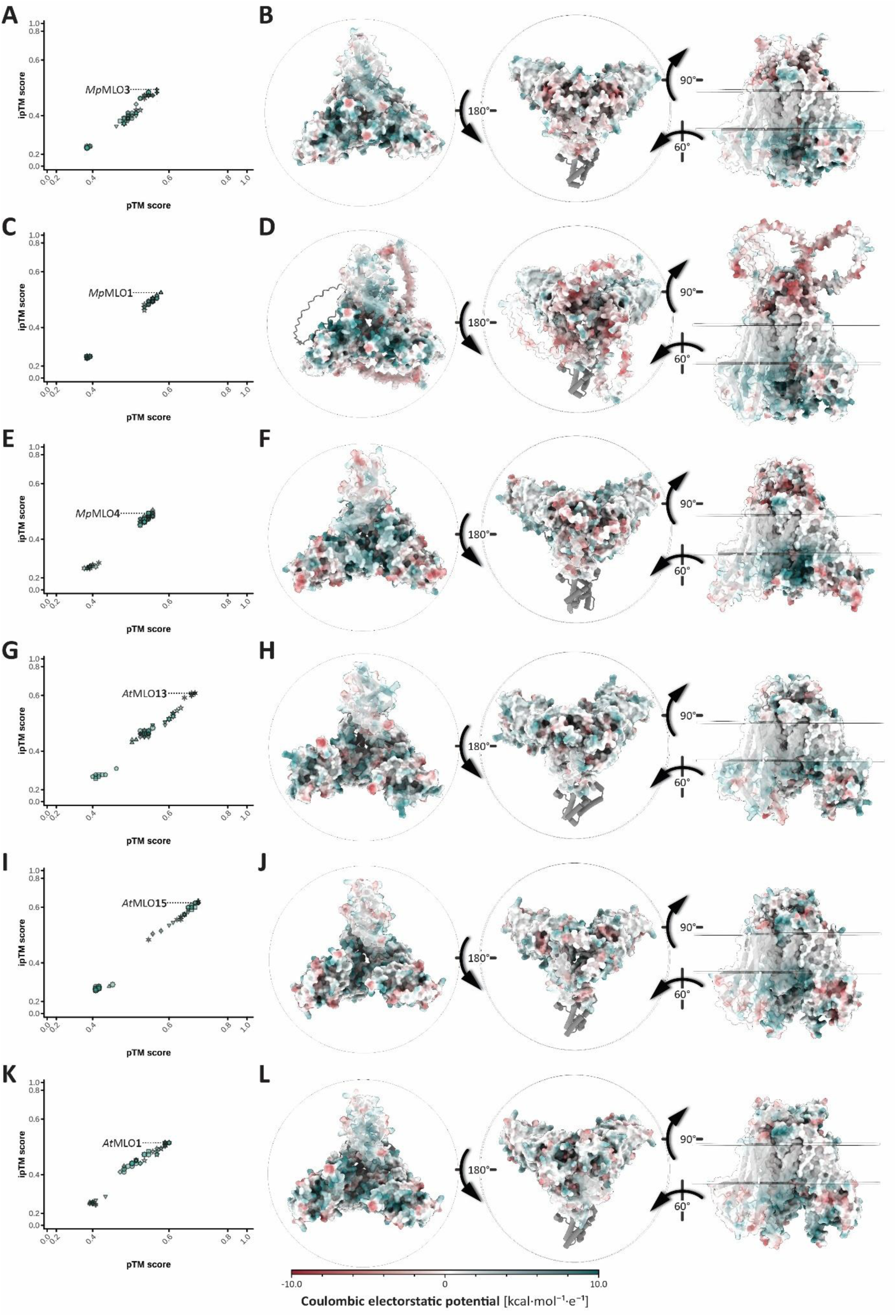
AF3 prediction of Clade II MLOΔIDR trimers. **A, C, E, G)** Joint distribution of pTM and ipTM scores for individual MLOΔIDR trimer models. For each MLO isoform, the model with the highest AF3 ranking score is labeled. Axes were transformed using sigmoidal (logistic) functions to improve the visibility of data points. **B, D, F, H)** Representative AF3 predictions corresponding to the labeled models in the plots to the left. Molecular surfaces are colored by Coulombic electrostatic potential as indicated by the color scale. The surface of subunit A is shown with 40% transparency. Transparent discs indicate the plasma membrane. Perspectives from left to right show the cytosolic (bottom) view, extracellular/luminal (top) view, and a side view (subunit C in the back).

**Supplemental Figure 8:**
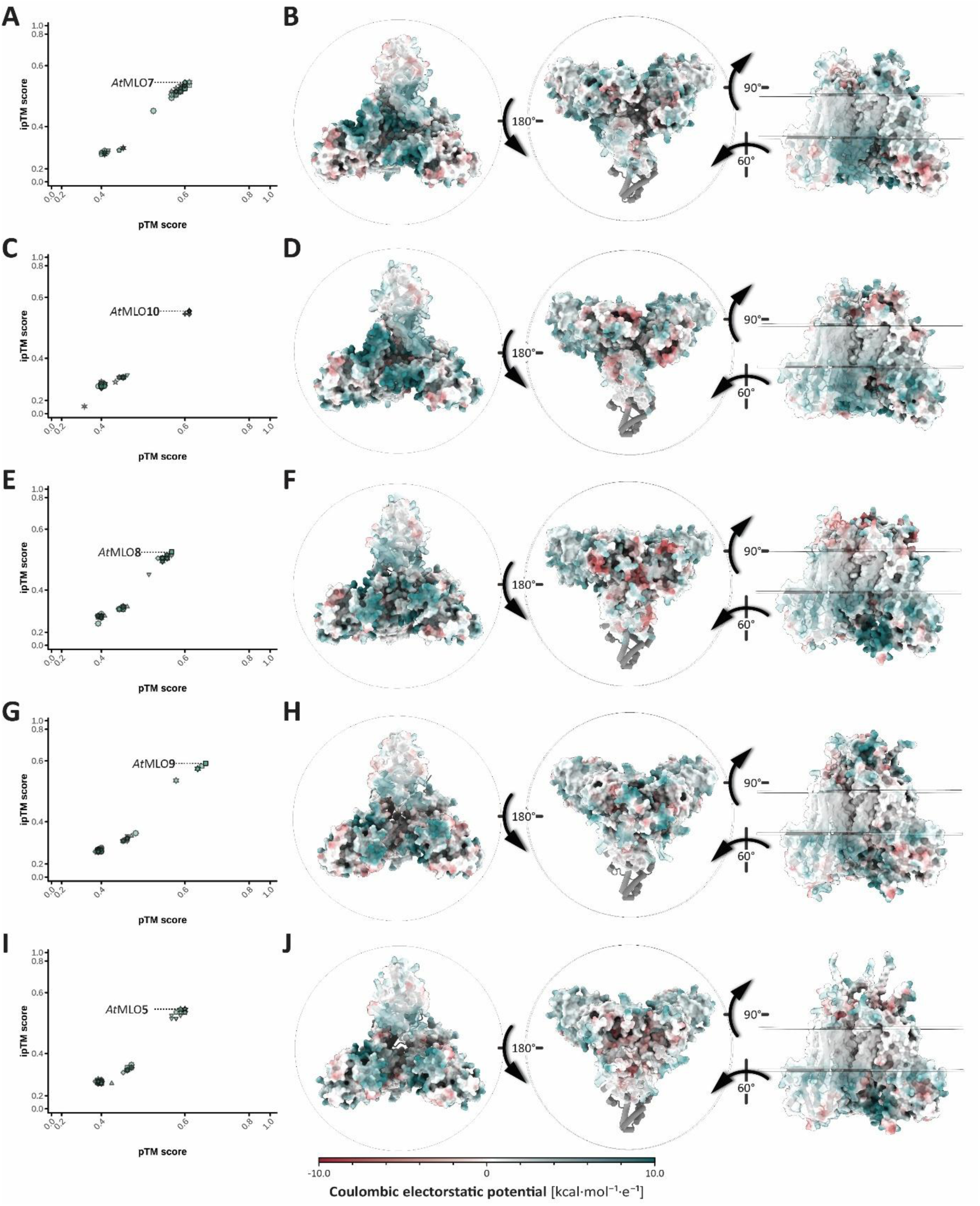
AF3 prediction of Clade III MLOΔIDR trimers. **A, C, E, G)** Joint distribution of pTM and ipTM scores for individual MLOΔIDR trimer models. For each MLO isoform, the model with the highest AF3 ranking score is labeled. Axes were transformed using sigmoidal (logistic) functions to improve the visibility of data points. **B, D, F, H)** Representative AF3 predictions corresponding to the labeled models in the plots to the left. Molecular surfaces are colored by Coulombic electrostatic potential as indicated by the color scale. The surface of subunit A is shown with 40% transparency. Transparent discs indicate the plasma membrane. Perspectives from left to right show the cytosolic (bottom) view, extracellular/luminal (top) view, and a side view (subunit C in the back).

**Supplemental Figure 9:**
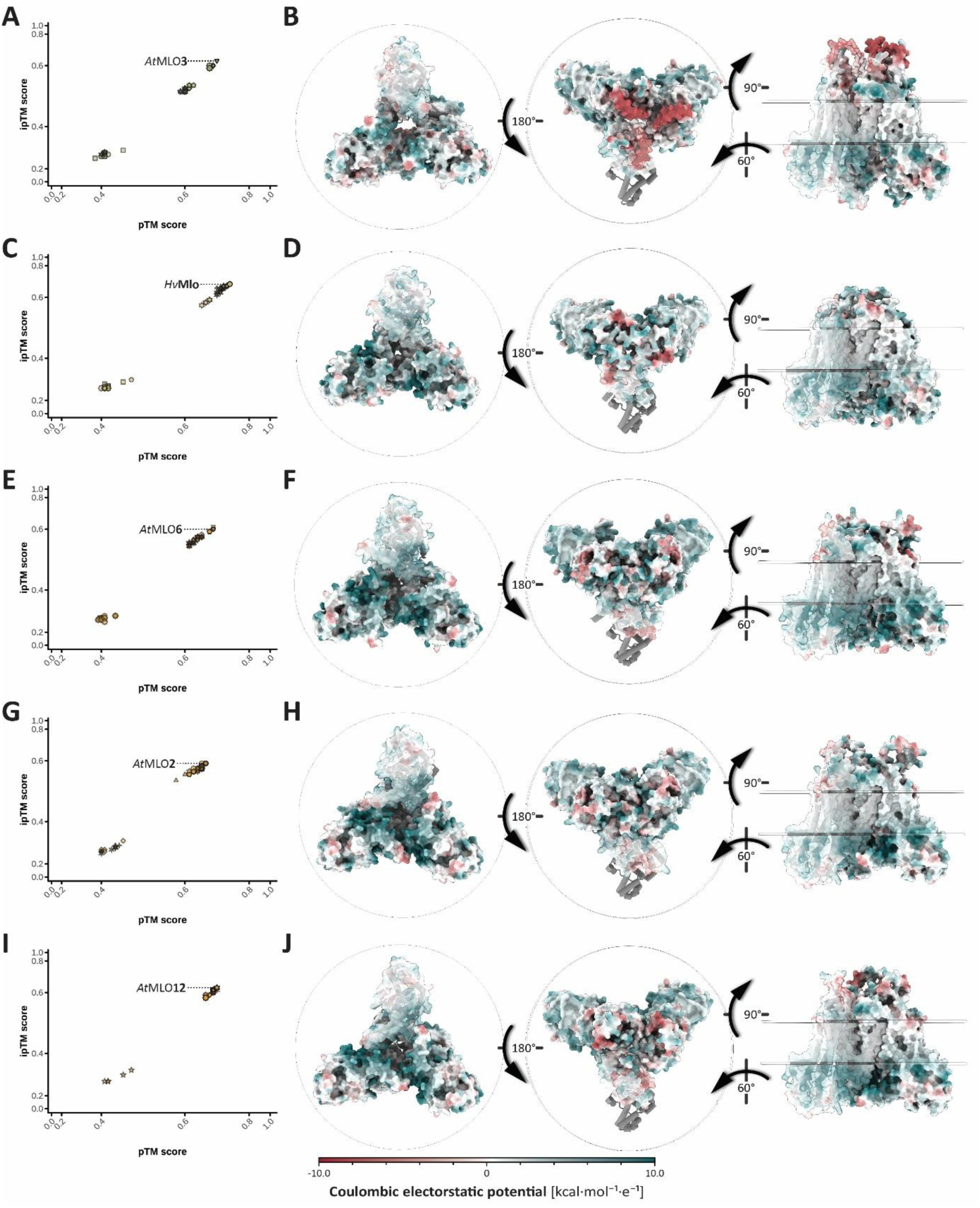
AF3 prediction of Clade IV, V, and VI MLOΔIDR trimers. **A, C, E, G)** Joint distribution of pTM and ipTM scores for individual MLOΔIDR trimer models. For each MLO isoform, the model with the highest AF3 ranking score is labeled. Axes were transformed using sigmoidal (logistic) functions to improve the visibility of data points. **B, D, F, H)** Representative AF3 predictions corresponding to the labeled models in the plots to the left. Molecular surfaces are colored by Coulombic electrostatic potential as indicated by the color scale. The surface of subunit A is shown with 40% transparency. Transparent discs indicate the plasma membrane. Perspectives from left to right show the cytosolic (bottom) view, extracellular/luminal (top) view, and a side view (subunit C in the back).

**Supplemental Figure 10:**
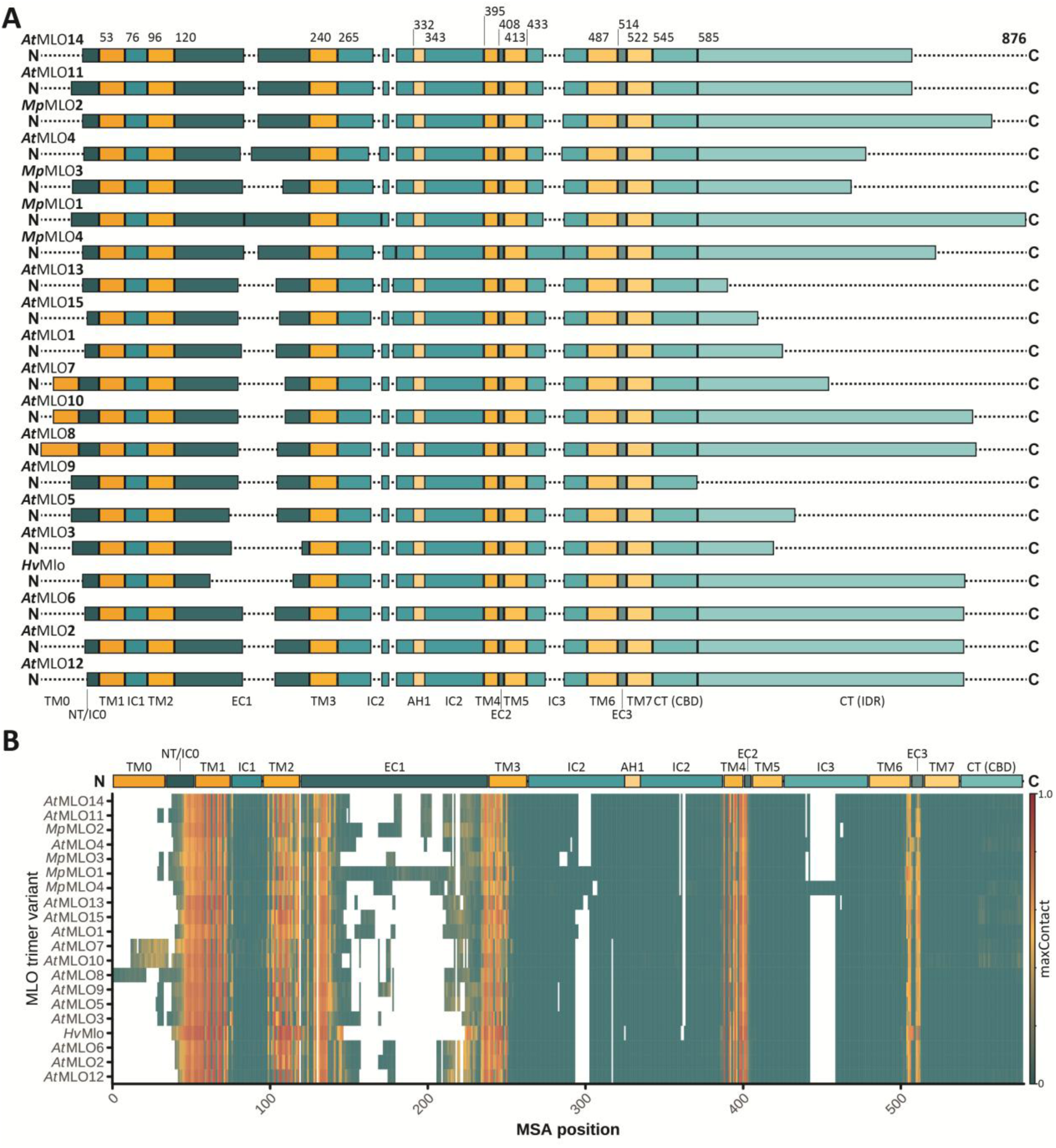
Domain architecture and trimer interface residues are conserved across *At*MLO isoforms, *Mp*MLO isoforms, and *Hv*Mlo. **A)** Domain architecture of *At*MLO and *Mp*MLO isoforms, together with *Hv*Mlo. Proteins are ordered according to their phylogeny. Plain numbers indicate the first residue of each domain, whereas the total sequence length (876) is shown in bold. **NT**: extracellular/luminal N-terminal domain; **TM**: transmembrane domain; **IC**: intracellular loop; **AH**: amphipathic helix; **EC**: extracellular/luminal loop; **CT**: intracellular C-terminal domain; **CBD**: CAM-binding domain; **IDR**: intrinsically disordered region. **B)** Average maxContact scores across the five models of the highest-ranked prediction run (Figure 3C; Supplemental File 4) for each tested MLOΔIDR trimer variant mapped on an MSA of the listed MLO isoforms. The color scale indicates maxContact values as shown. A consensus MLO domain architecture is shown above the heatmap.

**Supplemental Figure 11:**
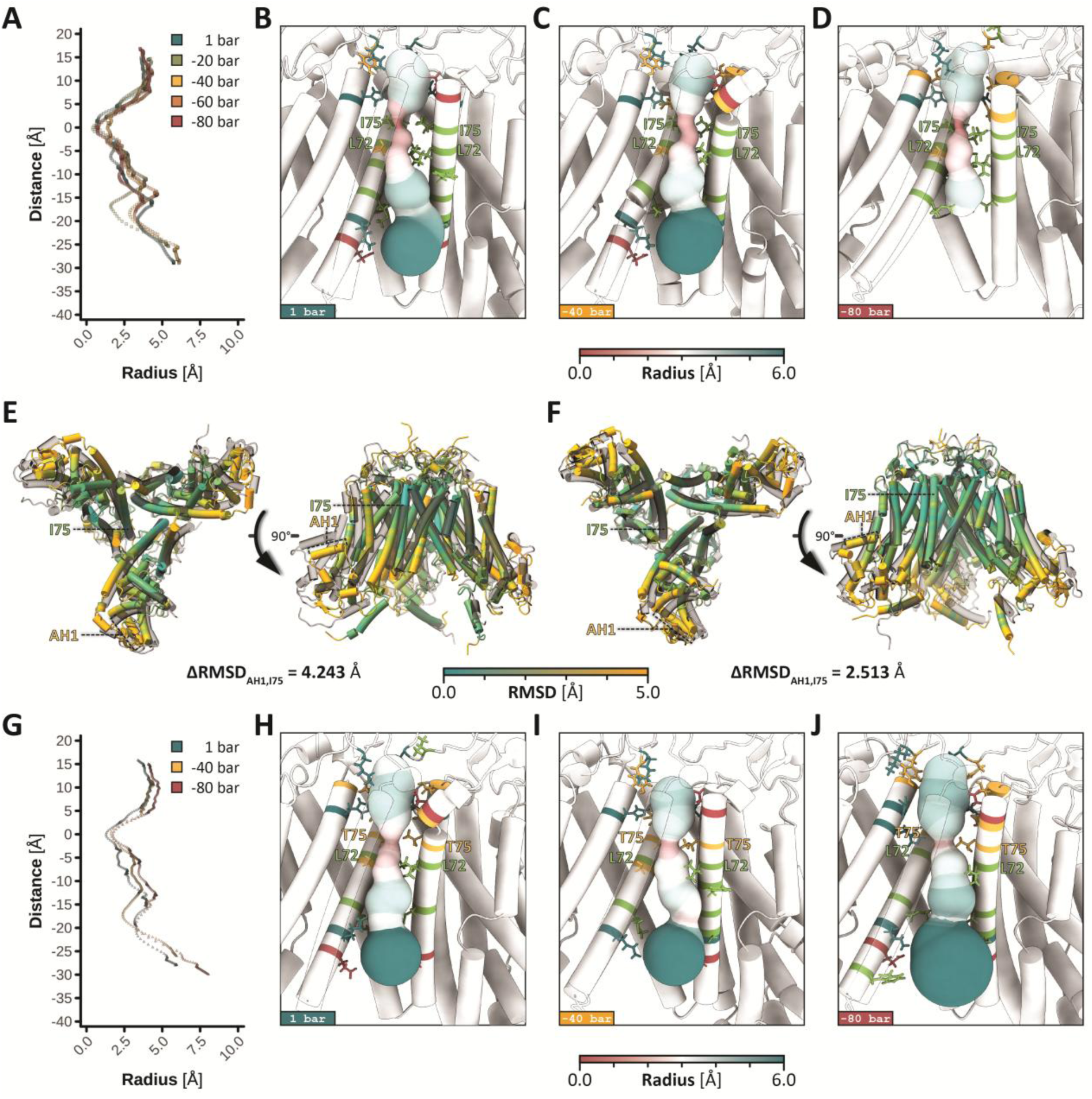
Expansion of *Hv*MloΔIDR variants during MD simulations with simulated lateral tension. **A, G)** Pore-radius profiles of native *Hv*MloΔIDR (seed 1710, model 0; **A**) and *Hv*MloΔIDR I75T carrying extracellular disulfide bonds (C86–C114, C98–C367; **G**). Pore descriptors for the last frame of each 50 ns MD run were obtained *via* the MOLE API. Point colors indicate the x–y pressures during MD simulations. **B–D, H–J)** Three-dimensional “balloon” representations of transmembrane pores generated from MOLE output. To improve pore visibility, residues 1–50 of subunit B (left) are shown with reduced opacity and subunit C is omitted. Pore-lining residues were identified in ChimeraX (*interface select*) and are shown as sticks, colored by polarity (nonpolar, green; polar, yellow; basic, petrol; acidic, red). Bottleneck residues are labeled. **E, F)** Structural superposition of *Hv*MloΔIDR trimer models from the last frames of 1 bar and –80 bar x-y pressure MD simulations, without (**E**) or with (**F**) disulfide bridges between conserved extracellular cysteines (Figure 4A, B). Per-residue RMSD is mapped onto the 1 bar models, while the –80 bar models are displayed as transparent overlays. Perspectives (left to right): cytosolic (bottom) view and side view (subunit C in the back). The pore-constricting residue I75 and the putative force-transmitting helix AH1 are labeled, and the difference between their average backbone RMSD values (ΔRMSD) is indicated.

**Supplemental Figure 12:**
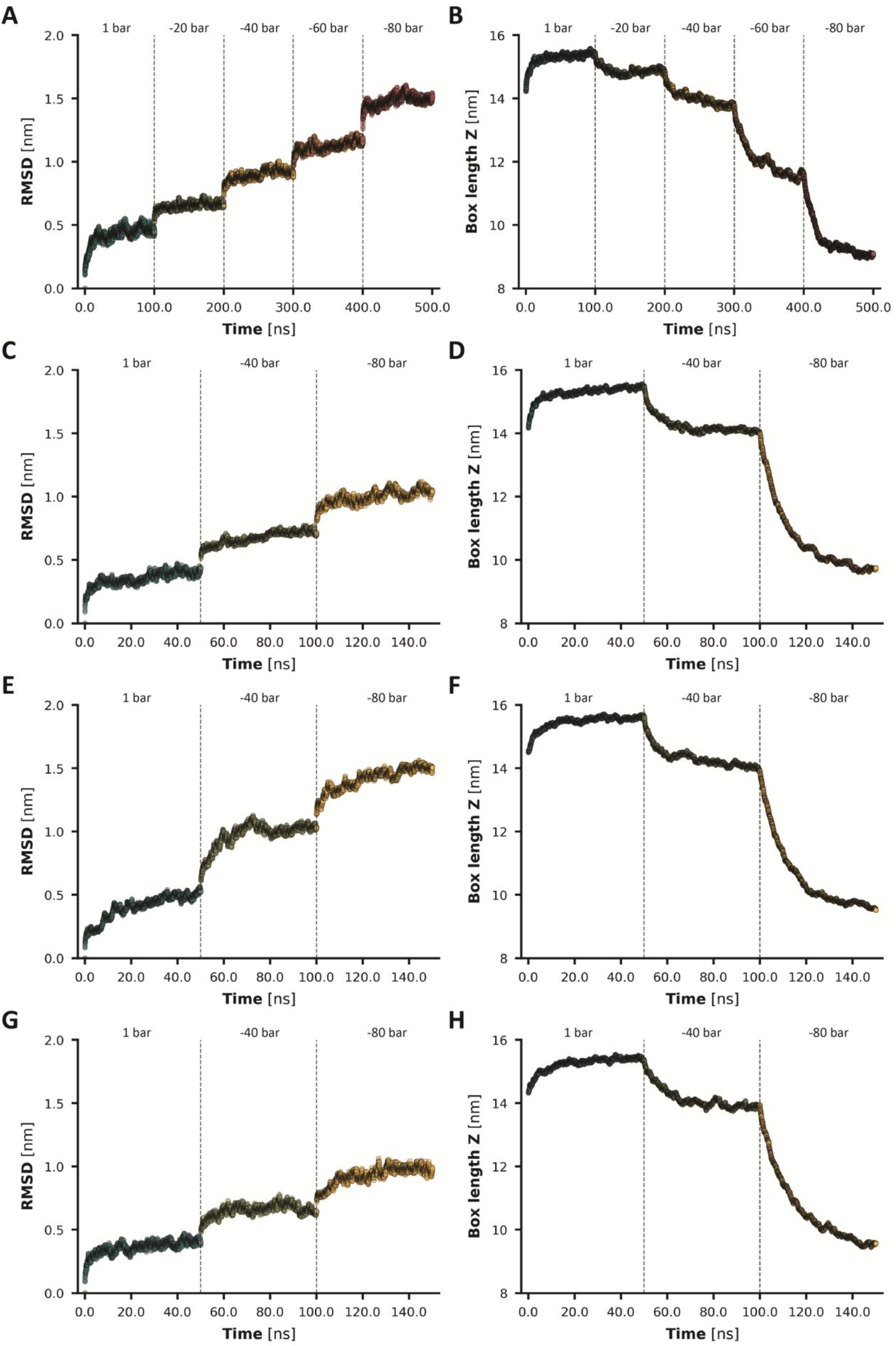
Backbone RMSD and box z-dimensions during MD simulations under progressively decreasing x–y pressure. **A, C, E, G)** Backbone RMSD traces over the simulation time, computed in GROMACS (gmx rms). **B, D, F, H)** Box z-dimensions over the simulation time, extracted in GROMACS (*gmx energy*). RMSD traces and box z-dimension are shown for the *Hv*MloΔIDR trimer (seed 1710, model 0) without (**A**, **B**) and with C86–C114 and C98–C367 disulfide bonds (**C**, **D**), the *At*MLO2ΔIDR trimer (seed 1710, model 0) with disulfide bonds between the corresponding residues (**E**, **F**), and the *Hv*MloΔIDR trimer with an I75T mutation and disulfide bonds (**G**, **H**).

**Supplemental Figure 13:**
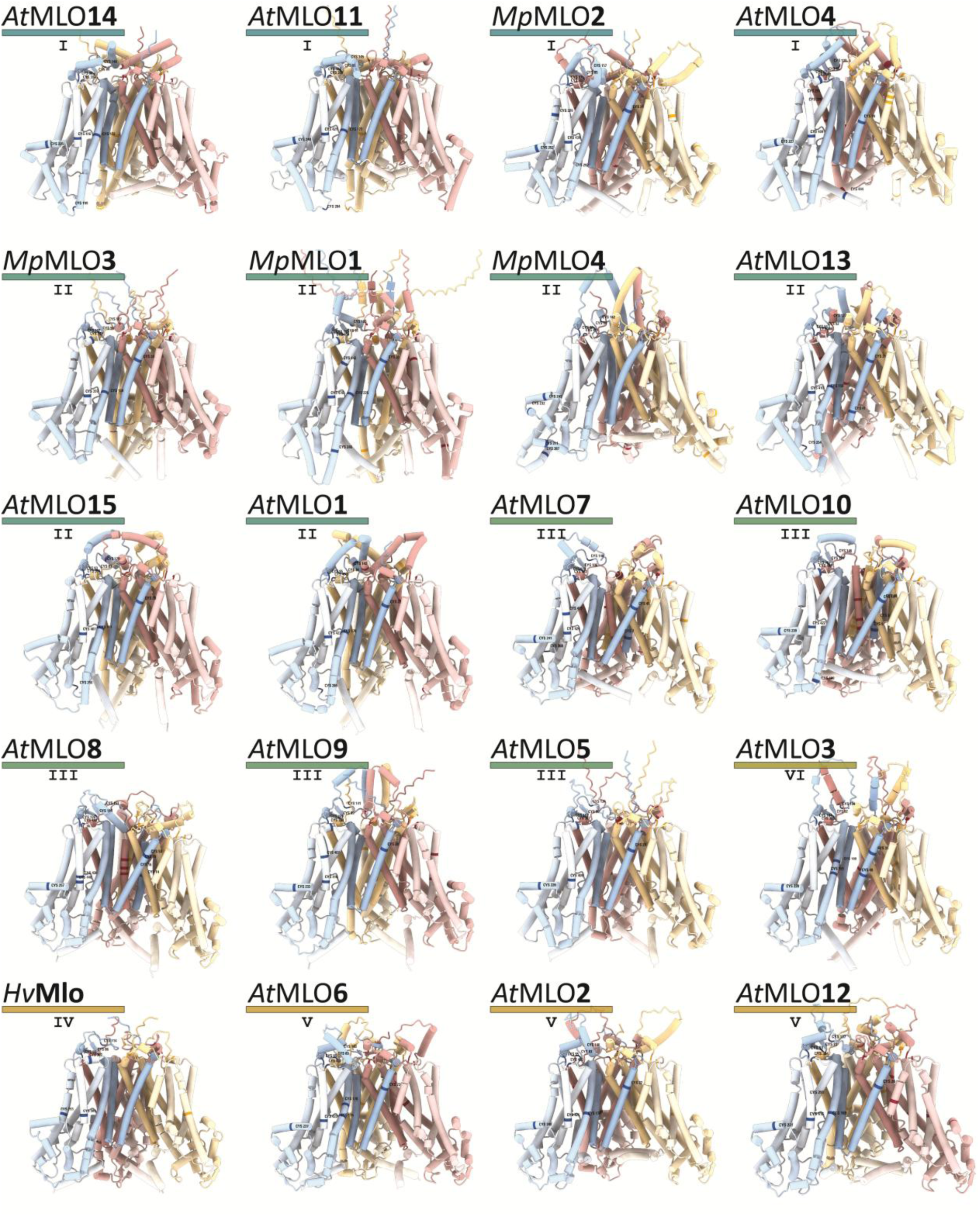
Cysteine positions in top-ranked MLOΔIDR trimer models. AF3-predicted top-ranked MLOΔIDR trimers are shown with color-coded subunits (A, blue; B, yellow; C, red). Cysteine residues are highlighted, and their positions are labeled for subunit A. Roman numerals indicate phylogenetic clades.

**Supplemental Figure 14:**
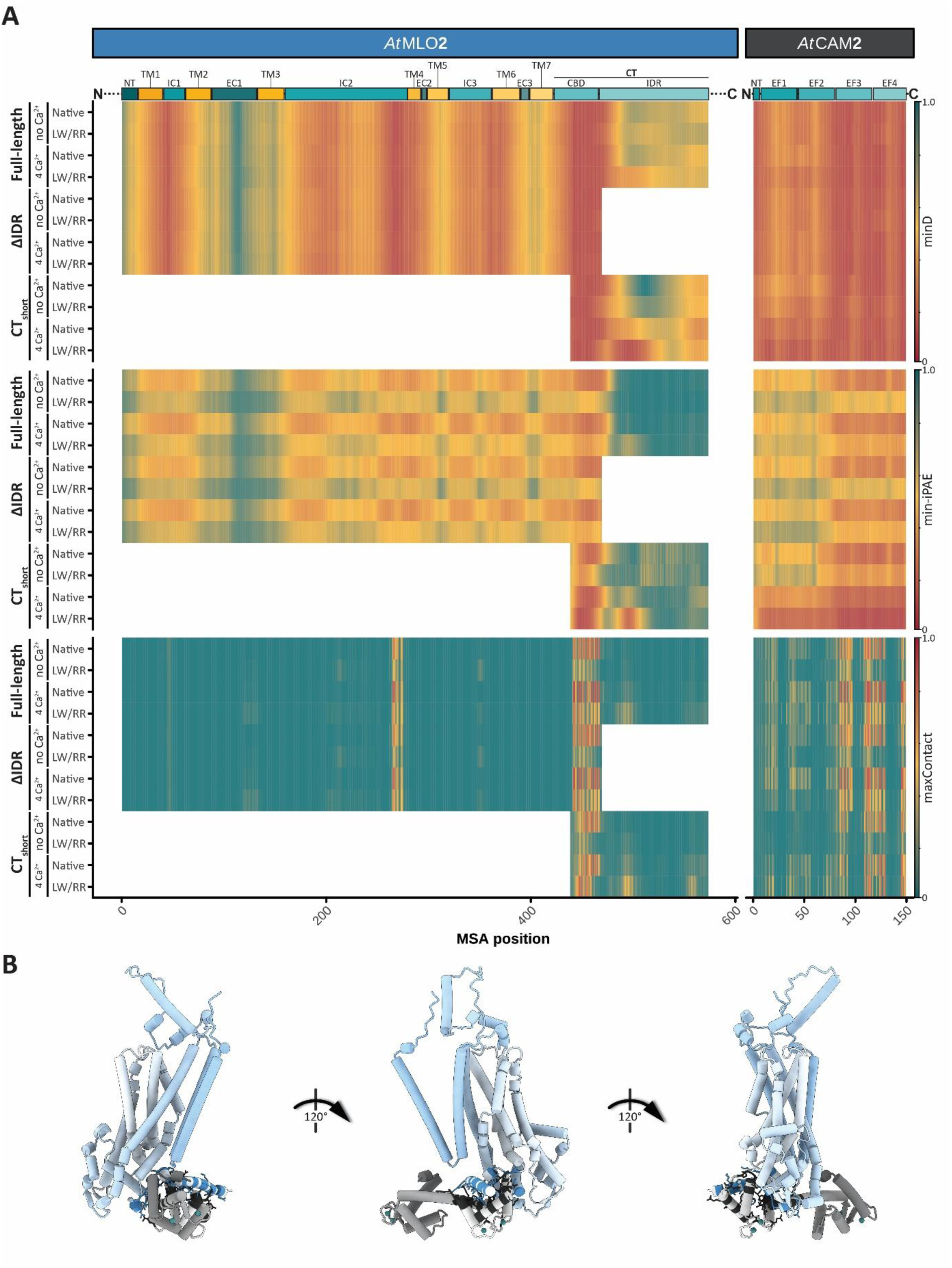
*At*CAM2 binds to the CBD and the IC2 of *At*MLO2 in AF3 predictions. **A)** AF3 per-residue metrics highlight interfaces in *At*MLO2–*At*CAM2 heterodimer predictions. Heatmaps show the average minD, min-iPAE, and maxContact scores over 12 AF3 runs (one per seed), with five models per run. The color scale indicates the respective metric values as shown. The *At*MLO2 and *At*CAM2 domain architectures are shown above the heatmaps. **NT**: extracellular/luminal N-terminal domain; **TM**: transmembrane domain; **IC**: intracellular loop; **AH**: amphipathic helix; **EC**: extracellular/luminal loop; **CT**: intracellular C-terminal domain; **CBD**: CAM-binding domain; **IDR**: intrinsically disordered region; **EF**: EF-hand-containing domain **B)** Representative AF3 prediction of the *At*MLO2-*At*CAM2 hetero-dimer (seed 1705, model 0). Interface residues are highlighted (*At*MLO2: blue; *At*CAM2: gray) with their side chains shown as sticks.

**Supplemental Figure 15:**
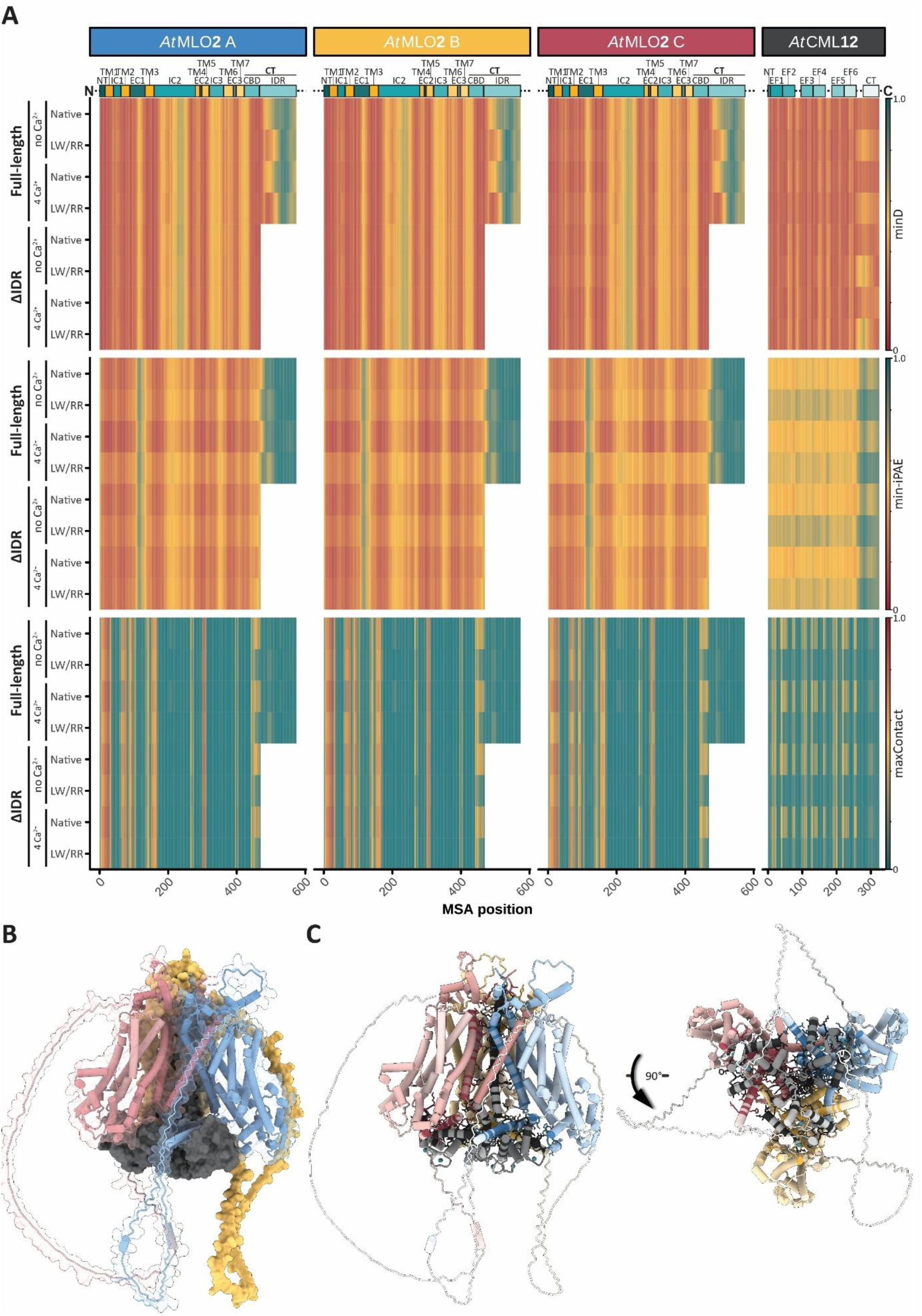
*At*CML12 binds to the CBDs and the pore surface of *At*MLO2 trimers in AF3 predictions. **A)** AF3 per-residue metrics highlight interfaces in *At*MLO2–*At*CML12 hetero-tetramer predictions. Heatmaps show the average minD, min-iPAE, and maxContact scores over 12 AF3 runs (one per seed), with five models per run. The color scale indicates the respective metric values as shown. The *At*MLO2 and *At*CML12 domain architectures are shown above the heatmaps. **NT**: extracellular/luminal N-terminal domain; **TM**: transmembrane domain; **IC**: intracellular loop; **AH**: amphipathic helix; **EC**: extracellular/luminal loop; **CT**: C-terminal domain; **CBD**: CAM-binding domain; **IDR**: intrinsically disordered region; **EF**: EF-hand-containing domain **B)** Surface representation of a representative AF3 prediction of the *At*MLO2–*At*CML12 hetero-tetramer (seed 1710, model 0). Surfaces of subunits A and C are shown semi-transparently to reveal the underlying ribbon representation. **C)** Same structure as in **B**, with interface residues highlighted according to the color scheme in **A** and side chains shown as sticks.

**Supplemental Figure 16:**
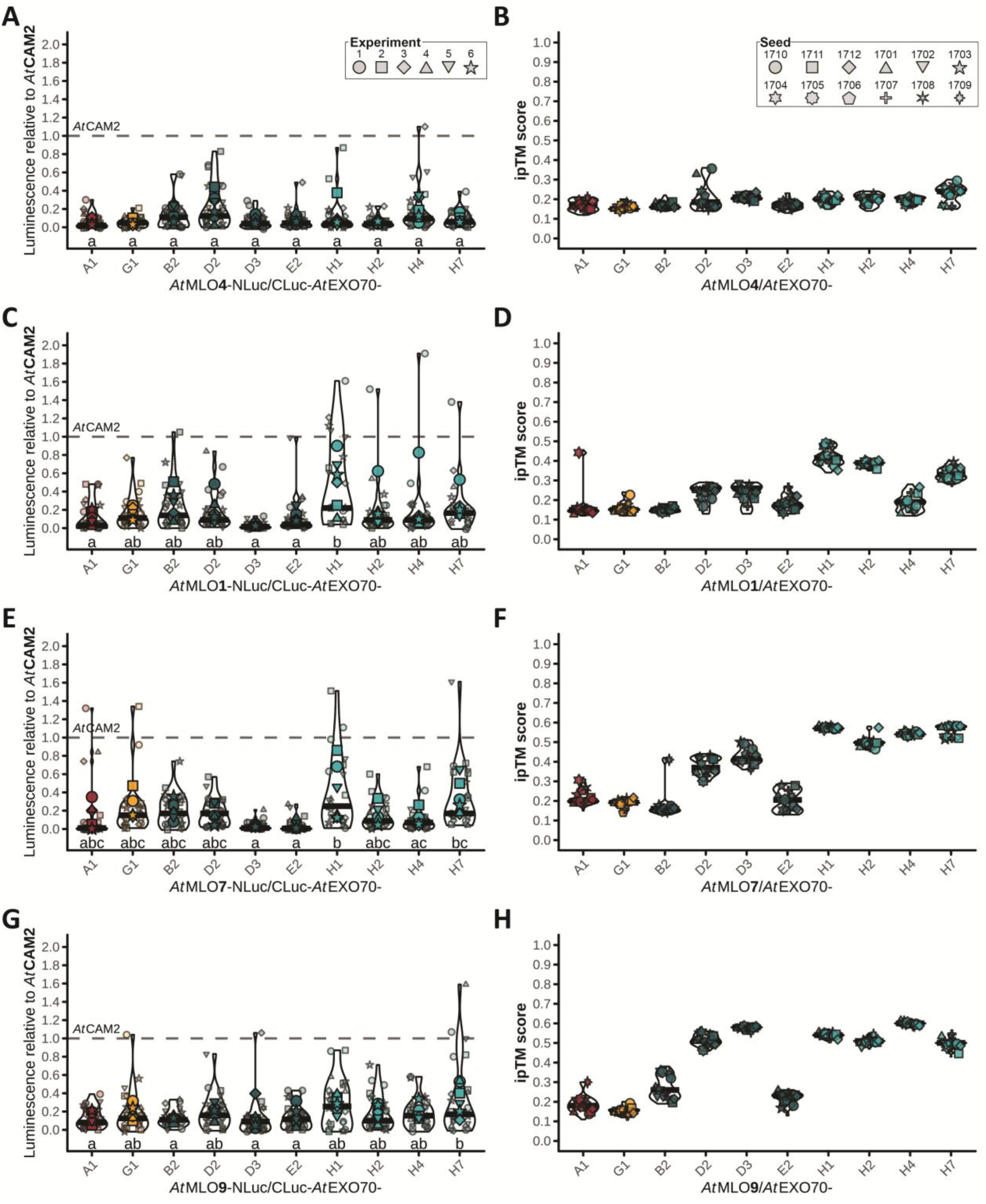
LCI-based and AF3-predicted interaction data of clade I, II, and III *At*MLO proteins with *At*EXO70 isoforms. Interactions between MLO and EXO70 proteins measured by LCI (**A**, **C**, **E**, **G**) and as predicted by AF3 ipTM scores (**B**, **D**, **F**, **H**). Violin plots show the probability distributions of luminescence signals relative to the interaction of MLO with *At*CAM2 or ipTM scores across 12 prediction runs (one per seed), with five AF3 models per run. For LCI, data points represent relative luminescence signals from six independent experiments (large points), with four replicates (small points) each. Statistically significant differences in LCI signals were determined by pairwise Student’s *t*-test (*α* = 0.01) and are indicated by different letters.

**Supplemental Figure 17:**
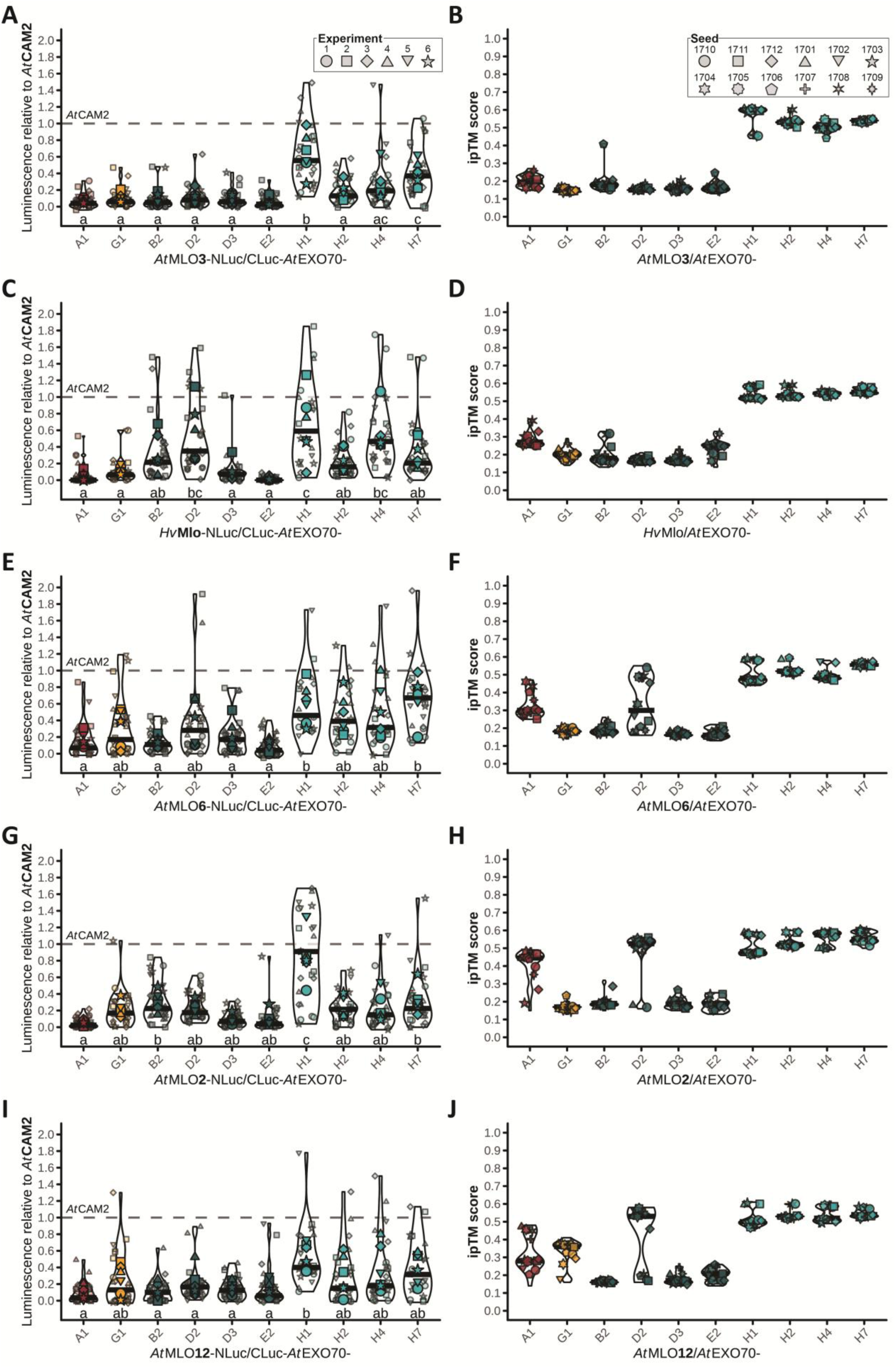
LCI-based and AF3-predicted interaction data for clade IV, V, and VI *At*MLO proteins with *At*EXO70 isoforms. Interactions between MLO and EXO70 proteins measured by LCI (**A**, **C**, **E**, **G**, **I**) and as predicted by AF3 ipTM scores (**B**, **D**, **F**, **H**, **J**). Violin plots show the probability distributions of luminescence signals relative to the interaction of MLO with *At*CAM2 or ipTM scores across 12 prediction runs (one per seed), with five AF3 models per run. For LCI, data points represent relative luminescence signals from six independent experiments (large points), with four replicates (small points) each. Statistically significant differences in LCI signals were determined by pairwise Student’s *t*-test (*α* = 0.01) and are indicated by different letters.

**Supplemental Figure 18:**
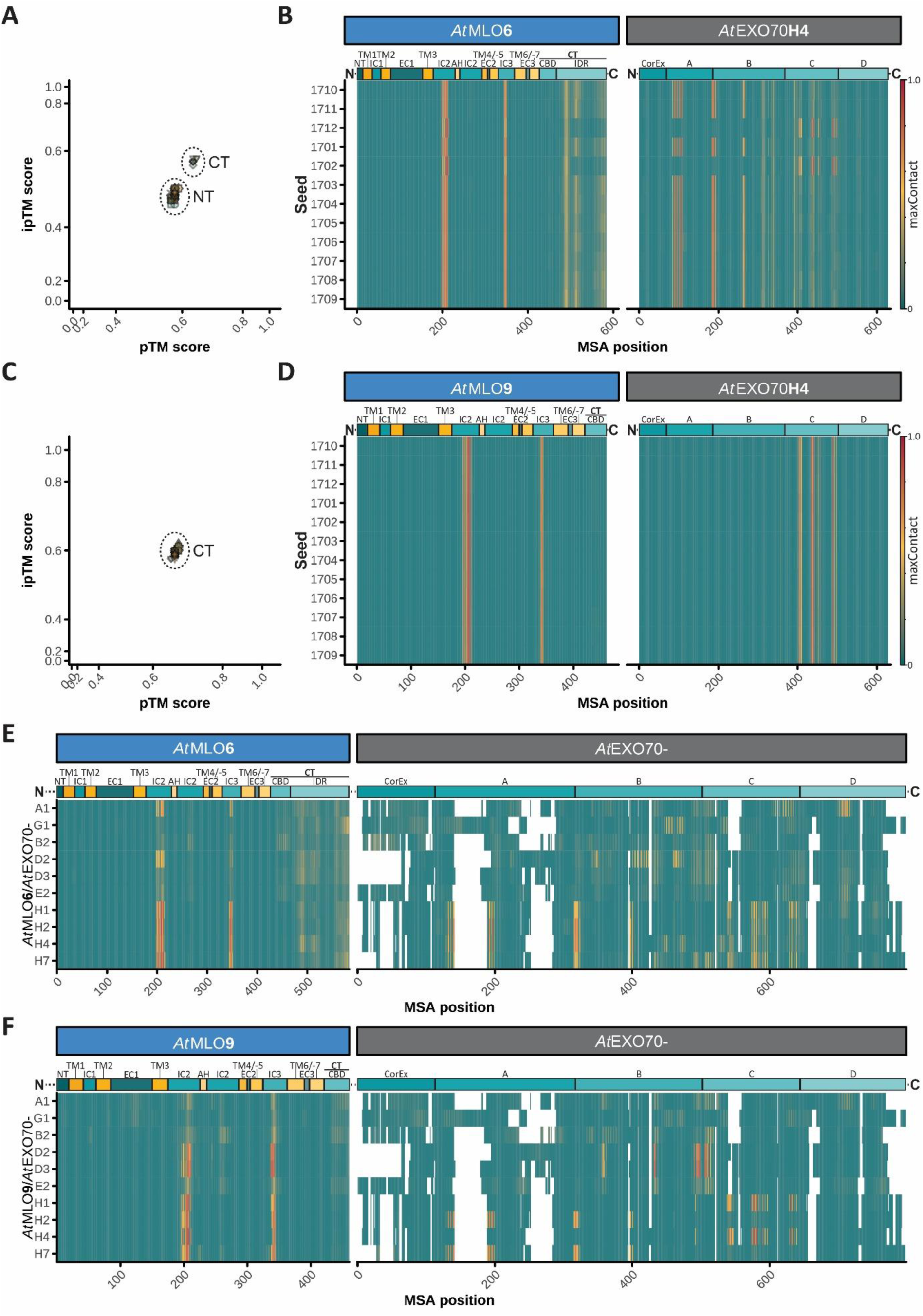
*At*EXO70H isoforms have two binding cavities for interacting *At*MLO proteins. **A, C)** Joint distribution of pTM and ipTM scores for *At*MLO6- (**A**) and *At*MLO9- (**C**) *At*EXO70H4 hetero-dimers. Clusters were labeled as N-terminal (NT) or C-terminal (CT) according to the binding cavity in *At*EXO70H4 targeted in the respective models. Axes were transformed using sigmoidal (logistic) functions to improve the visibility of data points. **B, D)** Average maxContact scores across the five models for the indicated seeds for *At*MLO6 (**B**) or *At*MLO9 (**D**) modeled with *At*EXO70H4. The color scales in **E** and **F** indicate maxContact values as shown. The *At*MLO and *At*EXO70H4 domain architectures are shown above the heatmaps. **NT**: extracellular/luminal N-terminal domain; **TM**: transmembrane domain; **IC**: intracellular loop; **AH**: amphipathic helix; **EC**: extracellular/luminal loop; **CT**: C-terminal domain; **CBD**: CAM-binding domain; **IDR**: intrinsically disordered region; **CorEx**: Core of exocyst domain; **A–D**: EXO70 domains following Dong *et al*. (2005). **E, F)** Average maxContact scores across 12 prediction runs (one per seed), with five AF3 models per run for *At*MLO6 (**E**) and *At*MLO9 (**F**) modeled with the indicated *At*EXO70 isoforms. The color scales indicate maxContact values as shown. The *At*MLO and *At*EXO70 consensus domain architectures are shown above the heatmaps.

**Supplemental Figure 19:**
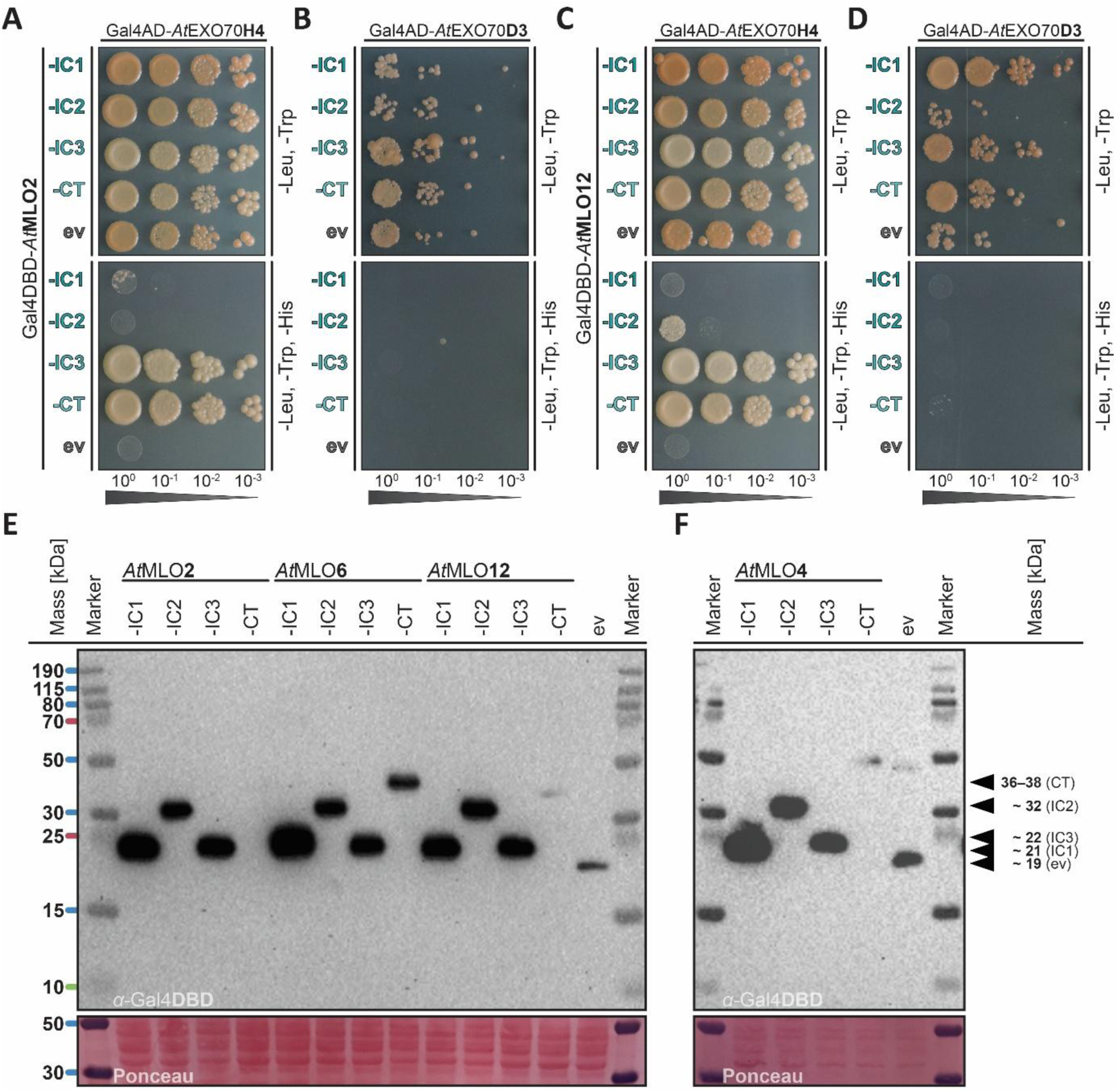
The IC3 domains and CTs of *At*MLO2 and -12 interact with *At*EXO70H4 but not -D3 in Y2H assays. **A–D)** Y2H assays probing interactions of intracellular domains from *At*MLO2 with *At*EXO70H4 (**A**) and *At*EXO70D3 (**B**) or *At*MLO12 with *At*EXO70H4 (**C**) and *At*EXO70D3 (**D**). Yeast cultures were spotted as 1:10 serial dilutions onto control medium lacking leucine and tryptophan (-Leu, -Trp) or selection medium additionally lacking histidine (-Leu, -Trp, -His). **AD**: activation domain; **DBD**: DNA-binding domain; **IC**: intracellular loop; **CT**: C-terminus; **ev**: empty vector. **E, F)** Immunoblot detection of the DBD of the yeast transcriptional regulator Gal4 fused to the intracellular domains of clade V *At*MLO2, -6, and -12 (**E**) or clade I *At*MLO4 (**F**). Molecular masses on the left correspond to the PageRuler Plus Prestained Protein Ladder (ThermoFisher Scientific, Waltham, MA). Masses given on the right represent the molecular mass of the recombinant proteins of interest. Ponceau S staining shows indicates the protein amount per SDS-PAGE pocket.

### Supplemental Tables

**Supplemental Table 1:** UniProt identifiers (IDs) of proteins investigated in this study.

| Name | UniProt ID | Organism |
| --- | --- | --- |
| **EF-hand proteins** |  |  |
| AtCAM2 | P0DH97 | *A. thaliana* |
| AtCML12 | P25071 | *A. thaliana* |
| **EXO70 proteins** |  |  |
| AtEXO70A1 | Q9LZD3 | *A. thaliana* |
| AtEXO70B2 | Q9LMJ4 | *A. thaliana* |
| AtEXO70D2 | Q9SYG5 | *A. thaliana* |
| AtEXO70D3 | Q9LJH9 | *A. thaliana* |
| AtEXO70E2 | Q9FNR3 | *A. thaliana* |
| AtEXO70G1 | Q7XYW9 | *A. thaliana* |
| AtEXO70H1 | Q8VY27 | *A. thaliana* |
| AtEXO70H2 | O80625 | *A. thaliana* |
| AtEXO70H4 | Q9SF51 | *A. thaliana* |
| AtEXO70H7 | Q9FN91 | *A. thaliana* |
| **MLO proteins** |  |  |
| AtMLO1 | O49621 | *A. thaliana* |
| AtMLO2 | Q9SXB6 | *A. thaliana* |
| AtMLO3 | Q94KB9 | *A. thaliana* |
| AtMLO4 | O23693 | *A. thaliana* |
| AtMLO5 | O22815 | *A. thaliana* |
| AtMLO6 | Q94KB7 | *A. thaliana* |
| AtMLO7 | O22752 | *A. thaliana* |
| AtMLO8 | O22757 | *A. thaliana* |
| AtMLO9 | Q94KB4 | *A. thaliana* |
| AtMLO10 | Q9FKY5 | *A. thaliana* |
| AtMLO11 | Q9FI00 | *A. thaliana* |
| AtMLO12 | O80961 | *A. thaliana* |
| AtMLO13 | Q94KB2 | *A. thaliana* |
| AtMLO14 | Q94KB1 | *A. thaliana* |
| AtMLO15 | O80580 | *A. thaliana* |
| HvMlo | P93766 | *H. vulgare* |
| MpMLO1 | A0A2R6XJW2 | *M. polymorpha* |
| MpMLO2 | A0A2R6W2I2 | *M. polymorpha* |
| MpMLO3 | A0A2R6X9C0 | *M. polymorpha* |
| MpMLO4 | A0A2R6X323 | *M. polymorpha* |

**Supplemental Table 2:**
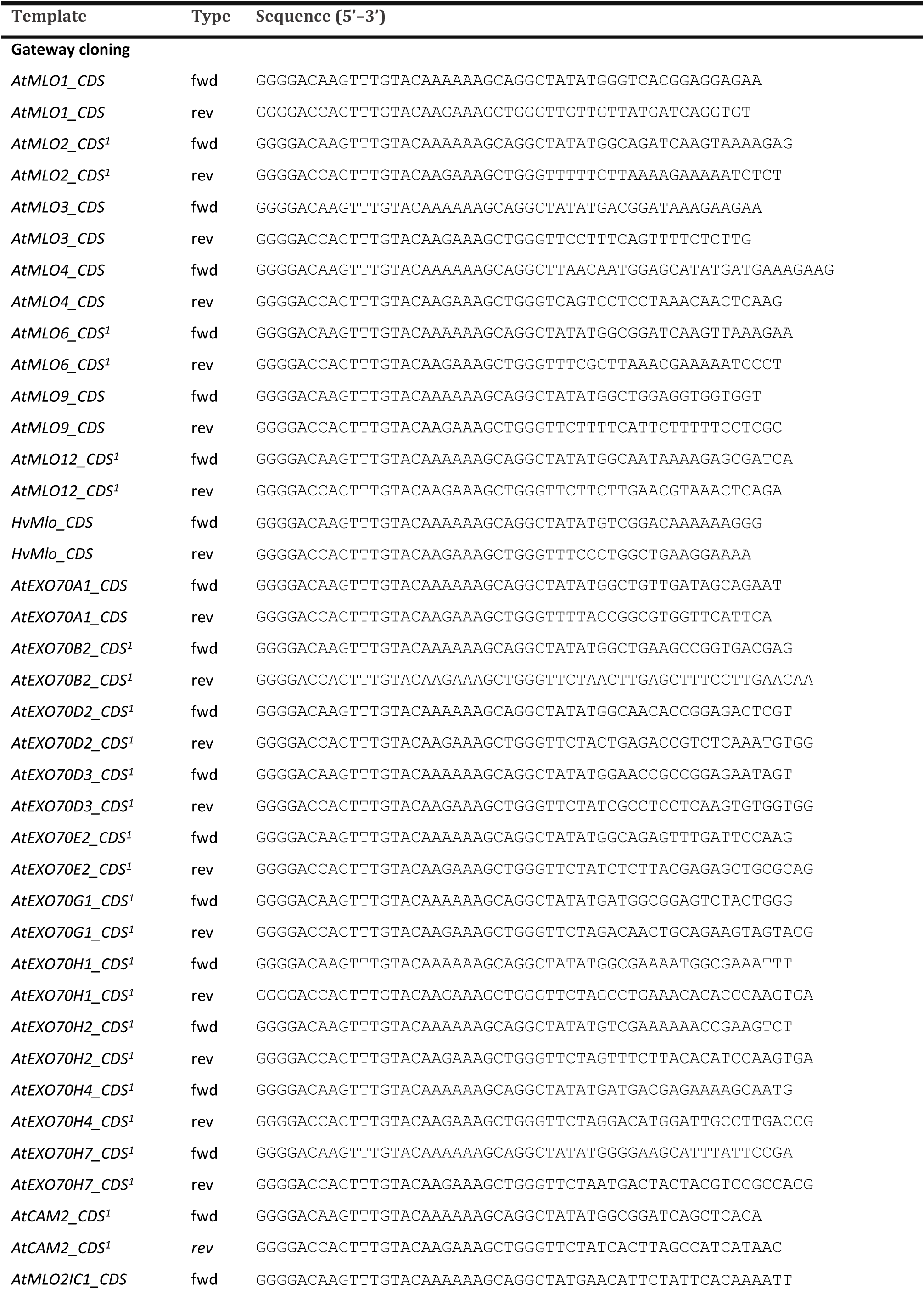
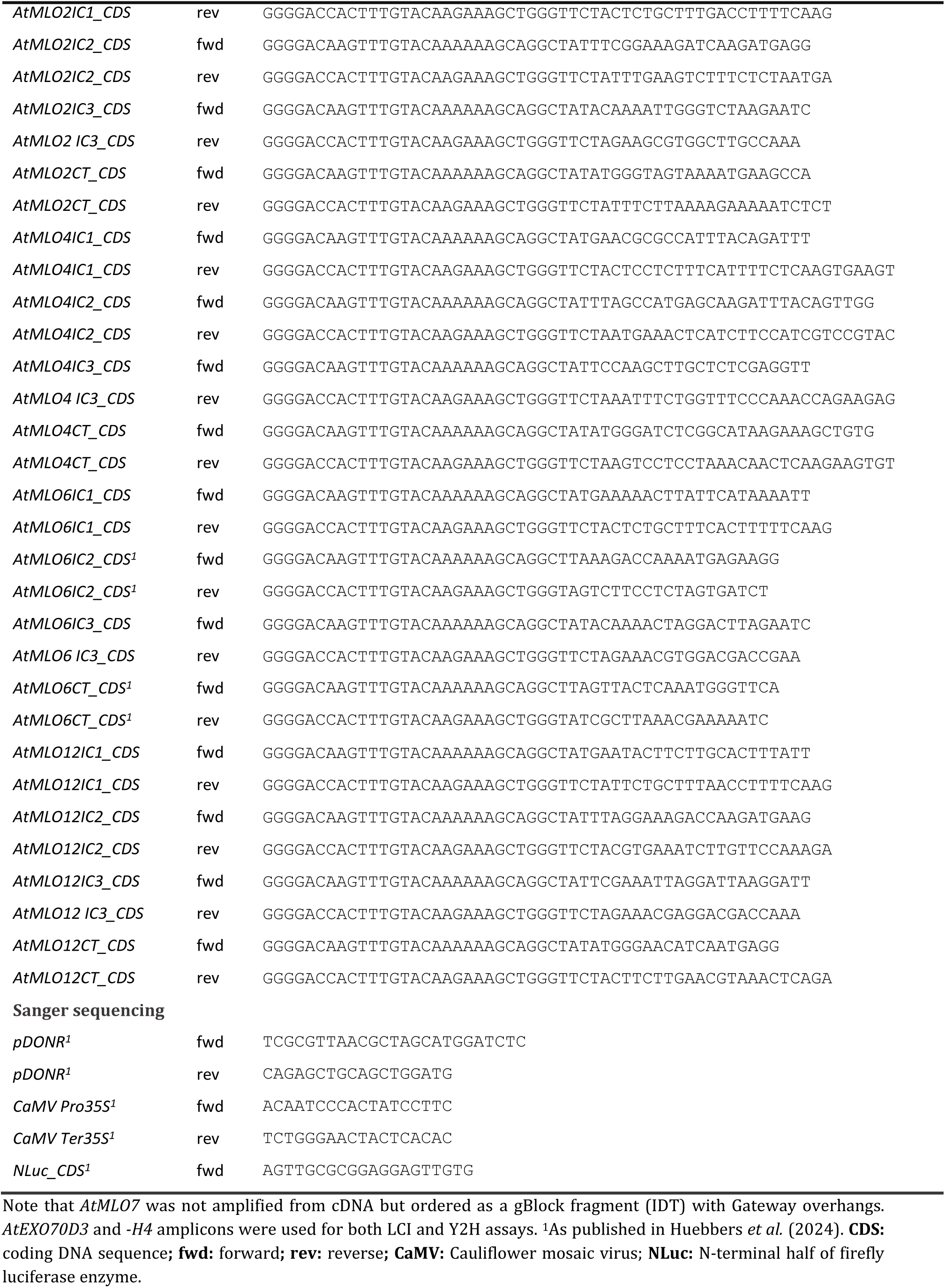
Oligonucleotides used in this study.

### Supplemental Movies

**Supplemental Movie 1: Transition between higher- and lower-confidence *Hv*MloΔIDR predictions.** Cytosolic (bottom) view of a ChimeraX *morph* trajectory interpolating between the higher-confidence model 1710_00 and the lower-confidence model 1705_02 from, obtained from AF3 predictions of *Hv*MloΔIDR trimers.

**Supplemental Movie 2: Ca²⁺ permeation and interactions within the *Hv*MloΔIDR pore.** Frames 4590–4650 (t = 91.800–93.000 ns) from a 100.000 ns all-atom MD simulation initiated from the *Hv*MloΔIDR trimer model 1705_02. Non-transparent cartoons of residues 1–160 of subunits A (blue) and B (yellow) are shown. Side chains of pore-lining anionic residues are displayed as sticks for all subunits (with subunit C in front). Ca²⁺ ions are shown in red; Cl⁻ counterions are omitted for clarity. The membrane is shown in cross-section.

### Supplemental Files

**Supplemental File 1: AF3 model confidences for predictions of *Hv*Mlo homo-oligomers and per-residue scores for *Hv*Mlo trimers.**

**Supplemental File 2: AF3 model and per-residue confidence scores for *Hv*MloΔIDR trimers.**

**Supplemental File 3: Composition of systems simulated in all-atom MD.**

**Supplemental File 4: AF3 model and per-residue confidence metrics for *At*MLO isoforms, *Mp*MLO isoforms, and *Hv*Mlo.**

**Supplemental File 5: AF3 model and mean per-residue confidence metrics for *At*MLO2-*At*CAM2 complexes.**

**Supplemental File 6: AF3 model and mean per-residue confidence metrics for *At*MLO2-*At*CML12 complexes.**

**Supplemental File 7: AF3 model metrics for *At*MLO-*At*EXO70 complexes.**

**Supplemental File 8: AF3 mean per-residue metrics for *At*MLO-*At*EXO70 complexes.**

**Supplemental File 9: MLO input sequenced for phylogenetic analysis.**

## Abbreviations

AF3: AlphaFold 3
CAM: Calmodulin
CBD: Calmodulin-binding domain
CT: Carboxy terminus
CML12: Calmodulin-like 12
CT: Carboxyl terminus
EXO70: Exocyst component of 70-kDa
FLIM: Fluorescence Lifetime Imaging
FRET: Fluorescence/Förster Resonance Energy Transfer
IC: Intracellular loop
IDR: Intrinsically disordered region
iPAE: Individual predicted aligned error
ipTM: Interface predicted template modeling score
LCI: Luciferase complementation imaging
MD: Molecular Dynamics
MLO: Mildew resistance Locus O
NT: Amino terminus
pTM: Predicted template modeling score
RMSD: Root mean square deviation
TM: Transmembrane domain
Y2H: Yeast two-hybrid

## References

Abraham, M.J., Murtola, T., Schulz, R., Páll, S., Smith, J.C., Hess, B., and Lindahl, E. (2015). GROMACS: High performance molecular simulations through multi-level parallelism from laptops to supercomputers. SoftwareX 1-2: 19–25.

Abramson, J., Adler, J., Dunger, J., Evans, R., Green, T., Pritzel, A., Ronneberger, O., Willmore, L., Ballard, A.J., et al.*, and* Jumper, J.M. (2024). Accurate structure prediction of biomolecular interactions with AlphaFold 3. Nature 630 (8016): 493–500.

Acharya, A., Nussberger, S., Wang, S., and Kleinekathöfer, U. (2025). Dynamic coupling between Tom22 motions and Tom40 pore dynamics modulates ion transport in the mitochondrial TOM complex. J. Chem. Inf. Model. 65 (22): 12475–12488.

Ali, S., Tyagi, A., Park, S., Varshney, R.K., and Bae, H. (2025). A molecular perspective on the role of FERONIA in root growth, nutrient uptake, stress sensing and microbiome assembly. J. Adv. Res. 75: 35–52.

Anderluh, A., Hofmaier, T., Klotzsch, E., Kudlacek, O., Stockner, T., Sitte, H.H., and Schütz, G.J. (2017). Direct PIP_2_ binding mediates stable oligomer formation of the serotonin transporter. Nat. Commun. 8: 14089.

Aryal, P., Jarerattanachat, V., Clausen, M.V., Schewe, M., McClenaghan, C., Argent, L., Conrad, L.J., Dong, Y.Y., Pike, A.C.W., et al.*, and* Tucker, S.J. (2017). Bilayer-mediated structural transitions control mechanosensitivity of the TREK-2 K2P channel. Structure 25 (5): 708–718.e2.

Aryal, P., Sansom, M.S.P., and Tucker, S.J. (2015). Hydrophobic gating in ion channels. J. Mol. Biol. 427 (1): 121–130.

Bavi, N., Cortes, D.M., Cox, C.D., Rohde, P.R., Liu, W., Deitmer, J.W., Bavi, O., Strop, P., Hill, A.P., et al., *and* Martinac, B. (2016). The role of MscL amphipathic N terminus indicates a blueprint for bilayer-mediated gating of mechanosensitive channels. Nat. Commun. 7: 11984.

Benjamini, Y., and Hochberg, Y. (1995). Controlling the false discovery rate: A practical and powerful approach to multiple testing. Journal of the Royal Statistical Society: Series B (Methodological) 57 (1): 289–300.

Bernetti, M., and Bussi, G. (2020). Pressure control using stochastic cell rescaling. J. Chem. Phys. 153 (11): 114107.

Bidzinski, P., Noir, S., Shahi, S., Reinstädler, A., Gratkowska, D.M., and Panstruga, R. (2014). Physiological characterization and genetic modifiers of aberrant root thigmomorphogenesis in mutants of *Arabidopsis thaliana MILDEW LOCUS O* genes. Plant Cell Environ. 37 (12): 2738–2753.

Binci, F., Guarneri, G., Somoza, S.C., Vascon, F., Capparotto, A., Di Nuzzo, E., Rago, G., Cendron, L., Navazio, L., and Giovannetti, M. (2025). A symbiotic MLO gene regulates root development via RALF34-triggered Ca^2+^ signalling in Lotus japonicus.

Bodenhofer, U., Bonatesta, E., Horejš-Kainrath, C., and Hochreiter, S. (2015). msa: an R package for multiple sequence alignment. Bioinformatics 31 (24): 3997–3999.

Bongartz, K. von, Sabelleck, B., Baquero Forero, A., Kuhn, H., Leissing, F., and Panstruga, R. (2023). Comprehensive comparative assessment of the *Arabidopsis thaliana* MLO2-CALMODULIN2 interaction by various *in vitro* and *in vivo* protein-protein interaction assays. Biochem. J. 480 (20): 1615–1638.

Braam, J., and Davis, R.W. (1990). Rain-, wind-, and touch-induced expression of calmodulin and calmodulin-related genes in Arabidopsis. Cell 60 (3): 357–364.

Bussi, G., Donadio, D., and Parrinello, M. (2007). Canonical sampling through velocity rescaling. J. Chem. Phys. 126 (1): 14101.

Callaway, E. (2024). Chemistry Nobel goes to developers of AlphaFold AI that predicts protein structures. Nature 634 (8034): 525–526.

Cao, M.-X., Li, S.-Z., and Li, H.-J. (2024). *Mp*MLO1 controls sperm discharge in liverwort. Nat. Plants.

Carpenter, E.P., Beis, K., Cameron, A.D., and Iwata, S. (2008). Overcoming the challenges of membrane protein crystallography. Current opinion in structural biology 18 (5): 581–586.

Chen, Z., Noir, S., Kwaaitaal, M., Hartmann, H.A., Wu, M.-J., Mudgil, Y., Sukumar, P., Muday, G., Panstruga, R., et al., and Jones, A.M. (2009). Two seven-transmembrane domain MILDEW RESISTANCE LOCUS O proteins cofunction in *Arabidopsis* root thigmomorphogenesis. Plant Cell 21 (7): 1972–1991.

Consonni, C., Humphry, M.E., Hartmann, H.A., Livaja, M., Durner, J., Westphal, L., Vogel, J., Lipka, V., Kemmerling, B., et al.*, and* Panstruga, R. (2006). Conserved requirement for a plant host cell protein in powdery mildew pathogenesis. Nat. Genet. 38 (6): 716–720.

Darden, T., York, D., and Pedersen, L. (1993). Particle mesh Ewald: An *N* ⋅log(*N*) method for Ewald sums in large systems. J. Chem. Phys. 98 (12): 10089–10092.

Demidchik, V., Shabala, S., Isayenkov, S., Cuin, T.A., and Pottosin, I. (2018). Calcium transport across plant membranes: mechanisms and functions. New Phytol. 220 (1): 49–69.

Devoto, A., Piffanelli, P., Nilsson, I., Wallin, E., Panstruga, R., Heijne, G. von, and Schulze-Lefert, P. (1999). Topology, subcellular localization, and sequence diversity of the Mlo family in plants. J. Biol. Chem. 274 (49): 34993–35004.

Dong, G., Hutagalung, A.H., Fu, C., Novick, P., and Reinisch, K.M. (2005). The structures of exocyst subunit Exo70p and the Exo84p C-terminal domains reveal a common motif. Nat. Struct. Mol. Biol. 12 (12): 1094–1100.

Dunbrack, R.L. (2025). Preprint: *Rēs ipSAE loquunt*: What’s wrong with AlphaFold’s *ipTM* score and how to fix it. bioRxiv.

Elliott, C., Müller, J., Miklis, M., Bhat, R.A., Schulze-Lefert, P., and Panstruga, R. (2005). Conserved extracellular cysteine residues and cytoplasmic loop-loop interplay are required for functionality of the heptahelical MLO protein. Biochem. J. 385 (Pt 1): 243–254.

Feng, Z.P., Hamid, J., Doering, C., Jarvis, S.E., Bosey, G.M., Bourinet, E., Snutch, T.P., and Zamponi, G.W. (2001). Amino acid residues outside of the pore region contribute to N-type calcium channel permeation. J. Biol. Chem. 276 (8): 5726–5730.

Ferreira, S.T., and Felice, F.G. de (2001). Protein dynamics, folding and misfolding: from basic physical chemistry to human conformational diseases. FEBS letters 498 (2-3): 129–134.

Gabriel, T.S., Hansen, U.-P., Urban, M., Drexler, N., Winterstein, T., Rauh, O., Thiel, G., Kast, S.M., and Schroeder, I. (2021). Asymmetric interplay between K^+^ and blocker and atomistic parameters from physiological experiments quantify K^+^ channel blocker release. Front. Physiol. 12: 737834.

Gao, Q., Wang, C., Xi, Y., Shao, Q., Hou, C., Li, L., and Luan, S. (2023). RALF signaling pathway activates MLO calcium channels to maintain pollen tube integrity. Cell. Res. 33 (1): 71–79.

Gao, Q., Wang, C., Xi, Y., Shao, Q., Li, L., and Luan, S. (2022). A receptor–channel trio conducts Ca^2+^ signalling for pollen tube reception. Nature 70: 809.

Gastwirth, J.L., Gel, Y.R., and Miao, W. (2009). The impact of levene’s test of equality of variances on statistical theory and practice. Statist. Sci. 24 (3).

Geilfus, C.-M. (2018). Chloride: from nutrient to toxicant. Plant Cell Physiol. 59 (5): 877–886.

Gietz, R.D., and Woods, R.A. (2002). Transformation of yeast by lithium acetate/single-stranded carrier DNA/polyethylene glycol method. Methods Enzymol. 350: 87–96.

Hamilton, E.S., Schlegel, A.M., and Haswell, E.S. (2015). United in diversity: mechanosensitive ion channels in plants. Annu. Rev. Plant Biol. 66: 113–137.

Hess, B., Bekker, H., Berendsen, H.J.C., and Fraaije, J.G.E.M. (1997). LINCS: A linear constraint solver for molecular simulations. J. Comput. Chem. 18 (12): 1463–1472.

Hollingsworth, S.A., and Dror, R.O. (2018). Molecular dynamics simulation for all. Neuron 99 (6): 1129–1143.

Huebbers, J.W., Caldarescu, G.A., Kubátová, Z., Sabol, P., Levecque, S.C.J., Kuhn, H., Kulich, I., Reinstädler, A., Büttgen, K., et al.*, and* Žárský, V. (2024). Interplay of EXO70 and MLO proteins modulates trichome cell wall composition and susceptibility to powdery mildew. Plant Cell 36 (4): 1007–1035.

James, P., Halladay, J., and Craig, E.A. (1996). Genomic libraries and a host strain designed for highly efficient two-hybrid selection in yeast. Genetics 144 (4): 1425–1436.

Jiang, W., Del Rosario, J.S., Botello-Smith, W., Zhao, S., Lin, Y.-C., Zhang, H., Lacroix, J., Rohacs, T., and Luo, Y.L. (2021). Crowding-induced opening of the mechanosensitive Piezo1 channel in silico. Communications biology 4 (1): 84.

Jo, S., Kim, T., Iyer, V.G., and Im, W. (2008). CHARMM-GUI: a web-based graphical user interface for CHARMM. J. Comput. Chem. 29 (11): 1859–1865.

Jones, D.S., Yuan, J., Smith, B.E., Willoughby, A.C., Kumimoto, E.L., and Kessler, S.A. (2017). MILDEW RESISTANCE LOCUS O function in pollen tube reception is linked to its oligomerization and subcellular distribution. Plant Physiol. 175 (1): 172–185.

Jørgensen, J.H. (1971). Comparison of induced mutant genes with spontaneous genes in barley conditioning resistance to powdery mildew. Mutation Breeding for Disease Resistance. Proceedings: 117–124.

Ju, Y., Yuan, J., Jones, D.S., Zhang, W., Staiger, C.J., and Kessler, S.A. (2021). Polarized NORTIA accumulation in response to pollen tube arrival at synergids promotes fertilization. Dev. Cell 56 (21): 2938–2951.e6.

Jumper, J., Evans, R., Pritzel, A., Green, T., Figurnov, M., Ronneberger, O., Tunyasuvunakool, K., Bates, R., Žídek, A., et al.*, and* Hassabis, D. (2021). Highly accurate protein structure prediction with AlphaFold. Nature 596 (7873): 583–589.

Kaur, A., Madhu, and Upadhyay, S.K. (2021). Mechanosensitive ion channels in plants. In Calcium Transport Elements in Plants (Elsevier), pp. 267–279.

Kefauver, J.M., Ward, A.B., and Patapoutian, A. (2020). Discoveries in structure and physiology of mechanically activated ion channels. Nature 587 (7835): 567–576.

Kessler, S.A., Shimosato-Asano, H., Keinath, N.F., Wuest, S.E., Ingram, G., Panstruga, R., and Grossniklaus, U. (2010). Conserved molecular components for pollen tube reception and fungal invasion. Science 330 (6006): 968–971.

Kim, M.C., Panstruga, R., Elliott, C., Müller, J., Devoto, A., Yoon, H.W., Park, H.C., Cho, M.J., and Schulze-Lefert, P. (2002). Calmodulin interacts with MLO protein to regulate defence against mildew in barley. Nature 416 (6879): 447–451.

Koncz, C., and Schell, J. (1986). The promoter of T_L_-DNA gene *5* controls the tissue-specific expression of chimaeric genes carried by a novel type of *Agrobacterium* binary vector. Mol. Gen. Genet. 204 (3): 383–396.

Kusch, S., and Panstruga, R. (2017). *mlo*-Based resistance: An apparently universal “weapon” to defeat powdery mildew disease. Mol. Plant Microbe Interact. 30 (3): 179–189.

Kusch, S., Pesch, L., and Panstruga, R. (2016). Comprehensive phylogenetic analysis sheds light on the diversity and origin of the MLO family of integral membrane proteins. Genome Biol. Evol. 8 (3): 878–895.

Kusch, S., Thiery, S., Reinstädler, A., Gruner, K., Zienkiewicz, K., Feussner, I., and Panstruga, R. (2019). Arabidopsis *mlo3* mutant plants exhibit spontaneous callose deposition and signs of early leaf senescence. Plant Mol. Biol. 101 (1-2): 21–40.

La Concepcion, J.C. de, Duverge, H., Kim, Y., Julian, J., Xu, H.D., Watt, M.N., Ikene, S.A., Bianchi, A., Grujic, N., et al., and Dagdas, Y. (2025). Electrostatic changes enabled the diversification of an exocyst subunit via protein complex escape. Nat. Plants.

Lam, A.K.M., Rheinberger, J., Paulino, C., and Dutzler, R. (2021). Gating the pore of the calcium-activated chloride channel TMEM16A. Nat. Commun. 12 (1): 785.

Larsen, S.T., Dannersø, J.K., Nielsen, C.J.F., Poulsen, L.R., Palmgren, M., and Nissen, P. (2024). Conserved N-terminal regulation of the ACA8 calcium pump with two calmodulin binding sites. J. Mol. Biol. 436 (20): 168747.

Lee, S., Tran, A., Allsopp, M., Lim, J.B., Hénin, J., and Klauda, J.B. (2014). CHARMM36 united atom chain model for lipids and surfactants. J. Phys. Chem. B 118 (2): 547–556.

Leicher, H., Schade, S., Huebbers, J.W., Munzert-Eberlein, K.S., Haljiti, G., Ludwig, C., Müller, M., Kinoshita, T., Engelsdorf, T., et al., and Stegmann, M. (2025). Preprint: Endogenous RALF peptide function is required for powdery mildew host colonization. bioRxiv.

Li, P., and Xiao, S. (2025). Diverse functions of plant MLO proteins: from mystery to elucidation. Annu. Rev. Phytopathol. 63 (1): 147–173.

Lidbrink, S.E., Howard, R.J., Haloi, N., and Lindahl, E. (2025). Resolving the conformational ensemble of a membrane protein by integrating small-angle scattering with AlphaFold. PLoS Comput. Biol. 21 (6): e1013187.

Lin, Y., Wallis, C., and Corry, B. (2026). BPS2026 – AlphaFold can be used to predict the oligomeric states of proteins. Biophys. J. 125 (4): 114a.

Lin, Y.-C., Guo, Y.R., Miyagi, A., Levring, J., MacKinnon, R., and Scheuring, S. (2019). Force-induced conformational changes in PIEZO1. Nature 573 (7773): 230–234.

Liu, X., Wang, J., and Sun, L. (2018). Structure of the hyperosmolality-gated calcium-permeable channel OSCA1.2. Nat. Commun. 9 (1): 5060.

Lomize, A.L., Todd, S.C., and Pogozheva, I.D. (2022). Spatial arrangement of proteins in planar and curved membranes by PPM 3.0. Protein Sci. 31 (1): 209–220.

Lomize, M.A., Pogozheva, I.D., Joo, H., Mosberg, H.I., and Lomize, A.L. (2012). OPM database and PPM web server: resources for positioning of proteins in membranes. Nucleic Acids Res. 40 (Database issue): D370–6.

Lopez-Mateos, D., Narang, K., and Yarov-Yarovoy, V. (2026). Exploring voltage-gated sodium channel conformations and protein-protein interactions using AlphaFold2. J. Gen. Physiol. 158 (2).

Luan, S. (2026). Calcium signaling in plants: Universal and unique paradigms. Cell 189 (4): 1001–1023.

Mann, H.B., and Whitney, D.R. (1947). On a test of whether one of two random variables is stochastically larger than the other. Ann. Math. Statist. 18 (1): 50–60.

Martzoukou, O., Karachaliou, M., Yalelis, V., Leung, J., Byrne, B., Amillis, S., and Diallinas, G. (2015). Oligomerization of the UapA purine transporter is critical for ER-exit, plasma membrane localization and turnover. J. Mol. Biol. 427 (16): 2679–2696.

Matsumura, M., Nomoto, M., Itaya, T., Aratani, Y., Iwamoto, M., Matsuura, T., Hayashi, Y., Mori, T., Skelly, M.J., et al., *and* Tada, Y. (2022). Mechanosensory trichome cells evoke a mechanical stimuli–induced immune response in *Arabidopsis thaliana*. Nat. Commun. 13 (1): 1216.

McCormack, E., and Braam, J. (2003). Calmodulins and related potential calcium sensors of Arabidopsis. New Phytol. 159 (3): 585–598.

Meng, J.-G., Liang, L., Jia, P.-F., Wang, Y.-C., Li, H.-J., and Yang, W.-C. (2020). Integration of ovular signals and exocytosis of a Ca^2+^ channel by MLOs in pollen tube guidance. Nat. Plants 6 (2): 143–153.

Miklis, M., Consonni, C., Bhat, R.A., Lipka, V., Schulze-Lefert, P., and Panstruga, R. (2007). Barley MLO modulates actin-dependent and actin-independent antifungal defense pathways at the cell periphery. Plant Physiol. 144 (2): 1132–1143.

Mölder, F., Jablonski, K.P., Letcher, B., Hall, M.B., van Dyken, P.C., Tomkins-Tinch, C.H., Sochat, V., Forster, J., Vieira, F.G., et al.*, and* Köster, J. (2021). Sustainable data analysis with Snakemake. F1000Res. 10: 33.

Moraes, I., Evans, G., Sanchez-Weatherby, J., Newstead, S., and Stewart, P.D.S. (2014). Membrane protein structure determination - the next generation. Biochim. Biophys. Acta 1838 (1 Pt A): 78–87.

Ngo, K., Yang, P.-C., Yarov-Yarovoy, V., Clancy, C.E., and Vorobyov, I. (2025). Harnessing AlphaFold to reveal hERG channel conformational state secrets. Elife 13.

Ogawa, S.T., Zhang, W., Staiger, C.J., and Kessler, S.A. (2025). MLO-mediated Ca^2+^ influx regulates root hair tip growth in Arabidopsis. New Phytol.

Opalski, K.S., Schultheiss, H., Kogel, K.-H., and Hückelhoven, R. (2005). The receptor-like MLO protein and the RAC/ROP family G-protein RACB modulate actin reorganization in barley attacked by the biotrophic powdery mildew fungus *Blumeria graminis* f.sp. *hordei*. Plant J. 41 (2): 291–303.

Park, S.-J., and Im, W. (2026). CHARMM-GUI Quick Bilayer: simple and intuitive one-stop membrane bilayer builder. J. Mol. Biol.: 169672.

Pettersen, E.F., Goddard, T.D., Huang, C.C., Meng, E.C., Couch, G.S., Croll, T.I., Morris, J.H., and Ferrin, T.E. (2021). UCSF ChimeraX: Structure visualization for researchers, educators, and developers. Protein Sci. 30 (1): 70–82.

Pinto-Anwandter, B.I. (2025). Voltage gating and 4-aminopyridine inhibition in the Shaker Kv channel revealed by a closed-state model. Biophys. J. 124 (15): 2500–2510.

Pogozheva, I.D., Armstrong, G.A., Kong, L., Hartnagel, T.J., Carpino, C.A., Gee, S.E., Picarello, D.M., Rubin, A.S., Lee, J., et al., *and* Im, W. (2022). Comparative molecular dynamics simulation studies of realistic eukaryotic, prokaryotic, and archaeal membranes. J. Chem. Inf. Model. 62 (4): 1036–1051.

Ponvert, N., and Johnson, M.A. (2024). Synergid cell calcium oscillations refine understanding of FERONIA/LORELEI signaling during interspecific hybridization. Plant Reprod. 37 (1): 57–68.

Raček, T., Vel’ký, D., Bučeková, G., Schindler, O., Hutařová Vařeková, I., Špačková, A., Bazgier, V., Berka, K., and Svobodová, R. (2025). MOLEonline: a web-based tool for analysing channels, tunnels, and pores (2025 update). Bioinformatics 41 (9).

Ryder, L.S., Sprakel, J., and Talbot, N.J. (2025). Mechanobiology of fungal invasion. Curr. Biol. 35 (11): R485–R490.

Schackert, F.K., Biedermann, J., Abdolvand, S., Minniberger, S., Song, C., Plested, A.J.R., Carloni, P., and Sun, H. (2023). Mechanism of calcium permeation in a glutamate receptor ion channel. J. Chem. Inf. Model. 63 (4): 1293–1300.

Schliep, K.P. (2011). phangorn: phylogenetic analysis in R. Bioinformatics 27 (4): 592–593.

Senior, A.W., Evans, R., Jumper, J., Kirkpatrick, J., Sifre, L., Green, T., Qin, C., Žídek, A., Nelson, A.W.R., et al.*, and* Hassabis, D. (2020). Improved protein structure prediction using potentials from deep learning. Nature 577 (7792): 706–710.

Shapiro, S.S., and Wilk, M.B. (1965). An analysis of variance test for normality (complete samples). Biometrika 52 (3-4): 591–611.

Sistrunk, M.L., Antosiewicz, D.M., Purugganan, M.M., and Braam, J. (1994). Arabidopsis TCH3 encodes a novel Ca^2+^ binding protein and shows environmentally induced and tissue-specific regulation. Plant Cell 6 (11): 1553–1565.

Smith, P.K., Krohn, R.I., Hermanson, G.T., Mallia, A.K., Gartner, F.H., Provenzano, M.D., Fujimoto, E.K., Goeke, N.M., Olson, B.J., et al.*, and* Klenk, D.C. (1985). Measurement of protein using bicinchoninic acid. Anal. Biochem. 150 (1): 76–85.

Student (1908). The probable error of a mean. Biometrika 6 (1): 1.

Sun, L., Qin, J., Wu, X., Zhang, J., and Zhang, J. (2022). TOUCH 3 and CALMODULIN 1/4/6 cooperate with calcium-dependent protein kinases to trigger calcium-dependent activation of CAM-BINDING PROTEIN 60-LIKE G and regulate fungal resistance in plants. Plant Cell 34 (10): 4088–4104.

Tao, E., and Corry, B. (2025). AlphaFold2 captures conformational transitions in the voltage-gated channel superfamily. Biophys. J. 124 (19): 3291–3303.

Torres, J., Pervushin, K., and Surya, W. (2024). Prediction of conformational states in a coronavirus channel using Alphafold-2 and DeepMSA2: Strengths and limitations. Comput. Struct. Biotechnol. J. 23: 3730–3740.

Tyagi, A., Ali, S., Park, S., and Bae, H. (2023). Deciphering the role of mechanosensitive channels in plant root biology: perception, signaling, and adaptive responses. Planta 258 (6): 105.

Vecchis, D. de, Beech, D.J., and Kalli, A.C. (2021). Molecular dynamics simulations of Piezo1 channel opening by increases in membrane tension. Biophys. J. 120 (8): 1510–1521.

Verkest, C., Schaefer, I., Nees, T.A., Wang, N., Jegelka, J.M., Taberner, F.J., and Lechner, S.G. (2022). Intrinsically disordered intracellular domains control key features of the mechanically-gated ion channel PIEZO2. Nat. Commun. 13 (1): 1365.

Voets, T., Prenen, J., Vriens, J., Watanabe, H., Janssens, A., Wissenbach, U., Bödding, M., Droogmans, G., and Nilius, B. (2002). Molecular determinants of permeation through the cation channel TRPV_4_. J. Biol. Chem. 277 (37): 33704–33710.

Wickham, H. (2016). ggplot2: Elegant graphics for data analysis.: Elegant graphics for data analysis (Switzerland: Springer).

Wickham, H., Averick, M., Bryan, J., Chang, W., McGowan, L., François, R., Grolemund, G., Hayes, A., Henry, L., et al.*, and* Yutani, H. (2019). Welcome to the Tidyverse. JOSS 4 (43): 1686.

Wu, E.L., Cheng, X., Jo, S., Rui, H., Song, K.C., Dávila-Contreras, E.M., Qi, Y., Lee, J., Monje-Galvan, V., et al., *and* Im, W. (2014). CHARMM-GUI Membrane Builder toward realistic biological membrane simulations. J. Comput. Chem. 35 (27): 1997–2004.

Xue, L., Yan, N., and Song, C. (2025). Deciphering Ca^2+^ permeation and valence selectivity in Ca_V_1: Molecular dynamics simulations reveal the three-ion knock-on mechanism. Proc. Natl. Acad. Sci. U S A 122 (22): e2424694122.

Yelshanskaya, M.V., Nadezhdin, K.D., Kurnikova, M.G., and Sobolevsky, A.I. (2021). Structure and function of the calcium-selective TRP channel TRPV6. J. Physiol. 599 (10): 2673–2697.

Yu, G., Smith, D.K., Zhu, H., Guan, Y., and Lam, T.T.-Y. (2017). ggtree an R package for visualization and annotation of phylogenetic trees with their covariates and other associated data. Methods Ecol Evol 8 (1): 28–36.

Yuan, J., Ogawa, S.T., Jones, D.S., Lucca, N., Ju, Y., and Kessler, S.A. (2025). Regulation of MILDEW RESISTANCE LOCUS-O trafficking by calmodulin-binding domains. J. Exp. Bot.

Yuan, R., Zhang, J., Kryshtafovych, A., Schaeffer, R.D., Zhou, J., Cong, Q., and Grishin, N.V. (2026). CASP16 protein monomer structure prediction assessment. Proteins 94 (1): 86–105.

Zhao, Q., Wu, K., Geng, J., Chi, S., Wang, Y., Zhi, P., Zhang, M., and Xiao, B. (2016). Ion permeation and mechanotransduction mechanisms of mechanosensitive piezo channels. Neuron 89 (6): 1248–1263.

Zhou, X., Li, B., Li, J., Sun, Y., Xie, R., Higashiyama, T., Xiao, S., Xin, G., and Su, S. (2025). A mechanosensitive ion channel controls touch-triggered stigma movement through manipulation of calcium signature in *Torenia*. Nat. Commun. 16 (1): 6296.

Zhu, L., Zhang, X.-Q., de Ye, and Chen, L.-Q. (2021). The Mildew resistance Locus O 4 interacts with CaM/CML and is involved in root gravity response. Int. J. Mol. Sci. 22 (11).

