## Supplementary figures and images for "AlphaFold 3 captures oligomeric states and interaction dynamics of MLO ion channels"

### Supplemental Movie 1

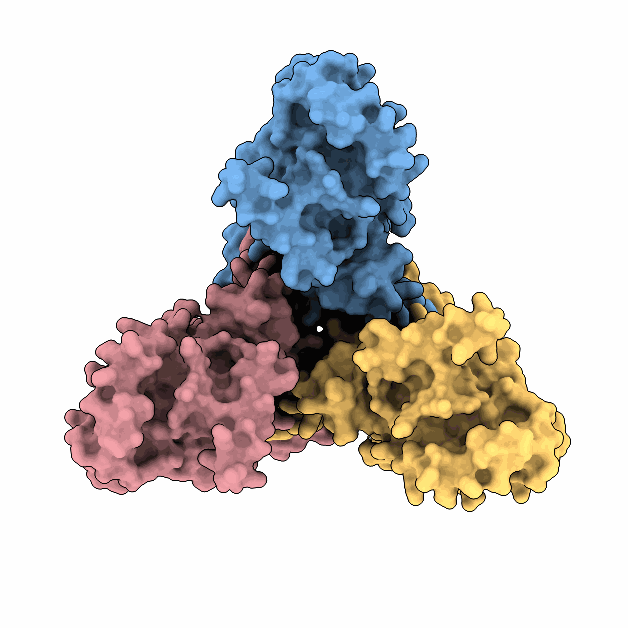

### Supplemental Movie 2

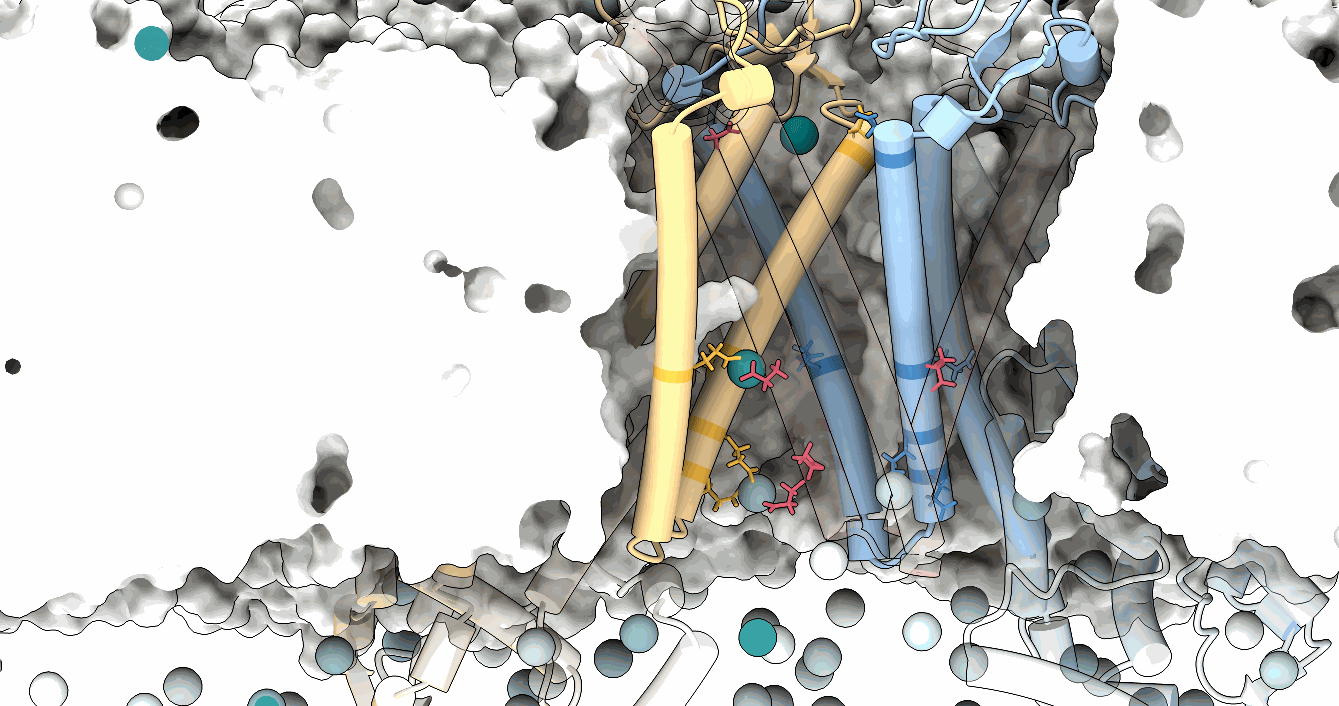
